# Evolutionary and functional dynamics of chimeric pseudogenes (φgenes)

**DOI:** 10.1101/2025.10.01.679709

**Authors:** Amruta R. Jadhav, Sushmita Sahoo, Yash L. Kuber, Avinash Varanwal, Khushi Tanwar, Shubhankar Palwankar, Gauri Chatrasal, Avinash M. Mali, Sharmila A. Bapat

## Abstract

Genome-associated changes during evolution followed by selection, facilitate development of new genes, gene families and even species. Pseudogenes (φgenes) generated during such genome rearrangements are often considered “dead on arrival” due to compromised expression arising from disruptive mutations or loss of regulatory elements. In an earlier study, we traced the expression of several φgenes, which challenges this notion. Here, we report segmental duplications at proximal genomic locations occasionally generate chimerism as a continuum of sequences from two or more genes, some of which involve intron fusions. These specific introns and flanking exons are likely to have co-evolved from ancestral parent genes. Notably, co-opting upstream regulatory sequences of 5′ parent gene may enable activation of chimeric φgenes, while modifications in 3′ sequences impart transcript stability. Few chimeric φgenes also harbor strong coding ORFs to be potentially translated into novel or truncated proteins retaining parental domains. Interestingly, chimeric φgenes are expressed only in human tissues, and display signatures of purifying selection. Examining potential functions associates *ANAPC1P2* with genotoxic stress responses, while *HYDIN2* may play a putative role in regulating cell cycle progression and neuronal differentiation. Taken together, our results suggest chimeric φgenes and their *de novo* functions may be relevant to speciation.

**Significance statement:** This study challenges the conventional paradigm that designates pseudogenes (φgenes) as “Dead-On-Arrival” sequences in the genome. Identification of chimeric φgenes formed by segmental duplications involving novel intron fusions highlights their role in evolutionary innovation. Notably, a subset of these chimeric φgenes demonstrates signatures of purifying selective pressure and exhibits exclusive, human-specific expression. More importantly, these φgenes may co-opt regulatory sequences, and contribute to novel gene functions such as genotoxic stress responses, and neuronal differentiation revealing putative mechanisms by which φgenes may foster evolutionary novelty.

**Graphical Abstract:** 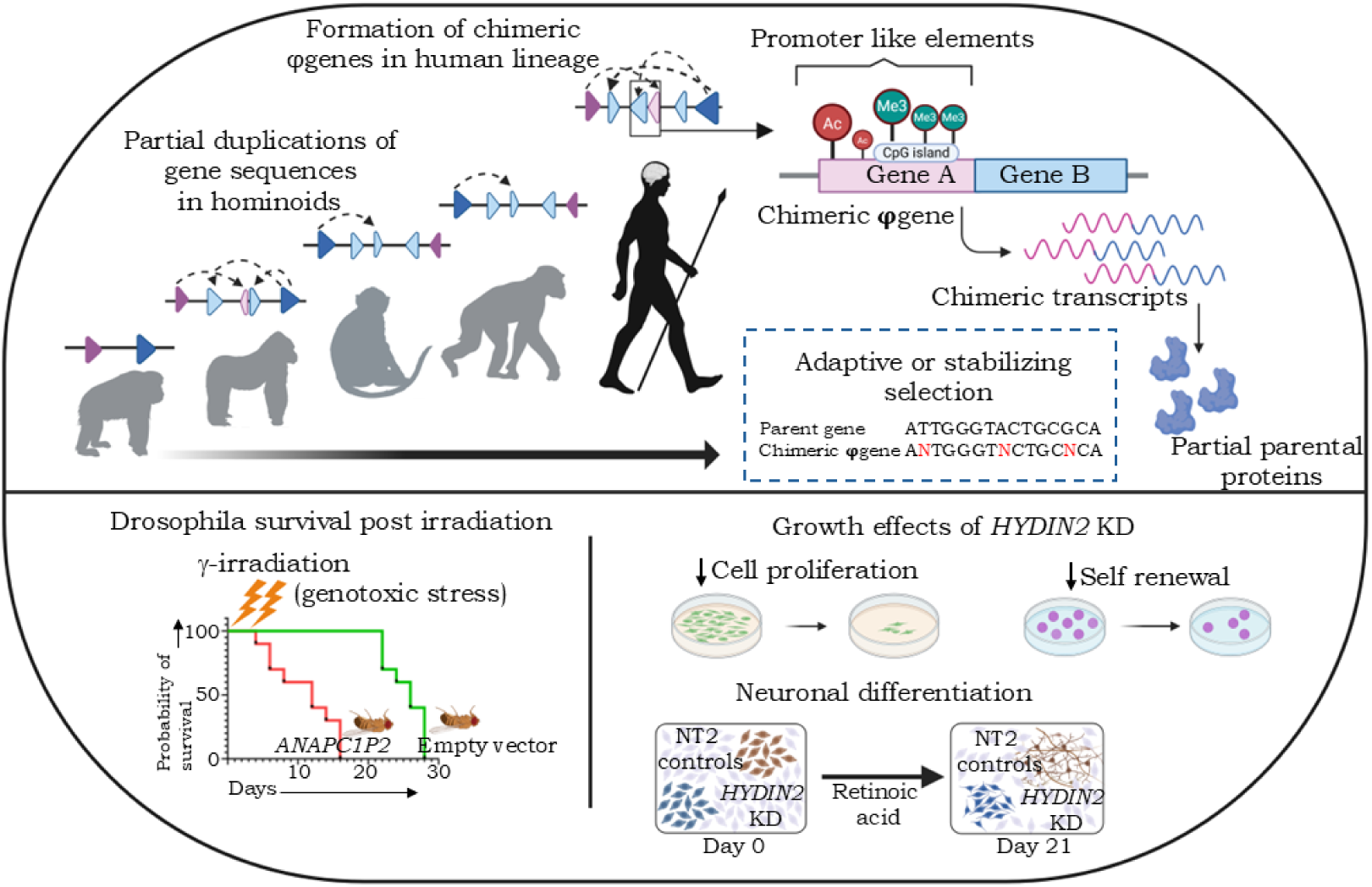

## Introduction

Dynamic turnover of genes through mutation, recombination, chromosomal fusion-fission, segmental rearrangements, transposon insertions, template switching, horizontal transfer, hybridization, and polyploidy is a cornerstone of evolution (1–5). Evolving genomes undergo frequent subtle and catastrophic events mediate stabilizing (favoring a phenotype), directional (broadening the phenotype spectrum to acquire new adaptive traits), or diversifying (promoting reproductive isolation and speciation) effects to drive natural selection (6,7). Positive selection shapes species through adaptation mediated by emergence of novel genes for neo-functionalization and gene families that display sub-functionalization (8–10). Conversely, negative / purifying selection removes non-beneficial sequences and is important in maintaining the reproductive isolation of newly emerged species and preventing them from merging back with the parental species after divergence. Thus, *de novo* sequences under purifying selection may persist under relaxed constraints, or evolve to gain functional potential, or confer selective advantages in speciation (11,12). Humans and great apes share highly similar genomes with divergence between lineages (∼2%) that despite being modest, impacts marked phenotypic and functional differences (13). Human-specific features include bipedalism, muscle composition, eye pigmentation, brain expansion, reduced hair, high sweat gland density, variable skin pigmentation balancing UV exposure with folate and vitamin D preservation, cognition and social behaviors, metabolic and immune adaptations, all of which reflect on adaptability to environmental challenges (14,15,16). Species divergence relies heavily on gene duplication, which in turn may involve pseudogenization. Numerous human-specific φgenes are identified, yet thought to be non-functional due to disabling mutations or loss of regulatory elements (17–19). Recent evidence however, indicates some transcribed φgenes to acquire novel cellular functions (20–22).

An earlier report from our lab identified several chimeric transcripts (CTs) representing sequences derived from two genes, a few of which matched sequences of reported φgenes (23). In the present study, following stringent selection we performed structural and functional annotation of a small subset of such chimeric φgenes. This revealed a novel feature of direct intron fusions and insertion of intervening sequences between exons of two parent genes, that suggests hitherto unknown mechanisms leading to chimerism. Further attributes including activation of ‘dormant’ chimeric φgenes by the 5′ parent promoter, CT stabilization by the 3′ partner, strikingly exclusive human-specific expression with possible relevance in either purifying or positive selection are also suggested. Our studies also established putative cellular roles for two chimeric φgenes. Collectively, these provide new insights into the emergence, regulatory dynamics, evolutionary significance, and functional contributions of chimeric φgenes.

## Results

### Evolution of ancestral parent gene sequences and intron fusions are associated with the emergence of chimeric φgenes that generate chimeric transcripts (CTs)

Our previous study had identified homology between 103 and 42 φgenes in NCBI BLASTn (https://www.ncbi.nlm.nih.gov/) and Ensembl BLAST (https://www.ensembl.org/) respectively (Methods). To ensure these represented authentic chimeras, we filtered out probable false positives, retaining only a small subset of 9 φgenes that display ∼ a 21 bp sequence of each parent gene flanking the fusion junction (Supplementary Figs. 1a, 1b; Supplementary Table 1). These φgenes are currently annotated as belonging to a single (usually 3′) parental gene and are assigned specific genomic locations in public databases. Such annotations often overlook the appended gene segment and failing to capture the chimeric nature of these transcripts. 9 of the purported φgenes identified as being chimeric were expressed at varying frequencies across tumors were selected for further studies (Fig.1a; Supplementary Figs.1a, 1b; Supplementary Table 1). Subsequent amplification and Sanger sequencing of their CTs across a panel of cell lines affirmed their chimeric nature and further revealed additional variations (Fig.1b; Supplementary Fig.1c). Towards understanding the emergence of chimeric φgenes, we studied the extent of sequence conservation between each chimera and its parent genes across 23 vertebrate species (Methods). Most parent genes are highly conserved and emerge early during vertebrate evolution; exceptions being *GATSL1* and *TRIM43* (Fig.1c; considered to emerge through *GATS* and *TRIM43B* duplication respectively; 24). Examination of the specific parent exon sequences flanking the fusionpoint in the 9 CTs further revealed that these emerged at a later branchpoint during evolution even in the highly conserved ancestral parent genes, and their CTs are expressed only in humans (Table 1; Supplementary Table 2). An example is of ancestral *GKAP1* and *KIF27* (highly conserved, branchpoint 0) that exhibit their specific exon sequences involved in chimerism at branchpoints 4 and 8 respectively, while chimeric *ENSG00000290924* (GKAP1-KIF27) sequences emerge in humans (branchpoint 13). In exploring the suggested altered splicing, significant splice site conservation was evident in RMND5A-ANAPC1, GATSL1-GTF2I, HIC2-PI4KA, GKAP1-KIF27 and RP4-565E6.1-HYDIN that led us to reject such a possibility (Supplementary Table 3).

**Table 1.**
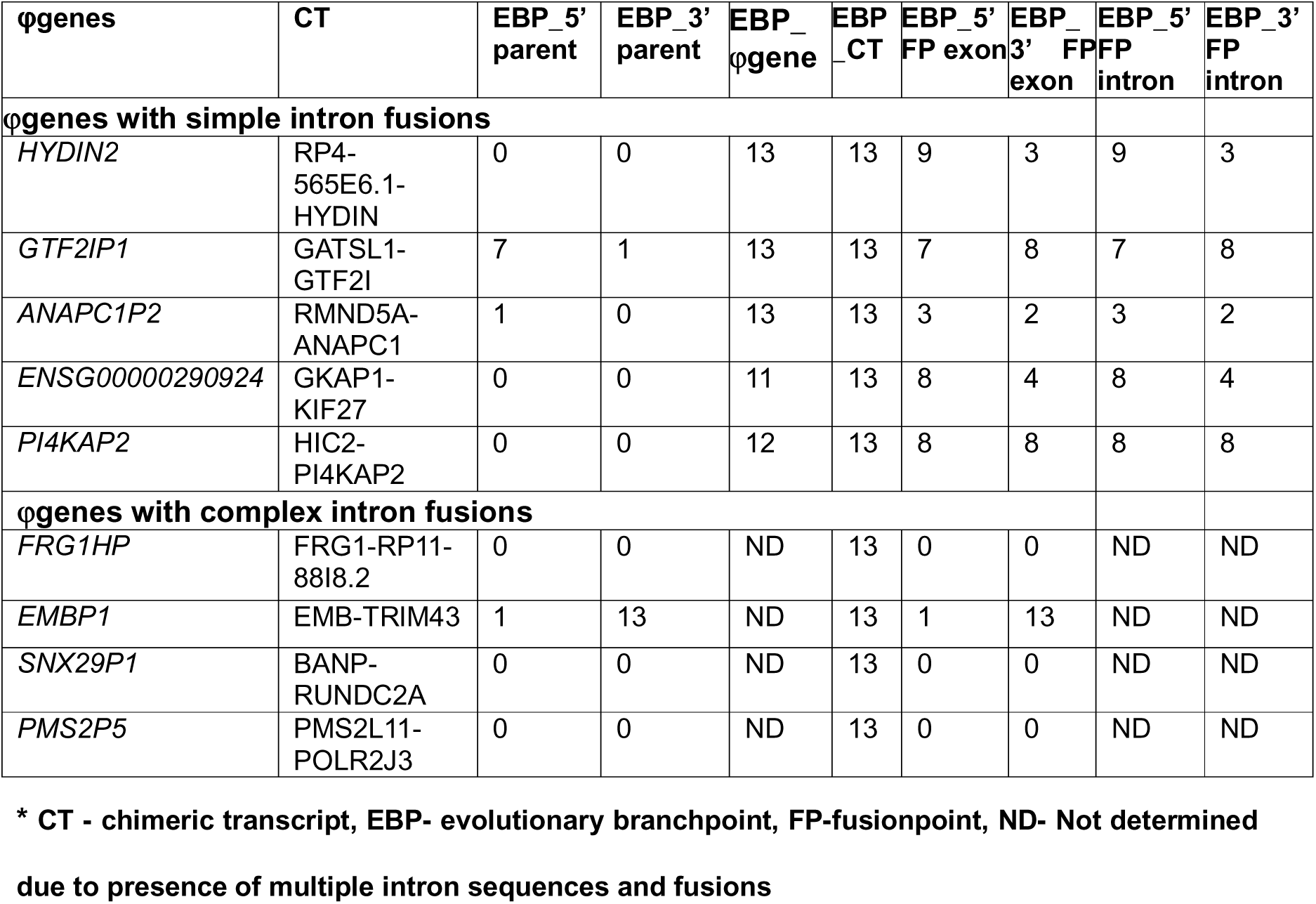
Conservation of parent genes, exons flanking the fusionpoint and CT sequences*.

Analyses of intron sequences positioned between parent exons flanking the fusionpoint in each φgene revealed two unique rearrangements (Fig.1d-i). The first of these involves generation of a chimeric intron by simple fusion of parent intron sequences (*HYDIN2, GTF2IP4*, *ANAPC1P2*, *ENSG00000290924*, *PI4KAP2*; Fig.1d-ii), which was validated through amplification and Sanger sequencing (Supplementary Fig.1d). The second type of intron sequence rearrangements involved complex, mosaic rearrangements including either insertion of intergenic sequences between parent introns (*EMBP1*, *SNX29P1*); *de novo* inclusion of unrelated gene sequences between parent introns (*FRG1HP*), or intronization of parental exons along with insertion of intergenic sequences (*PMS2P5;* Fig.1d-iii).

Studying these sequences further, we also realized that the specific parent introns undergoing fusion coevolved and emerged at the same branchpoint as did the flanking exons (Fig.1e; Table 2; Supplementary Table 2). We thus postulated that emergence of specific exon-intron sequence continuum involved in chimerism, intron fragmentation/breaks and fusion may occur as independent events over evolutionary time following ancestral gene formation. For instance, specific continuum of *RP4-565E6.1* and *HYDIN* emerged at branchpoints 9 and 3 respectively, while intron fusion and chimeric *ENSG00000290924* emergence is noted in the human genome at branchpoint 13. A similar case is of early emergence of the specific continuum in parental genes (branchpoints 3 and 2) with chimeric *ANAPC1P2* emerged in the human genome. A variation in this theme of human-specific chimerism is that chimeric introns of *ENSG00000290924* and *PI4KAP2* were traced earlier in the Gorilla genome, however transcripts have so far being identified only in the human lineage. Conclusively, coevolution of the specific introns and exons flanking the fusionpoint, and hitherto unidentified intron fusions are likely to be central in generation of chimeric φgenes.

**Fig.1.**
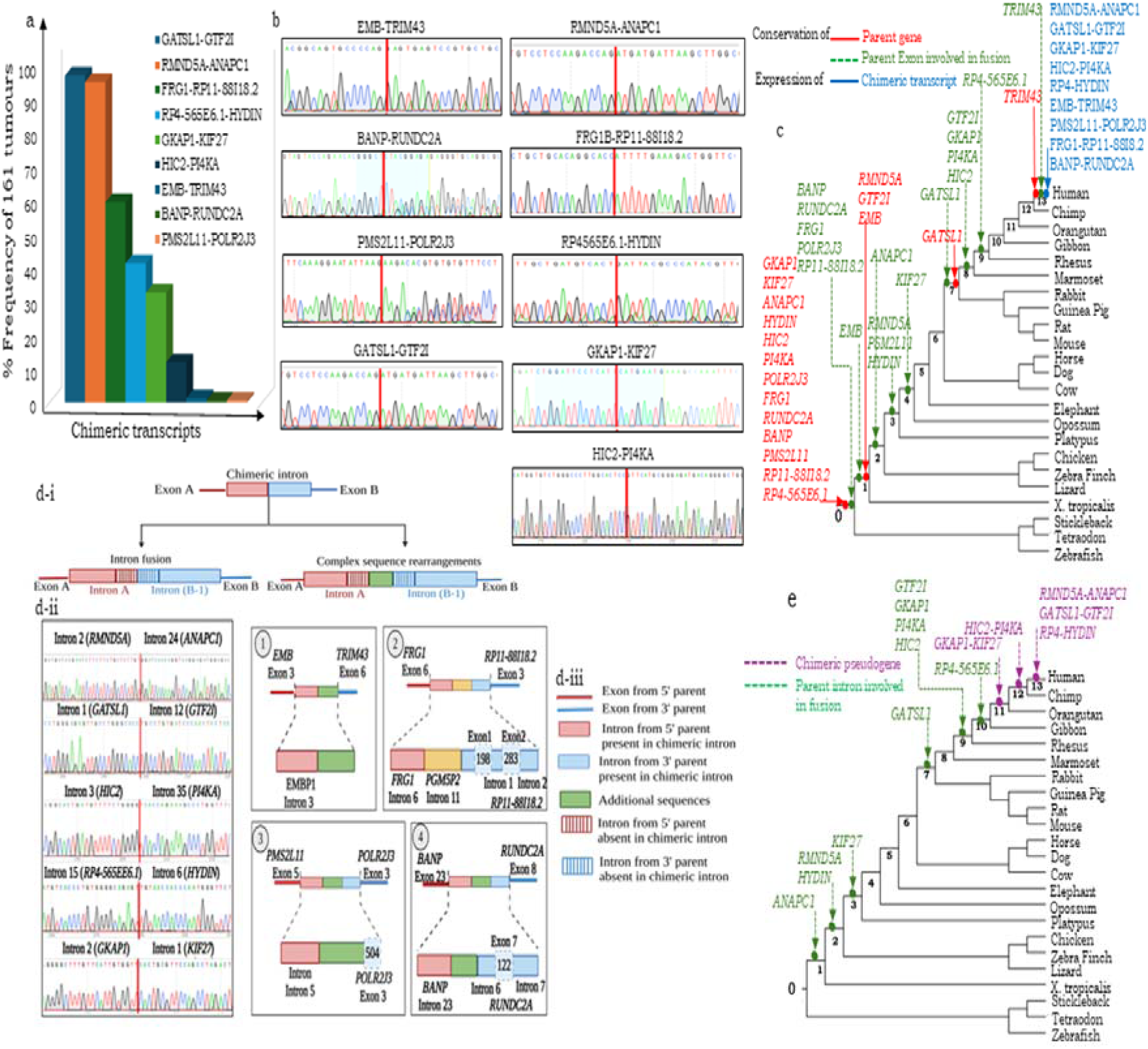
a. Distribution of candidate chimeric transcripts (CTs) across 161 tumour RNA-sequencing datasets in TCGA; b. PCR-based validation of 9 candidate CTs in HGSC cell lines and Sanger sequencing of amplicons flanking the fusionpoint; c. Phylogram representing the evolutionary conservation of parent gene and CT expression; d-i. Schematic representation of the two types of intron fusions resolved in φgenes, d-ii. Validation of simple intron fusions through genomic PCR and Sanger sequencing, d-iii. Schematic depicting complex intron fusions in 4 φgenes (dotted lines encompass the chimeric introns); e. Evolutionary conservation of sequences involved in intron fusions.

**Table 2.** Number of Duplications of parent genes observed in Hominoids.

| Organism | Number of Duplication of parent genes |  |  |  |  |  |  |  |  |  |
| --- | --- | --- | --- | --- | --- | --- | --- | --- | --- | --- |
|  | *RMND5A* | *ANAPC1* | *GKAP1* | *KIF27* | *RP4* | *HYDIN* | *GATSL1* | *GTF2I* | *HIC2* | *PI4KA* |
| Gorilla | 1 | 2 | 2 | 3 | 1 | 2 | 1 | 6 | 1 | 2 |
| Bonobo | 0 | 3 | 1 | 2 | 6 | 0 | 1 | 2 | 1 | 2 |
| Chimpanzee | 0 | 2 | 1 | 2 | 3 | 0 | 1 | 8 | 1 | 2 |
| Human | 1 | 9 | 1 | 2 | 3 | 4 | 2 | 8 | 2 | 2 |

### Chimeric **φ**genes emerge from segmental duplication events

Gene duplication and subsequent divergence during speciation is known to enhance the genome by creating gene families, each member of which can further acquire tissue-specific expression and functions (25). To evaluate the 5 chimeric φgenes generated through simple intron fusions as a part of such processes, we first listed and examined all parental φgenes and specific genomic loci. All φgenes thus examined harbour partial sequence duplications of parent genes with some or no overlap between different variants, however all including chimeric derivatives are at present reported as single parent φgenes (usually 3’ parent; Figs.2a,2b; Table 2; Supplementary Fig.2). Further study of duplication patterns of the 5 chimeric φgenes suggests that these may have emerged after the evolutionary split between other hominoids and humans (Fig.2c). Moreover, while most φgenes tend to cluster in tandem near the parent some *GTF2I*, *HYDIN* φgenes also transpose to other chromosomes. While *GTF2I* exhibits extensive duplication across hominids ranging from orangutan to humans, its chimeric φgenes located close to the parental genomic locus are exclusive to humans (Supplementary Fig.3a). Three unannotated duplications of RP4-565E6.1 were identified through sequence similarities, while four annotated φgenes of HYDIN (*HYDINP1, ENSG00000276298, ENSG00000259798, HYDIN2 /* RP4-565E6.1-HYDIN) exist, of which *HYDIN2* is exclusive to humans; chimpanzees and bonobo display six and three duplications of *RP4-565E6.1* respectively and none of *HYDIN* (Supplementary Fig.3b). No duplications of *RMND5A* are reported in hominoids, while several *ANAPC1* duplications are reported in Gorilla, chimpanzee, bonobo, and humans (*ANAPC1P1*-*ANAPC1P6* being annotated, while *ANAPC1P7-ANAPC1P9* were identified through sequence homology; Supplementary Fig.3c). Chimeric *KIF27* φgenes have an interesting trajectory across great apes, one emerged in orangutans and is subsequently conserved, gorillas harbour two duplications; one was lost in evolution (due to genomic rationalization or lack of selective advantage), the second persisted and is expressed in humans (*ENSG00000290924*; Fig.2d). The *PI4KA* locus harbouring several duplications is similarly conserved across hominoids; presence of one chimeric *PI4KA* φgene in gorillas and two in humans (*PI4KAP1*, *PI4KAP2*) suggests continuing gene duplication and/or rearrangement during speciation (Fig.2e). This points to a critical distinction between presence of chimeric sequences in the genome and their transcriptional activation in a species-specific manner, where orthologous genomic sequences in a particular lineage remain transcriptionally silent or non-functional, while becoming biologically active and potentially adaptive in closely related species (26). A unique feature of *RMND5A* and *ANAPC1* φgenes is their location on human chromosome 2, which itself is believed to emerge from the fusion of ancestral primate chromosomes 2a and 2b; with *ANAPC1* and its φgenes being localized at the 2q13–2q14.1 region wherein dynamic restructuring of the two of four ancestral telomeric sequences is believed to have occurred (Supplementary Figs.3d).

**Fig.2.**
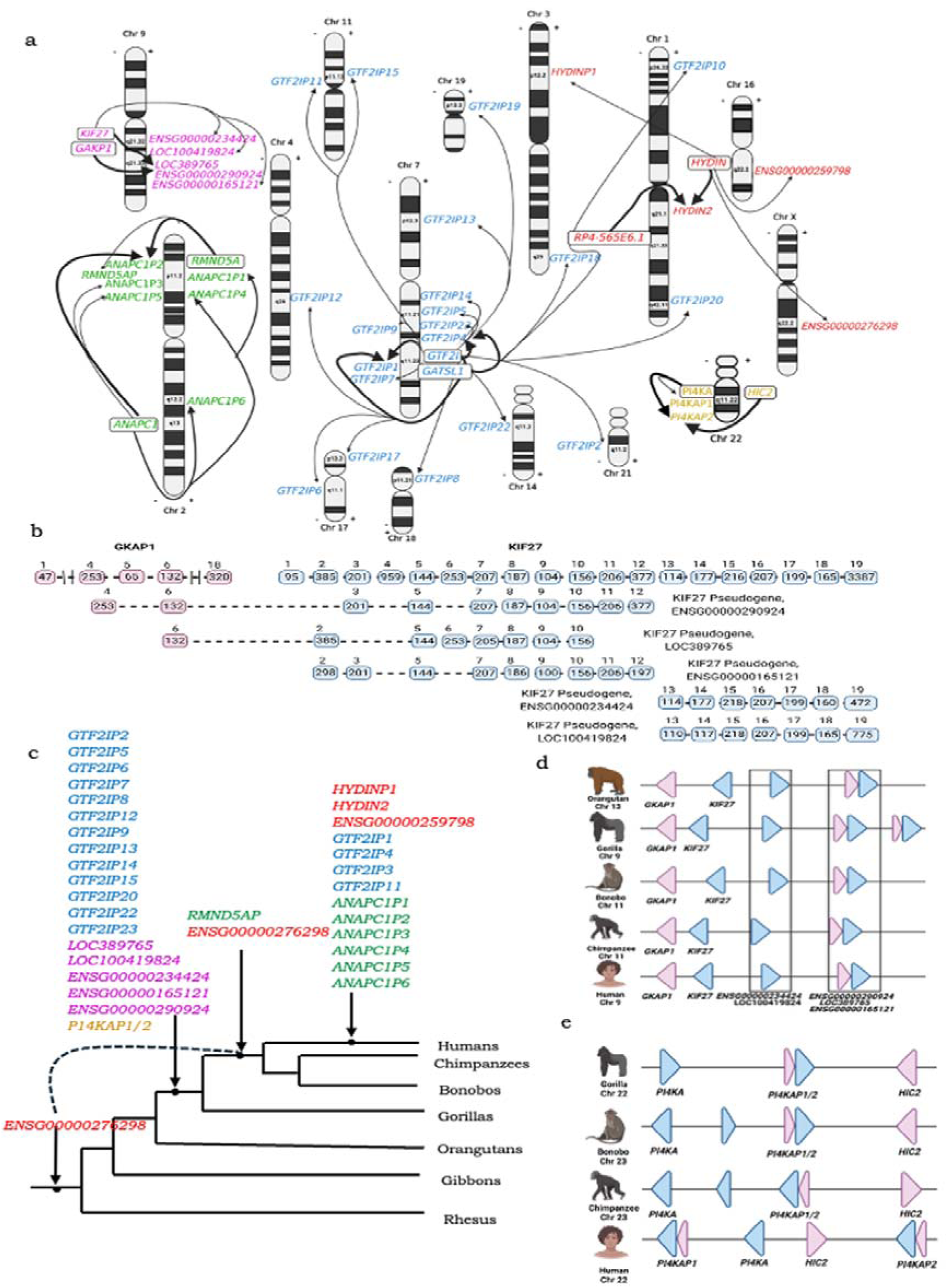
Segmental duplication events in the formation of chimeric φgenes involving simple intron fusions. a. Chromosome mapping of parental genes and their reported φgenes; b. Schematic indicating partial sequence homology of *KIF27* φgenes with parental genes; c. Conservation of these φgenes across various species; d. Schematic representation of conservation of segmental duplication of *GKAP-KIF27*; e. Schematic representation of conservation of segmental duplication of *HIC2-PI4KA*.

### Chimeric **φ**genes undergo evolutionary selection, with discrete regulatory elements contributing to their expression and stabilization

Given their human-specific expression, we assessed the evolutionary significance of parental *vs.* φgenes. Chimeric φgenes displayed a significantly higher GC content compared to their single-parent counterparts aligning them with prior reports of functional conservation in GC-rich φgenes (Fig. 3a; 27). Further calculating the Ka/Ks ratios (ω) for their coding regions to associate the expression of chimeric φgenes to be evolving under selective adaptation, or acquiring lineage-specific functions, indicated four of five chimeric φgenes (*ANAPC1P2, HYDIN2, PI4KAP2, ENSG00000290924*) showed evidence of purifying selection, while *GTF2IP4* is indicated to be under positive selection, (Fig.□3b). Together, these data suggest a non-neutral evolution of chimeric φgenes with sustained transcriptional activity of the former group likely to be coupled with selective constraints that preserve coding potential and functional relevance while *GTF2IP4* may be associated with possible neofunctionalization.

Further deep dive into the transcription of chimeric φgenes by extracting their profiles across a wide range of human tissues in the Genotype-Tissue Expression database [GTEx; https://services.healthtech.dtu.dk/services/Promoter-2.0/] revealed a discrete expression across multiple human tissue types as compared with parental genes (Supplementary Fig.□4a). Since chimeric φgenes are assigned distinct genomic loci, this suggests them to be regulated independently of the parent loci, or specifically being transcribed in contexts where parental genes cannot be expressed. Examining the 5’ upstream regulatory elements of each φgene in the UCSC Genome Browser (https://genome.ucsc.edu/) indeed identified the association of promoter-like elements CpG islands, FANTOM5 reads, ReMap density and transcription start sites (TSS) and active chromatin marks (H3K27Ac; H3K4me1, H3K4me3), some of which were positioned within or in the proximity of the first exon of the 5’ parent gene (Figs.3c, 3d; Supplementary Fig.4b). Interestingly such features were enriched in chimeric over single-parent φgenes (Supplementary Fig.4b), although activation of few single-parent φgenes relies on shared proximal promoter elements of adjacent genes; for example, *ANAPC1P1* and *ANAPC1P4* share *CD8B* and *CYTOR* promoters respectively (Supplementary Figs.4c). It is also quite likely that SDs creating φgenes include parental regulatory elements in continuum with gene sequences over those involving partial duplications; for example, the first exon of *GTF2IP20* displays promoter activity, while partially duplicated *GTF2IP10, GTF2IP2, GTF2IP3* lack regulatory regions (Supplementary Figs.4d). Complementing the enrichment of 5’ regulatory regions in chimeric φgenes is increased frequency and distribution of m6A methylation marks across their sequences over those in single-parent φgenes (Fig.3e; Supplementary Figs.4e). Further integrative analysis of chromatin states and transcriptional activity demonstrated that chimeric φgenes and their corresponding parental genes frequently co-cluster based on shared regulatory elements and expression patterns (Fig.□3f). Thereby, distinct regulatory signatures may contribute to enhanced stability and expression of chimeric φgenes that are under evolutionary constraint, and are likely evolve and become potentially functional.

**Fig.3.**
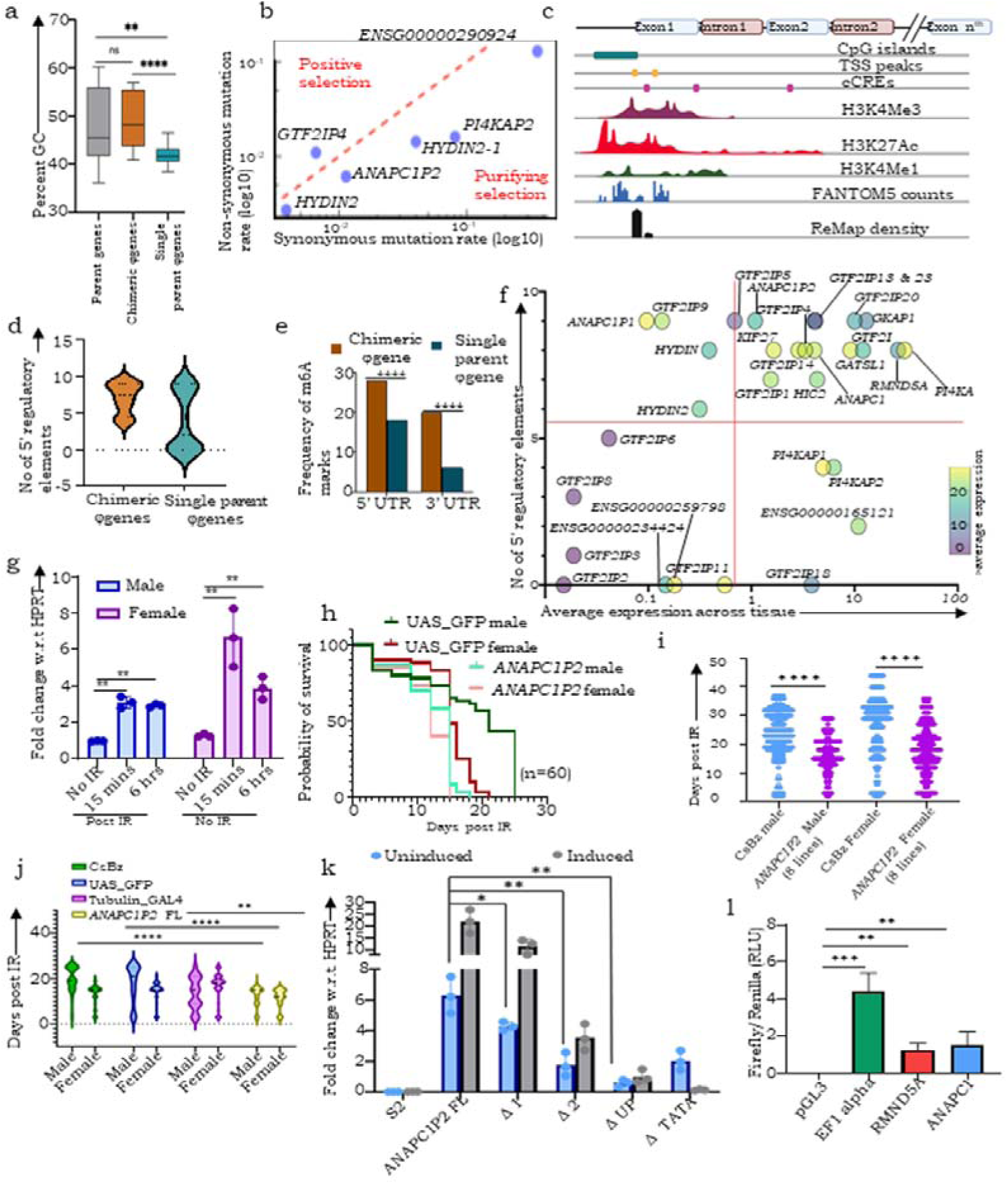
Promoter sequences in their neighbourhood drive transcription of φgenes generated through segmental duplication. a. Quantification of %GC content of parent, chimeric and single-parent φgene transcripts; b. Ka/Ks ratio indicating selection on chimeric φgenes; c. Schematic of 5’ regulatory elements in chimeric φgenes; d. Quantification of 5’ regulatory marks in chimeric and single-parent φgenes; e. Quantification of predicted m6A marks in chimeric and single parent φgenes; f. Scatter plot distribution of parent, chimeric and single-parent φgenes by regulatory element and tissue range; g. Expression levels of *ANAPC1P2* transcript in male and female *Drosophila melanogaster* lines overexpressing *ANAPC1P2* post γ-irradiation; h. Kaplan-Meier survival curves comparing UAS-GFP control flies to those expressing *ANAPC1P2*; i. Survival comparison between control (CsBz flies) and *ANAPC1P2* expressing flies (8 lines) (males and females)(n=480 per group); j. Survival comparison between control groups and *ANAPC1P2*-expressing fly lines (males and females); k. Ectopic expression of different *ANAPC1P2* constructs in s2 cells; l. Luciferase activity of different *RMND5A-ANAPC1(ANAPC1P2)* insertion constructs in OVCAR3 cells.

### *ANAPC1P2* activation mediated through chimerism may contribute to its functions in purifying selection and restricting adaptation to genotoxic stress

To assess if the elevated expression of *ANAPC1P2* in sun-exposed skin (a tissue prone to DNA damage) identified in GTEx profiling were associated with genotoxic stress, *ANAPC1P2* expressing *Drosophila* fly lines were generated. While transgenic flies revealed no overt phenotypic abnormalities (body length and wing morphology), surprisingly *ANAPC1P2* expression was independent of GAL4 induction despite being cloned downstream of a UAS-GAL4-TATA promoter (Supplementary Figs. 4f, 4g, 4h). On exposing flies to γ-irradiation, enhanced expression of *ANAPC1P2* with significantly decreased transgenic male and female fly survival as compared with controls (parental lines, GFP-expressing / vector / additional negative controls; Figs. 3g,3h,3i). Notably, these effects were consistent across eight transgenic fly lines likely to represent random integration of the *ANAPC1P2* construct, indicating them to be independent of genomic insertion (Fig. 3j). Further, *ANAPC1P2* expression was similarly independent of GAL4 induction in S2 cells (*Drosophila* late-stage embryo cell line) suggesting the presence of intrinsic regulatory elements within the *ANAPC1P2* sequence (Supplementary Fig. 4i). To test this, we generated a series of 5′ deletion constructs, which on transfection in S2 cells were all pervasively expressed, although deletion of a 1177bp region upstream of the Transcription Start Site (TSS) reduced expression (Fig.3k; Supplementary Fig. 4j). Supporting these results, luciferase assays in a mammalian system showed that segments of the chimeric transcript (*RMND5A*: 1–636 bp; *ANAPC1*: 637–2225 bp) could drive luciferase expression, albeit at lower levels than the strong EF1α promoter (Fig.3l). TSS prediction analysis further identified two putative transcription start sites with moderate confidence in the *ANAPC1P2* sequence (Promoter 2.0; 32) (Supplementary Fig. 4j). These data identify a role for chimerism and the presence of multiple intrinsic regulatory elements in ensuring the activation of *ANAPC1P2*. A broader involvement in maintaining of sensitivity to γ-Irradiation that restricts adaptation to genotoxic stress through permissive genetic variations may imply its contribution to purifying selection and speciation.

### **φ**genes may harbour sORFs that potentially can generate novel proteins

To explore if the species- and tissue-specific expression of chimeric φgenes and associated Ka/Ks ratios suggest possible acquisition of cellular functions through generation of functional protein(s), we mapped intact open reading frames (ORFs) in the CTs and predicted high coding potential (CPC scores>0.9, Kozak scores >0.1) in chimeric and few single-φgene ORFs (Fig.4a). Interestingly, most of these maintain the same reading frame as that of the 3’ parent protein, while a few encode for entirely novel sequences (Supplementary Fig.5, Supplementary Table 4). The *PI4KAP2*-ORF encodes a truncated version of parent PI4KA protein along with deletion of a region spanning residues 1429– 1707, and hence displays conformational differences at its N-terminal and in the proximity of the deleted residues with moderate C-terminal sequence similarity (Figs.4b,4c; Supplementary Table 4). This may account for its reported loss of kinase activity and position it as a potential antagonist to PI4KA (28). *ANAPC1P2*-ORF is predicted to code for a soluble protein that retains some of the PC repeats of parent ANAPC1protein which functions in cell cycle regulation (Supplementary Fig. 6a). *HYDIN2*-ORF is predicted to code for a truncated soluble protein localizing to the cytoplasm and is structurally divergent from parent HYDIN protein (Supplementary Fig. 6b). The KIF27 φgene *ENSG000000290924* is predicted to code for a 129 amino acid protein that while retaining the parental KIF27 helix structure, adopts three unique b-sheets that form two distinct catalytic pockets (tunnel analysis using CaverWeb) suggesting that it may harbour unconventional ligand-binding sites and thereby evolved novel roles (Supplementary Fig.7a). Similarly, of the 2 chimeric *GTF2I* φgenes *GTF2IP1*-ORF is predicted to code for a truncated protein harbouring significant homology with GTF2-like repeats, is soluble in nature, likely to localize in the nucleus and may possibly function as a regulatory decoy or transcription regulator, while *GTF2IP4*-ORF may generate a novel protein that is highly dissimilar to parent protein and have unique functions (Supplementary Fig.7b). In conclusion, chimeric proteins may retain parent sequences with or without structural similarities, which deems the possibility that each may either have novel cellular functions or server to synergise with / antagonise the parent protein from which its sequences are derived.

**Fig. 4.**
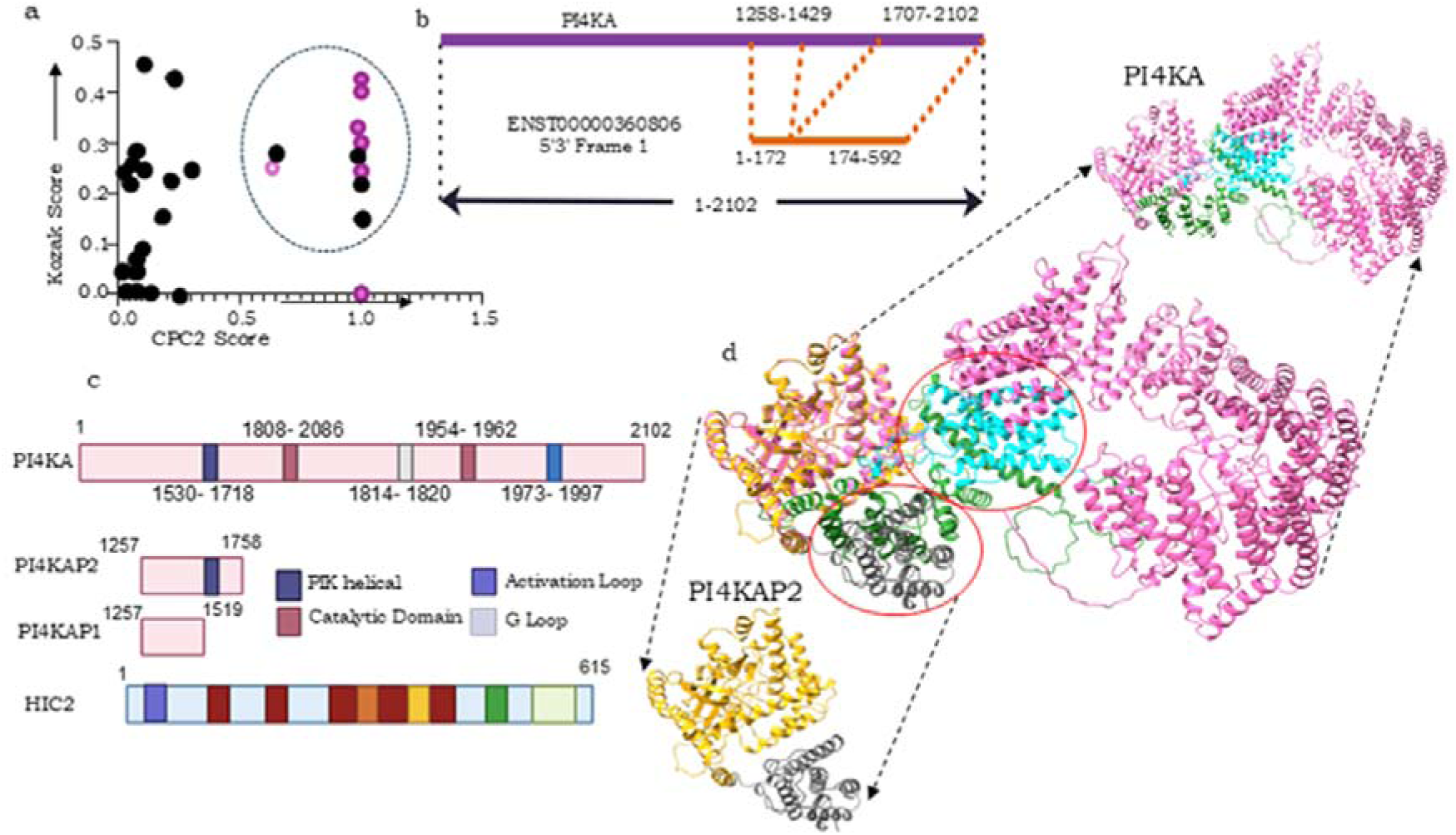
Chimeric φgenes may translate into proteins; a. Dot-plot representing Kozak vs CPC2 scores of single-parent and chimeric φgenes that predicts the latter with higher coding potential; b. *PI4KAP2* protein is likely to be a smaller derivative of *PI4KA* parent protein; c. Domain mapping of *PI4KA* and *PI4KAP2* proteins; d. Superimposition of *PI4KA* (Yellow) and *PI4KAP2* (Pink) protein structures, grey and cyan regions in *PI4KAP2* and *PI4KA* have the same amino acid (aa) sequences but present with discrete conformation that may also be attributed to the deleted region of 1429-1707aa in *PI4KAP2* (highlighted in green in PI4KA).

### Knockdown of *HYDIN2* perturbs neuronal differentiation

We further investigated the functionality of *HYDIN2* since microduplications and deletions of the human genomic locus (1q21.1) wherein it is located are associated with a spectrum of neurological and developmental disorders including macrocephaly, developmental delays, intellectual disability, psychiatric disturbances, etc. (22). Profiling of its CT across a panel of cell lines displayed a ∼10-fold higher expression in NT2 cells over others (Fig. 5a; Supplementary Fig. 8a). Hence, for investigating its relevance in cellular functions, we generated *HYDIN2* knockdown NT2 cells (KD; Supplementary Fig. 8b). RNA sequencing of KD *vs.* empty vector (EV) control cells revealed several differentially expressed genes (Supplementary Fig. 8c; Supplementary Table 5). Pathway analysis of upregulated genes indicated enrichment of processes including proximal-distal axis specification, forebrain regionalization, neuronal development and differentiation, lipid metabolism and high-density lipoprotein (HDL) particle remodeling; while downregulated pathways included cellular response to retinoic acid (RA) and negative regulation of neurogenesis (Fig. 5c; Supplementary Figs. 8d, 8e). Collectively, these results suggest the involvement of *HYDIN2* in neurodevelopmental processes.

**Fig. 5.**
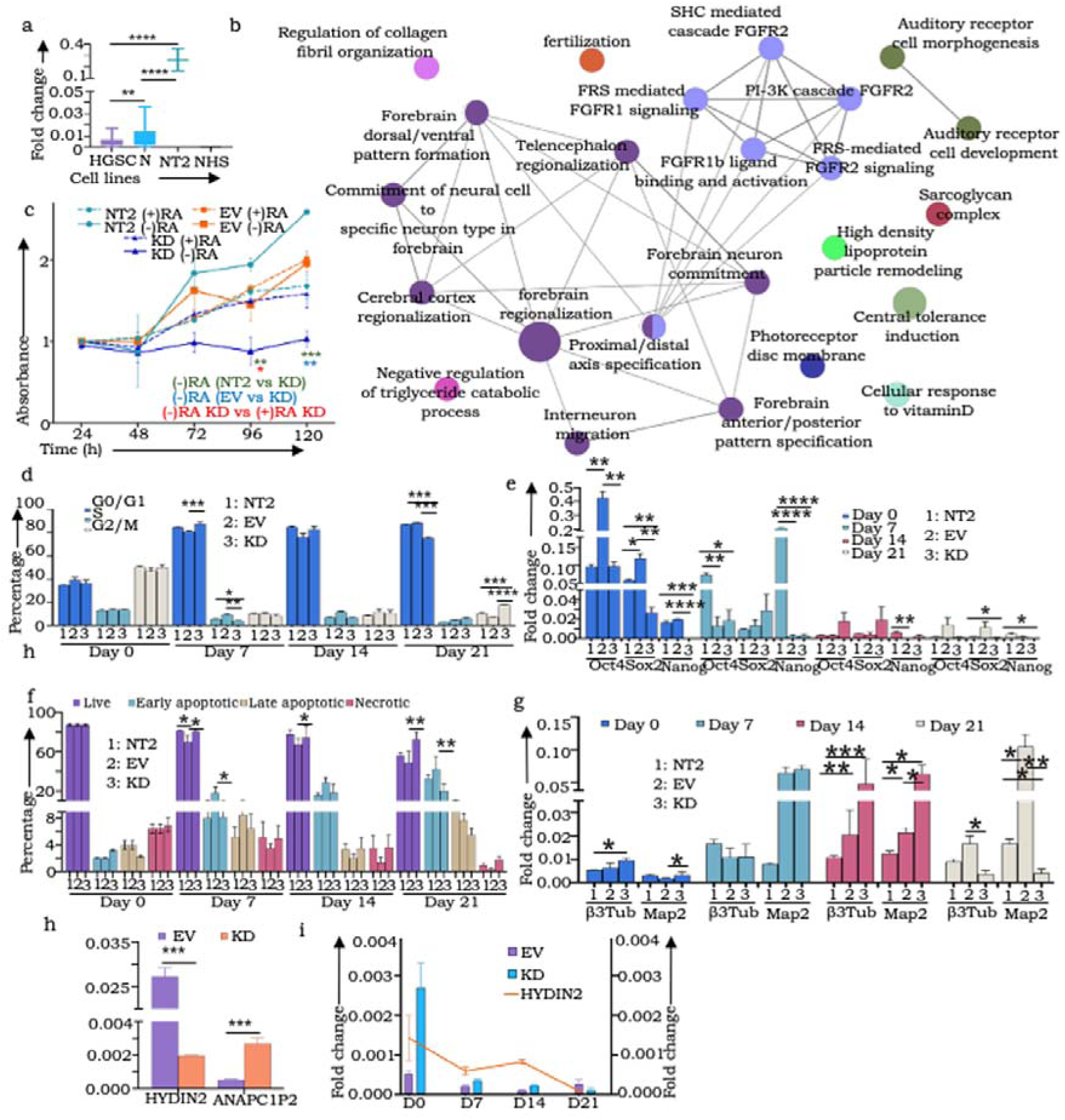
*HYDIN2* silencing perturbs cellular functions; a. Expression profiling of *HYDIN2* high grade serious ovarian cancer (HGSC), normal (N), NTERA-2 (NT2), and non-human species (NHS) cell lines; b. ClueGO networks representing differential pathways in HYDIN2-KD cells; c. Line graph representing MTT assay-based cell proliferation of all cell derivatives (empty vector, EV, KD, NT2) in the presence and absence of retinoic acid (RA); d. Propidium Iodide staining of all cell derivatives of G0/G1 synchronized cells with and without RA induction at various time points (Day-0, 7, 14, and 21); e. Expression of self-renewal markers (Oct3/4, Sox2, Nanog) at 0, 7, 14, 21 days following RA induction; f. Expression neuronal markers (β3-tubulin, Map2) at 0, 7, 14, 21 days following RA induction; g. Bar graph representing annexin V/PI staining upon RA induction till 21 days; h. mRNA levels of *ANAPC1P2* in EV and *HYDIN2* KD cells; i. *ANAPC1P2* expression in *HYDIN2*-KD and EV cells upon RA induction; Data represents mean ± SEM from three independent experiments. *p* < 0.05, p < 0.01 (Student’s *t*-test).

Since response to RA was an important perturbed differential feature on HYDIN2 KD, we evaluated RA-mediated neuronal differentiation in NT2 cells that is considered a classical model for the same (29). In the absence of RA, KD cells revealed a significantly slower rate of cell proliferation over controls (EV and NT2), however upon RA induction all cell derivatives proliferated at the same rate till Day 5 (Fig. 5d). Cell cycle analysis revealed no significant differences between EV and KD cells at steady-state; RA induction however led to significantly higher proportion of KD cells in the G0/G1 phase at Days 7 and 14, followed by an accumulation in the G2/M phase by Day 21 (Fig.5e). These subtle changes suggest lowered kinetics of cell cycle progression of KD cells through G2/M including an early exit from the cell cycle in response to RA treatment. In alignment with these observations, uninduced KD cells displayed reduced expression of self-renewal markers (Sox2, Oct4, Nanog) over controls while neuronal markers β3-tubulin and MAP2 were enhanced on RA induction over control cells, peaking at Day 14 followed by a sharp decline by Day 21 (Figs.5f, 5g). Concurrently, downregulation of cell proliferation genes including Cyclin D was also evident (Supplementary Fig. 8f). Functionally, this was reflected in significant reduction in the frequency and size of adherent colonies formed by KD cells and total failure to organize into well-defined, organized non-adherent 3D spheroids in the absence of RA treatment as compared with controls. Surprisingly, following RA treatment while no adherent colonies were generated either group of cells, KD cells generated larger spheroids compared to controls (Supplementary Figs. 9a, 9b). All these features led us to surmise insufficient build-up of critical mass of KD progenitors at initial phases followed by impaired maturation into neurons. In exploring additional pathways likely to be perturbed, we identified a lower frequency of KD cells in early apoptosis following RA treatment as compared with controls, along with downregulation of pro-apoptotic genes (BAX, PUMA, NOXA, Caspase 8 albeit with elevated BCL2 expression (Fig. 5h; Supplementary Figs. 9c,9d). Interestingly, RNA sequencing of KD cells demonstrated upregulation of several novel φgenes including a five-fold upregulation of *ANAPC1P2*; the latter expression was however completely abolished on RA induction (Figs.5b, 5i). This possibly may extend the involvement of *ANAPC1P2* and other φgenes with similar functions in genotoxicity, possibly encompassing a broader role in mitigating / effecting responses to environmental stress. While a further exploration towards a deeper mechanistic understanding of these findings is necessitated, we can surmise that *HYDIN2-*KD cells exhibit impaired neuronal maturation but resist apoptosis, potentially resulting in the persistence of incompletely differentiated cells assigning it a subtle regulatory role in core cellular processes of cell cycle progression, cellular organization, balancing self-renewal with apoptosis and ensuring proper neuronal maturation especially in response to retinoic acid.

## Discussion

A novel revelation of our study is of chimeric φgenes generated through complex segmental duplications, intron fusions and/or sequence rearrangements of two or more genes that give rise to CTs. This adds a new facet to the understanding of CTs currently believed to emerge through readthrough transcription, aberrant splicing, TAD-mediated chromatin reorganization or retrotransposition (30–34). The human-specific expression of chimeric φgenes to generate new regulatory RNAs or proteins further assigns them an important role in lineage-specific evolution and divergence, through either purifying or positive selection. Several single-parent transcribed φgenes such as mouse *Makorin1-p1*, despite being subject to random mutations generate proteins with partially conserved parental coding potential, retain functional constraints, and are reportedly under purifying selection (12, 35–41). Consistent with this, Ka/Ks analysis of four from the five chimeric φgenes in our study revealed signatures of purifying selection, suggesting persistence of their rearranged sequences under relaxed constraints. All five φgenes also harbour ORFs and can potentially generate small proteins retaining parental domains likely to display dominant-negative, decoy, or novel effects. Such chimeric φgenes may hence be coopted for regulatory- and/or retained for neo-functionalization. Surprisingly, *PI4KAP2* earlier reported to encode a truncated PI4KA kinase that disrupts phosphatidylinositol signaling by forming non-functional complexes disrupting or alternative pathways, was not realized to be chimeric (28). Divergence of regulatory elements of φgenes or a provision of alternative regulatory elements acquired during chimerism can drive species-specific traits and contribute to speciation through development of reproductive barriers and isolation, hybrid misexpression leading to perturbed developmental pathways and niche adaptation (42, 43). This is exemplified in instances emerging from insertion of mitochondrial NUMT φgenes near fertility genes like *RNF141*, or divergence of regulatory elements in *Drosophila* that contribute to hybrid incompatibilities (44, 45). Our study indicated a similar role for *ANAPC1P2* in responding to environmental genotoxic stress by eliminating unfit populations, reinforcing the essentiality of purifying selection.

Notably, the chimeric φgenes studied are present in functionally rich, human disease-linked loci associated with congenital, neurodevelopmental syndromes. The human 1q21.1 locus recognized as a hotspot of SD-driven innovation and recurrent deletions is linked with neurodevelopmental disorders (46–48). This locus houses *HYDIN2* and drives its human-specific expression through a 5′ promoter-like element active in the cerebellum and cortex, implicating it in forebrain patterning and neurodevelopment (49). Our results affirm this as depletion of *HYDIN2* perturbs neuronal differentiation while avoid lethality by upregulating compensatory φgene networks that subtly yet crucially shape neural lineage divergence either by elimination of unfit populations, or buffer fitness and mitigate phenotypic effects to maintain cellular resilience (50–52). Likewise, other chimeric φgenes of our study at 7q11.23 (Williams-Beuren syndrome), 22q11.2 (DiGeorge syndromes), 9q21.33 (global developmental delay), and 2p11.2(mild-to-moderate developmental delay) represent loci enriched in genes regulating cell cycle, differentiation, and apoptosis (53). Aligning our findings with broader associations between lineage-specific SDs generated *de novo* genes with enhanced neuronal connectivity, cognitive capacity, and brain plasticity, suggest chimeric φgenes to persist and contribute to traits that shape human evolution and speciation (54). Future functional investigations will be key to fully uncovering their specific roles.

## Methods

### Sequence analyses

14 chimeric transcripts (CTs) exhibiting sequence homology with known φgenes were previously identified by analyzing the longest read sequences using NCBI (30) and Ensembl BLAST tools (23). The original dataset comprised of 137 CTs demonstrating > 50% query coverage with reported φgenes (103 from NCBI, 42 from Ensembl, with 7 overlapping entries). This dataset was re-evaluated with increased stringency through manual curation of transcript annotation, redundancy elimination, query cover (>95%), RNA-seq expression data (presence in ≥10% of tumor samples) and verification of a 21-base sequence match flanking both sides of the fusionpoint to refine the list of candidates to focus on 9 CTs and the φgenes from which they were generated. For mapping out the chimeric introns generated during segmental duplications and gene fusions, intronic sequences flanking the identified chimeric junctions were extracted from Ensembl genome assemblies (GRCh37 and GRCh38) and aligned against their corresponding parental introns using BLAST. To map potential segmental duplications, sequences corresponding to the φgenes were aligned using NCBI BLAST across hominoid genomes (Human, Chimpanzee, Orangutan, Gibbon and Gorilla) with a minimum alignment length of 1000 base pairs used as the criterion for identifying duplication events. Splice site strength and fidelity were evaluated using Spliceator 2.0 (http://www.lbgi.fr/spliceator/).

### Determination of conserved sequences

Evolutionary conservation of parental genes was assessed using GenTree focusing specifically on vertebrate-specific lineage conservation. To investigate the conservation of the specific parental exons, introns involved in the fusion events and chimeric fusionpoint sequences, these were queried against the genomic and transcriptomic databases of 23 vertebrate species (including Human, Chimp, Orangutan, Gibbon, Rhesus, Marmoset, Rabbit, Guinea Pig, Rat, Mouse, Horse, Dog, Cow, Elephant, Opossum, Platypus, Chicken, Zebra finch, Lizard, X. tropicalis, Stickleback, Tetraodon and Zebrafish) using NCBI BLASTn.

### Cell culture and Functional Assays

High-grade serous carcinoma (HGSC) cell lines derived from patient ascites (A2, A4, and G1M2), five human lymphoblastoid cell lines (LBLs) derived from peripheral blood and five primary human skin fibroblast (SF) cultures, were previously established in the laboratory (55)(29). Additional cell lines used in this study [OVCAR3 (RRID: CVCL_0465), OVCAR4 (RRID: CVCL_1627), A2780 (RRID: CVCL_0134), MDCK (RRID: CVCL_0422), CP70 (RRID: CVCL_0135), and CV1 (RRID: CVCL_0229)] were obtained from the NCCS Cell Repository. IOSE364 (RRID: CVCL_5540), FT282 (RRID: CVCL_A4AX), HEK293T (RRID: CVCL_0063), and *Drosophila* S2 cells (RRID: CVCL_Z232) cells were kindly gifted by Dr. Ray (ACTREC, Mumbai), Dr. Bhat (IISc, Bangalore), Dr. Shiras and Dr. Majumdar (NCCS, Pune) respectively. Culture protocols followed per previous reports include NTERA-2 cells [NT2; RRID: CVCL_0034; (29)], OV90 (RRID: CVCL_3768), OVMZ6 (RRID: CVCL_4005), OVCA420 (RRID: CVCL_3935), OVCA432 (RRID: CVCL_3769), and CAOV3 (RRID: CVCL_0201) (56). All cell lines were authenticated by short tandem repeat (STR) profiling and confirmed to be mycoplasma-free by testing at the NCCS Cell Repository. The NCCS Institutional Ethics Committee (IEC) (tissue samples for primary culture - skin and peripheral blood) and Institutional Biosafety Committee (IBSC) approved all experiments that were conducted in accordance to the law and institutional guidelines.

Cell cycle analysis was carried out using Propidium Iodide (Invitrogen™, #P1304MP) and RNAaseA (Thermo Scientific™, # EN0531). Briefly, 1*10^5^ cells of all cell derivatives were seeded in 6 well plates, synchronization was achieved through serum starvation for 24hrs. Cells were harvested on Days 0,7,14; suspensions were washed with 1X PBS, fixed with 70% ethanol, and further subjected to FACS acquisition. Apoptosis assays were performed with cell suspensions at mentioned time points washed and resuspended in 1X binding buffer followed by staining with Annexin V-FITC for 15 minutes. Data was acquired using FITC and PE channels on BD FACSCanto™ II System and FlowJo_V10 software used for data analysis.

5,000 cells of all cell derivatives were seeded in a 96-well plate and stained with MTT (Sigma, #M5655-100MG-5mg/ ml) at specific time points (24, 48, 72, 96, and 120 hours) as described previously (29), with and without RA induction. For Clonogenicity assay, RA-induced and uninduced cell derivatives were harvested and seeded in 6-well plate (1000/ well). After incubation for 10 days, colonies were fixed and stained with crystal violet as described earlier (57). For Spheroid assay, 5000 cells/well was seeded in 24-well low attachment plate, and grown in serum free media. Images were captured on Olympus FV3000 microscope; Image J software was used for further analyses.

For RA-induced neuronal differentiation of all cell derivatives of NT2, cells were seeded in 6-well plates (40,000/well), cultured in medium supplemented with 10=μM all-trans retinoic acid (ATRA; Sigma-Aldrich, #R2625) for 21 days to induce differentiation, as described earlier (29). Media was replaced every 4th day, and cell pellets were collected in replicates at various time points (Days 0, 7, 14, 21 after RA treatment) for further analysis.

### Oligo design, RNA isolation, cDNA synthesis, PCR, real time PCR and Sanger sequencing

Oligos for amplification around the fusionpoint were designed using GeneRunner software (Ver.5.1.01) and synthesized by Sigma-Aldrich (25 nM; sequences available upon request). RNA extraction from cell lines and primary cultures performed using TRIzol reagent (RNAiso Plus TAKARA, #9108) was followed by cDNA synthesis and quantification (DeNovix spectrophotometer). PCR amplification was performed with TaKaRa DNA Polymerase (#R050B), amplified products resolved on a 1.8% agarose gel (Sigma-Aldrich, #A9539– 100G), resulting bands excised and purified using the QIAEX II Gel Extraction Kit (Qiagen #20021, #20051). Sanger sequencing of purified amplicons was conducted at the NCCS Sequencing Facility, and sequence data analyzed using SnapGene Viewer (39). Quantitative PCR (qPCR) was conducted on a CFX96 Touch Deep Well Real-Time PCR Detection System (Bio-Rad), using TB Green® Premix Ex Taq™ (Tli RNase H Plus) SYBR Green Master Mix. Reactions included 1 pMol of primer mix and 100ng cDNA. GAPDH was used as the endogenous normalization control.

### Cloning and transfections

Insert and vector DNA fragments (pUAST and pGL3) were enzymatically digested using restriction endonucleases appropriate for each cloning strategy. For the full-length and Δ constructs (Δ1 and Δ2) in pUAST, KpnI-HF (NEB, #R3142S) and NotI (NEB, #R0189S) were employed; Δ UP (upstream region) was achieved using NdeI (NEB, #R0111S) and BglII (NEB, #R0144S); Δ TATA promoter using SbfI-HF (NEB, #R3642S) and BglII. For insertion of RMND5A and ANAPC1 sequences into the pGL3 vector, digestion was performed using KpnI and HindIII (NEB, #R0104S). All ligation reactions were carried out using T4 DNA ligase (TAKARA, #2011B) according to the manufacturer’s instructions.

shRNA oligonucleotide sequences against *HYDIN2* fusionpoint were designed using The RNAi Consortium (TRC) public portal hosted by the Broad Institute (https://portals.broadinstitute.org/gpp/public/). Forward and reverse oligonucleotides were reconstituted to a final concentration of 0.1 nmol/μL. For annealing, a 25 μL reaction mixture was prepared containing 11.25 μL of each oligonucleotide and 2.5 μL of 10X annealing buffer (NEB Buffer 2 / reconstituted with 1 M NaCl + 100 mM Tris-HCl, pH 7.4). The mixture was heated in boiling water, allowed to gradually cool to room temperature to facilitate duplex formation, and diluted 1:100 in 0.5× annealing buffer. 1 μL of this mixture with 10–20 ng linearized pLKO.1-TRC vector T4 DNA ligase (TAKARA, #2011B) was used in a 20 μL ligation reaction. Recombinant constructs were subsequently transformed into chemically competent DH5α cells. Plasmid DNA was extracted using the Nucleospin Plasmid Isolation Kit (MACHEREY-NAGEL, #740588.250). Sequences encoding shRNAs were confirmed by Sanger DNA sequencing using U6 promoter-specific primer.

Lentiviral particles were generated and concentrated as described earlier (58). NT2 cells were transduced with the lentiviral particles in the presence of Polybrene (Sigma-Aldrich, #TR-1003), and selected using puromycin (Gibco™, #A1113803) at a final concentration of 0.8 μg/mL. Transfection of *Drosophila* S2 cells was performed using Effectene Transfection Reagent per following the manufacturer’s instructions (QIAGEN, #301425). Luciferase reporter assays were performed using the Dual-Glo™ Luciferase Assay System per manufacturer’s protocol (Promega, #E2920).

### Prediction of Coding potential (ORFs, Kozak sequence) and evolutionary selection (GC content and K_a_/K_s_ ratios)

Canonical parent protein sequences were obtained from Uniprot (https://www.uniprot.org/) (42). Putative proteins were predicted from CT sequences across all six reading frames using Expasy translation tool (https://web.expasy.org/translate/) (43). Coding potential of transcripts was assessed using the Coding Potential Calculator 2.0 (CPC2) (https://cpc2.gao-lab.org/) (44). Transcripts with a coding probability greater than 0.6 were classified as protein-coding. To identify potential Kozak sequences, the ATGpr tool (http://atgpr.dbcls.jp/) was utilized. Open reading frames (ORFs) were considered putative if they exhibited a reliability score exceeding 0.1, including a stop codon, and had a minimum length of 10 codons.

GC content was calculated using a GC content calculator (https://en.vectorbuilder.com/tool/gc-content-calculator.html). Full-length transcript sequences of chimeric φgenes, their parental genes, and single-parent φgenes were used as input to assess and compare GC content across genes. Evolutionary selection pressures between parental genes and their φgenes were assessed using initially, the ParaAT tool (https://ngdc.cncb.ac.cn/tools/paraat) to construct protein-coding DNA alignments from inputs including CDS and protein/peptide/amino acid sequences for the corresponding proteins (as FASTA files) and a list of parental and chimeric proteins (CSV file). This output was provided as input to the K_a_K_s__Calculator (https://ngdc.cncb.ac.cn/tools/kaks), that applies maximum likelihood method of Model Selection/Model Averaging to K_a_, K_s_ values and their ratio (ω) in addition to a p-value (Fisher’s test). K_a_/K_s_ < 1 indicates purifying selection, K_a_/K_s_ = 1 represents neutral selection, while K_a_/K_s_ > 1 suggests positive selection).

### Transgenic Drosophila Line Generation, **γ**-irradiation, and Survival Assay

Transgenic Drosophila lines were generated by microinjection of pUAST-*ANAPC1P2* construct into *w[1118]* embryos (WellGenetics, Taipei). Eight UAS-*ANAPC1P2* transgenic lines were established using this method. Fly morphology was examined (Nikon SMZ18), and transgene expression was confirmed by qPCR of cDNA from total RNA. For irradiation assays, 1–3-day-old transgenic and control flies were exposed to 700 Gy ionizing radiation, and survival was monitored daily.

### Predictions of protein structure and function

The Alphafold server (https://alphafoldserver.com) was used to model structures of parental and chimeric proteins. CIF files were loaded into ChimeraX (https://www.cgl.ucsf.edu/chimerax/) for visualization, and structures downloaded as coordinate PDB files. Superimposition of parent and chimeric proteins were done using the “Matchmaker” feature in ChimeraX. The protein structure comparison service PDBeFold (https://www.ebi.ac.uk/msd-srv/ssm/) was used for comparison of parent and chimeric protein structures. DeepLoc2.1 (https://services.healthtech.dtu.dk/services/DeepLoc-2.1/) was used to predict subcellular localization and the associated membrane type of eukaryotic proteins with a cutoff probability 0.6. Interpro (https://www.ebi.ac.uk/interpro/), was used for identification of functional domains.

### Sequence-Based RNA Adenosine Methylation Site Predictor Analysis (SRAMP)

The SRAMP tool (http://www.cuilab.cn/sramp/) was used to predict m6A modification sites in RNA sequences. SRAMP assigns a prediction score for each potential m6A site sequence. These scores are classified into confidence levels: very high (99%), high (95%), moderate (90%), and low (85%), based on specificity from cross-validation tests. Scores were obtained for chimeric and single parent φgene mRNA sequences from National Center for Biotechnology Information (NCBI) and Ensembl38 (GRCh38) and analyzed in mature mRNA mode. Predicted m6A marks were used to compare the frequency of m6A modifications on 5’ and 3’ Untranslated regions (UTR) between chimeric and single parent φgene sequences.

### RNA sequencing and data analysis

Cell pellets were submitted to MedGenome Labs Ltd for RNA isolation, library preparation, and sequencing on Illumina HiSeqX (60M, 2 x 150 bp reads/sample). A standard pipeline (QC, preprocessing, alignment, expression estimation, DESeq2 normalization) identified differentially expressed genes (p<0.05, Fold Change>2&<-2). Pathway analysis of differentially expressed candidate transcripts was achieved through REACTOME and gene set enrichment analysis (GSEA software v 2023.2 - biocarta, kegg, Reactome, wiki pathways, gene ontology, GO - molecular function, cellular compartment, and biological process sets). Further pathway enrichment analysis was achieved using ClueGO (v2.5.10) (59) as a Cytoscape (3.9.1) plugin (21) with GO-biological process, GO (Cellular Component, Immune System Process, Molecular Function), KEGG, Reactome (Pathways and Reactions) considering minimum 2 genes per term, Kappa score 0.4 cutoffs, p<=0.05.

### Statistical analysis

Unless otherwise mentioned, all experiments were conducted in triplicates (n=3, for both experimental and biological replicates). Data visualization and statistical analysis (paired t-test) were carried out using GraphPad Prism software (version 8.4.2, 67). Statistical significance was defined as *p<0.05, **p<0.01, ***p<0.001, and ****p<0.0001. Standard deviation is represented as error bars.

## Availability of data

The RNA-sequencing datasets supporting the conclusions of this article are available in the Gene Expression Omnibus (GEO) repository (GSE298099).

## Acknowledgments

We sincerely thank Prof. Judith Clements (Translational Research Institute, Australia) for the generous gift of cell lines OVCA420, OVCA432, and CAOV3; Dr. S. Mok (MD Anderson Cancer Center, TX) for OV90; Prof. V. Magdalen (Klinische Forschergruppe der Frauenklinik, TU München) for OVMZ6; Dr. Pritha Ray (Advanced Centre for Treatment, Research & Education in Cancer, Tata Memorial Centre) for IOSE364; Dr. Ramray Bhat (Indian Institute of Science) for FT282; Dr. Majumdar and late Dr. Shiras (NCCS) for providing S2 and HEK293T respectively. The efforts of Meghna Killaru (GKAP-KIF27, EMB-TRIM43, BANP-RUNDC2A, RP4-HYDIN), Mrunal Desai (PMS2P2-POLR2J3), Ankita More (HIC2-PI4KA), and Vartika Khanchandani (RMND5A-ANAPC1) in validating fusion point sequences are gratefully acknowledged. We extend our thanks to Swarali Gogate for her contributions in standardization of NT2 cell differentiation. The support rendered by NCCS Cell Repository in providing cell lines, culture media, and STR profiling services is appreciated. We also thank the NCCS flow cytometry, bioimaging, and sequencing core facilities for their support in data collection. Some of the illustrations in this manuscript were created with BioRender.com.

## Authors contribution

ARJ, SS, YLK: Contributed equally towards data curation, experimentation, analysis, data interpretation, writing – original draft, editing; KT: Experimentation, data analysis; SP: Data analysis; AV: Experimentation; GC: Experimentation; AMM: Experimentation; SAB: Conceptualization, methodology, supervision, data analysis and interpretation, project administration, manuscript writing, editing and finalization.

## Classification

Major category: Biological sciences

Minor category: Evolution, Cell Biology

## Supplementary Information

Supplementary figures: Supplementary figures 1 to 9

Supplementary table 1: List of all φgenes from NCBI and Ensembl 38

Supplementary table 2: Exon and intron conservation of all 5 φgenes in 23 vertebrate species

Supplementary table 3: Splicing site predictions around intron fusion and their respective parental sequences for 5 φgenes.

Supplementary table 4: Chimeric φgene ORF-associated protein structure-function prediction

Supplementary table 5: Differentially expressed genes in HYDIN2 Knockdown compared to empty vector control

## Declarations

### Ethics approval and consent

All experiments were approved by the NCCS Institutional Ethics Committee (IEC) and Institutional Biosafety Committee (IBSC) and were conducted in accordance with institutional guidelines and applicable laws.

### Consent for publication

None of the authors have any potential competing interests associated with the publication.

