## Supplementary Table.1 for "Evolutionary and functional dynamics of chimeric pseudogenes (φgenes)"

**Supplementary table 5: Differentially expressed genes in HYDIN2 Knockdown compared to empty vector control**

### Upregulated genes

| Gene | Gene symbol | log2FoldChange | pvalue |
|---|---|---|---|
| ENSG00000206652.1 | RNU1-1 | 14.4294269 | 0.000408 |
| ENSG00000174080.12 | CTSF | 8.05863493 | 0.00012 |
| ENSG00000184564.11 | SLITRK6 | 7.418480306 | 0.000479 |
| ENSG00000214456.8 | PLIN5 | 7.343038797 | 0.000599 |
| ENSG00000143994.14 | ABHD1 | 7.250116695 | 0.000671 |
| ENSG00000277954.1 | ENSG00000277954 | 6.935698005 | 0.001339 |
| ENSG00000146047.7 | H2BC1 | 6.831901698 | 0.001706 |
| ENSG00000207513.1 | RNU1-3 | 6.683645738 | 0.003192 |
| ENSG00000205056.9 | LINC02397 | 6.572111021 | 0.002629 |
| ENSG00000290091.1 | ENSG00000290091 | 6.541956594 | 0.002742 |
| ENSG00000287075.2 | ENSG00000287075 | 6.486127211 | 0.003219 |
| ENSG00000196611.6 | MMP1 | 6.466029064 | 0.003574 |
| ENSG00000179082.4 | LINC02913 | 6.462765213 | 0.003371 |
| ENSG00000262265.2 | ENSG00000262265 | 6.460720439 | 0.003737 |
| ENSG00000252577.1 | SCARNA20 | 6.398440502 | 0.003964 |
| ENSG00000153266.13 | FEZF2 | 6.388110699 | 0.003633 |
| ENSG00000132837.15 | DMGDH | 6.373559933 | 0.003884 |
| ENSG00000231128.7 | ENSG00000231128 | 6.372188636 | 0.004013 |
| ENSG00000231965.4 | ENSG00000231965 | 6.367670134 | 0.003845 |
| ENSG00000224039.1 | CDYLP1 | 6.287514481 | 0.004562 |
| ENSG00000272079.2 | ENSG00000272079 | 6.276672768 | 0.004781 |
| ENSG00000164266.11 | SPINK1 | 6.235053586 | 0.004807 |
| ENSG00000187398.12 | LUZP2 | 6.225826907 | 0.005394 |
| ENSG00000270933.1 | ENSG00000270933 | 6.209573925 | 0.005249 |
| ENSG00000134594.5 | RAB33A | 6.209187344 | 0.005294 |
| ENSG00000165181.17 | SHOC1 | 6.193105933 | 0.005313 |
| ENSG00000225420.1 | ENSG00000225420 | 6.175366225 | 0.005478 |
| ENSG00000234004.4 | ENSG00000234004 | 6.162634754 | 0.00599 |
| ENSG00000259439.3 | LINC01833 | 6.139609174 | 0.006753 |
| ENSG00000007372.25 | PAX6 | 6.1291618 | 0.006618 |
| ENSG00000183389.5 | OR56A4 | 6.121048232 | 0.007115 |
| ENSG00000288835.1 | ENSG00000288835 | 6.119702173 | 0.006118 |
| ENSG00000230366.9 | DSCR9 | 6.045178304 | 0.007041 |
| ENSG00000273391.1 | ENSG00000273391 | 6.00394847 | 0.007564 |
| ENSG00000226903.3 | LINC00354 | 5.988152814 | 0.008672 |
| ENSG00000263393.1 | ENSG00000263393 | 5.984144903 | 0.007664 |
| ENSG00000260428.3 | SCX | 5.96188943 | 0.007997 |
| ENSG00000254270.1 | ERHP1 | 5.956346133 | 0.008034 |
| ENSG00000176236.6 | RPP38-DT | 5.932982086 | 0.010582 |
| ENSG00000287817.1 | ENSG00000287817 | 5.930547274 | 0.009547 |
| ENSG00000120664.11 | SPART-AS1 | 5.90823673 | 0.01053 |

| | | | |
|---|---|---|---|
| ENSG00000225746.13 | MEG8 | 5.908081086 | 0.008876 |
| ENSG00000183638.6 | RP1L1 | 5.902294007 | 0.008877 |
| ENSG00000267127.7 | ENSG00000267127 | 5.900459692 | 0.008995 |
| ENSG00000278934.1 | ENSG00000278934 | 5.895166563 | 0.009337 |
| ENSG00000184702.20 | SEPTIN5 | 5.856493329 | 0.010537 |
| ENSG00000267024.1 | ENSG00000267024 | 5.855981959 | 0.009664 |
| ENSG00000288911.2 | ENSG00000288911 | 5.850103418 | 0.010433 |
| ENSG00000197320.5 | ENSG00000197320 | 5.837891918 | 0.009924 |
| ENSG00000288586.1 | ENSG00000288586 | 5.836264547 | 0.01035 |
| ENSG00000164304.16 | CAGE1 | 5.804049185 | 0.011446 |
| ENSG00000199912.1 | Y_RNA | 5.79867337 | 0.011545 |
| ENSG00000070193.5 | FGF10 | 5.798438534 | 0.010946 |
| ENSG00000287437.1 | ENSG00000287437 | 5.776940653 | 0.013405 |
| ENSG00000286650.2 | ENSG00000286650 | 5.775565246 | 0.011167 |
| ENSG00000230224.1 | PHB1P9 | 5.768618953 | 0.012175 |
| ENSG00000176840.14 | MIR7-3HG | 5.763711423 | 0.012105 |
| ENSG00000241627.3 | UBQLN4P1 | 5.758021519 | 0.011988 |
| ENSG00000201448.1 | SNORA63C | 5.749803908 | 0.011678 |
| ENSG00000227492.2 | CHUK-DT | 5.746951056 | 0.011989 |
| ENSG00000236776.1 | RPL21P23 | 5.74363942 | 0.01216 |
| ENSG00000270175.2 | ENSG00000270175 | 5.740875231 | 0.011893 |
| ENSG00000247624.6 | CPEB2-DT | 5.737854122 | 0.013053 |
| ENSG00000254653.1 | ENSG00000254653 | 5.736671064 | 0.012378 |
| ENSG00000187553.10 | CYP26C1 | 5.718388421 | 0.012296 |
| ENSG00000203446.2 | SUGCT-AS1 | 5.716718251 | 0.01247 |
| ENSG00000162482.5 | AKR7A3 | 5.71625206 | 0.012179 |
| ENSG00000101162.4 | TUBB1 | 5.70973454 | 0.015942 |
| ENSG00000254896.1 | OPCML-IT1 | 5.695160367 | 0.013598 |
| ENSG00000185988.15 | PLK5 | 5.675845214 | 0.013931 |
| ENSG00000110245.12 | APOC3 | 5.651877144 | 0.014032 |
| ENSG00000254208.1 | ENSG00000254208 | 5.650164429 | 0.01439 |
| ENSG00000228204.2 | ENSG00000228204 | 5.641816263 | 0.014211 |
| ENSG00000207741.1 | MIR590 | 5.638032611 | 0.015593 |
| ENSG00000285884.3 | ENSG00000285884 | 5.635428295 | 0.015202 |
| ENSG00000266282.1 | UBL5P2 | 5.629843573 | 0.015053 |
| ENSG00000285928.1 | ENSG00000285928 | 5.622960913 | 0.014669 |
| ENSG00000163283.7 | ALPP | 5.584285863 | 0.01686 |
| ENSG00000264448.5 | MIR378D2HG | 5.579773281 | 0.015909 |
| ENSG00000229671.3 | LINC01150 | 5.577092113 | 0.019156 |
| ENSG00000239212.2 | RPL6P7 | 5.572061038 | 0.016068 |
| ENSG00000233560.2 | KRT8P39 | 5.569264762 | 0.016985 |
| ENSG00000289908.1 | ENSG00000289908 | 5.563311034 | 0.015795 |
| ENSG00000287724.1 | ENSG00000287724 | 5.560191835 | 0.01698 |
| ENSG00000232613.7 | LINC02576 | 5.548331421 | 0.020199 |
| ENSG00000277171.1 | ENSG00000277171 | 5.545403237 | 0.01922 |
| ENSG00000070388.12 | FGF22 | 5.544753844 | 0.016671 |
| ENSG00000287078.1 | ENSG00000287078 | 5.525986737 | 0.017365 |

| | | | |
|---|---|---|---|
| ENSG00000171747.9 | LGALS4 | 5.522723564 | 0.017 |
| ENSG00000253838.1 | ENSG00000253838 | 5.509060094 | 0.018636 |
| ENSG00000256407.2 | ENSG00000256407 | 5.506372793 | 0.019533 |
| ENSG00000225093.1 | RPL3P7 | 5.490291188 | 0.018867 |
| ENSG00000199683.1 | RN7SKP185 | 5.48854135 | 0.018246 |
| ENSG00000279026.1 | ENSG00000279026 | 5.483766912 | 0.018056 |
| ENSG00000290041.1 | ENSG00000290041 | 5.479806186 | 0.018831 |
| ENSG00000227161.1 | ENSG00000227161 | 5.469939796 | 0.019348 |
| ENSG00000232814.2 | COL4A2-AS1 | 5.454105064 | 0.020932 |
| ENSG00000259447.1 | ENSG00000259447 | 5.452095792 | 0.020378 |
| ENSG00000283405.2 | ENSG00000283405 | 5.447998422 | 0.02174 |
| ENSG00000246820.2 | RIC3-DT | 5.444785709 | 0.01945 |
| ENSG00000290545.1 | INTS4P1 | 5.434236059 | 0.019834 |
| ENSG00000275709.1 | ENSG00000275709 | 5.423029558 | 0.021156 |
| ENSG00000135355.4 | GJA10 | 5.422074774 | 0.020429 |
| ENSG00000267203.1 | SNRPGP4 | 5.41988921 | 0.021175 |
| ENSG00000260123.2 | CARMAL | 5.418939342 | 0.020724 |
| ENSG00000218416.4 | GPC1-AS1 | 5.413721152 | 0.02276 |
| ENSG00000274750.3 | H3C6 | 5.409841874 | 0.020907 |
| ENSG00000244538.1 | ENSG00000244538 | 5.409519994 | 0.021946 |
| ENSG00000275383.1 | ENSG00000275383 | 5.40161541 | 0.020718 |
| ENSG00000185053.14 | SGCZ | 5.393676253 | 0.022013 |
| ENSG00000289229.1 | ENSG00000289229 | 5.392650976 | 0.02539 |
| ENSG00000287980.1 | ENSG00000287980 | 5.391548772 | 0.022365 |
| ENSG00000275160.1 | RPL7AP45 | 5.390451792 | 0.022323 |
| ENSG00000121764.12 | HCRTR1 | 5.383951333 | 0.02323 |
| ENSG00000238217.7 | LINC01877 | 5.382050096 | 0.024524 |
| ENSG00000232587.1 | EEF1A1P3 | 5.378656413 | 0.022196 |
| ENSG00000237249.1 | ENSG00000237249 | 5.374793037 | 0.024143 |
| ENSG00000272721.5 | EIF2B5-DT | 5.370415172 | 0.024491 |
| ENSG00000161664.7 | ASB16 | 5.365073452 | 0.022181 |
| ENSG00000122592.8 | HOXA7 | 5.364994798 | 0.021927 |
| ENSG00000250980.1 | NSA2P6 | 5.364436015 | 0.022388 |
| ENSG00000225573.4 | RPL35P5 | 5.358167755 | 0.022617 |
| ENSG00000286931.3 | ENSG00000286931 | 5.349960272 | 0.024233 |
| ENSG00000179057.14 | IGSF22 | 5.347959071 | 0.023553 |
| ENSG00000274001.1 | ENSG00000274001 | 5.341270418 | 0.028397 |
| ENSG00000235138.1 | ENSG00000235138 | 5.340445148 | 0.022817 |
| ENSG00000226148.1 | SLC25A39P1 | 5.337323275 | 0.023267 |
| ENSG00000265185.6 | SNORD3B-1 | 5.336665717 | 0.024599 |
| ENSG00000275084.4 | SNORD91B | 5.335175675 | 0.026263 |
| ENSG00000289126.2 | ENSG00000289126 | 5.33175438 | 0.024097 |
| ENSG00000277991.4 | ENSG00000277991 | 5.329573989 | 0.035198 |
| ENSG00000261924.1 | ENSG00000261924 | 5.328608496 | 0.023281 |
| ENSG00000270300.2 | PHACTR2P1 | 5.326875979 | 0.027324 |
| ENSG00000256995.8 | LINC02955 | 5.321652376 | 0.025692 |
| ENSG00000264569.2 | DCXR-DT | 5.320458005 | 0.025724 |

| | | | |
|---|---|---|---|
| ENSG00000072858.11 | SIDT1 | 5.311670579 | 0.024481 |
| ENSG00000249737.1 | ENSG00000249737 | 5.309055449 | 0.027408 |
| ENSG00000270402.1 | ENSG00000270402 | 5.300167507 | 0.026726 |
| ENSG00000199032.1 | MIR425 | 5.296848033 | 0.024925 |
| ENSG00000150275.20 | PCDH15 | 5.290928053 | 0.029316 |
| ENSG00000227827.3 | PKD1P2 | 5.287857333 | 0.024894 |
| ENSG00000260012.1 | ENSG00000260012 | 5.286194681 | 0.025525 |
| ENSG00000234946.1 | SDHCP3 | 5.285384773 | 0.027149 |
| ENSG00000221303.1 | SNORA79 | 5.283917311 | 0.029629 |
| ENSG00000250240.6 | ENSG00000250240 | 5.282833559 | 0.025019 |
| ENSG00000285555.1 | ENSG00000285555 | 5.280819102 | 0.029753 |
| ENSG00000225486.1 | ENSG00000225486 | 5.267185785 | 0.0281 |
| ENSG00000163914.5 | RHO | 5.266295833 | 0.02733 |
| ENSG00000260172.2 | LINC01413 | 5.266215849 | 0.02727 |
| ENSG00000286576.1 | ENSG00000286576 | 5.26137212 | 0.0265 |
| ENSG00000258274.1 | CRADD-AS1 | 5.258547568 | 0.027786 |
| ENSG00000237668.1 | RPS15AP38 | 5.255111972 | 0.026853 |
| ENSG00000228126.1 | FALEC | 5.251309261 | 0.027641 |
| ENSG00000181322.15 | NME9 | 5.250075936 | 0.027099 |
| ENSG00000243107.1 | ENSG00000243107 | 5.247801357 | 0.027857 |
| ENSG00000116690.13 | PRG4 | 5.245475889 | 0.02909 |
| ENSG00000289032.1 | ENSG00000289032 | 5.24541816 | 0.026783 |
| ENSG00000286365.1 | ENSG00000286365 | 5.244311946 | 0.029199 |
| ENSG00000235834.1 | ENSG00000235834 | 5.23901719 | 0.027148 |
| ENSG00000287348.1 | ENSG00000287348 | 5.236648823 | 0.028275 |
| ENSG00000285793.1 | ANAPC1P2 | 5.231545785 | 0.027914 |
| ENSG00000288156.1 | ENSG00000288156 | 5.229305483 | 0.027546 |
| ENSG00000232939.3 | ENSG00000232939 | 5.228058751 | 0.027326 |
| ENSG00000229207.1 | SERPINH1P1 | 5.226067664 | 0.029561 |
| ENSG00000267784.1 | ENSG00000267784 | 5.224020631 | 0.028107 |
| ENSG00000271474.1 | UNC5C-AS1 | 5.222140598 | 0.032635 |
| ENSG00000184933.6 | OR6A2 | 5.219886139 | 0.028151 |
| ENSG00000137871.21 | ZNF280D | 5.219143972 | 0.033925 |
| ENSG00000138083.5 | SIX3 | 5.218269366 | 0.030523 |
| ENSG00000272983.1 | ENSG00000272983 | 5.214144955 | 0.03397 |
| ENSG00000174417.3 | TRHR | 5.213244679 | 0.02974 |
| ENSG00000065325.13 | GLP2R | 5.21271073 | 0.03152 |
| ENSG00000215452.5 | ZNF663P | 5.20869468 | 0.028274 |
| ENSG00000203647.2 | ENSG00000203647 | 5.208241952 | 0.029009 |
| ENSG00000264458.1 | ENSG00000264458 | 5.207007418 | 0.03716 |
| ENSG00000234332.1 | BCAS2P2 | 5.205217837 | 0.032758 |
| ENSG00000269506.2 | ENSG00000269506 | 5.204638595 | 0.02884 |
| ENSG00000289198.1 | ENSG00000289198 | 5.201764668 | 0.02965 |
| ENSG00000199325.1 | RNU4-39P | 5.193427892 | 0.029301 |
| ENSG00000287029.1 | ENSG00000287029 | 5.193358181 | 0.028829 |
| ENSG00000225963.8 | ENSG00000225963 | 5.191488948 | 0.030111 |
| ENSG00000229414.2 | KCNQ1-AS1 | 5.187787823 | 0.031591 |

| | | | |
|---|---|---|---|
| ENSG00000272817.1 | ENSG00000272817 | 5.186818209 | 0.029126 |
| ENSG00000240776.1 | ENSG00000240776 | 5.186267122 | 0.030495 |
| ENSG00000267289.1 | PIN1-DT | 5.182611602 | 0.029542 |
| ENSG00000248367.2 | ENSG00000248367 | 5.176364969 | 0.032269 |
| ENSG00000235514.1 | LLPHP2 | 5.172394185 | 0.030764 |
| ENSG00000170476.16 | MZB1 | 5.171687661 | 0.030955 |
| ENSG00000286679.1 | ENSG00000286679 | 5.171199202 | 0.032931 |
| ENSG00000261535.1 | ENSG00000261535 | 5.169616028 | 0.033042 |
| ENSG00000267498.2 | UQCRFS1-DT | 5.15790228 | 0.03338 |
| ENSG00000229276.1 | REV3L-IT1 | 5.152128139 | 0.034524 |
| ENSG00000244256.3 | RN7SL130P | 5.147904003 | 0.03132 |
| ENSG00000232969.1 | ENSG00000232969 | 5.146672285 | 0.031216 |
| ENSG00000183981.8 | MAGEA13P | 5.146204776 | 0.032127 |
| ENSG00000254536.2 | ENSG00000254536 | 5.14608298 | 0.032466 |
| ENSG00000177465.5 | ACOT4 | 5.143906583 | 0.034513 |
| ENSG00000103355.14 | PRSS33 | 5.14372932 | 0.03232 |
| ENSG00000272081.1 | FCHO2-DT | 5.140965361 | 0.03376 |
| ENSG00000235358.2 | SCMH1-DT | 5.135650678 | 0.032829 |
| ENSG00000259341.2 | ENSG00000259341 | 5.128749087 | 0.032805 |
| ENSG00000255008.5 | LINC02739 | 5.12678827 | 0.035236 |
| ENSG00000276651.1 | ENSG00000276651 | 5.126229318 | 0.032612 |
| ENSG00000141579.7 | ZNF750 | 5.123821607 | 0.032214 |
| ENSG00000275710.1 | ENSG00000275710 | 5.120981522 | 0.038328 |
| ENSG00000105610.6 | KLF1 | 5.118865386 | 0.032619 |
| ENSG00000288679.1 | ENSG00000288679 | 5.118052222 | 0.036176 |
| ENSG00000289317.2 | ENSG00000289317 | 5.112481319 | 0.03358 |
| ENSG00000261829.1 | ENSG00000261829 | 5.11189471 | 0.035716 |
| ENSG00000217512.1 | ENSG00000217512 | 5.109923506 | 0.032973 |
| ENSG00000255974.8 | CYP2A6 | 5.108669988 | 0.032941 |
| ENSG00000245711.2 | NADK2-AS1 | 5.107270184 | 0.037388 |
| ENSG00000272895.1 | ENSG00000272895 | 5.09595671 | 0.034134 |
| ENSG00000286882.1 | ENSG00000286882 | 5.0955203 | 0.040524 |
| ENSG00000232470.1 | ENSG00000232470 | 5.090889182 | 0.034306 |
| ENSG00000231944.1 | PHKA1-AS1 | 5.08843612 | 0.034274 |
| ENSG00000288577.1 | ENSG00000288577 | 5.08715833 | 0.034595 |
| ENSG00000143001.5 | TMEM61 | 5.086483113 | 0.034201 |
| ENSG00000262089.1 | ENSG00000262089 | 5.086314694 | 0.035728 |
| ENSG00000140478.16 | GOLGA6D | 5.084746314 | 0.036432 |
| ENSG00000288923.1 | ENSG00000288923 | 5.084036552 | 0.037437 |
| ENSG00000259420.5 | ENSG00000259420 | 5.084030814 | 0.035775 |
| ENSG00000234546.5 | LNCTAM34A | 5.083281529 | 0.036067 |
| ENSG00000242593.8 | ENSG00000242593 | 5.077254514 | 0.036751 |
| ENSG00000286771.1 | ENSG00000286771 | 5.071304928 | 0.037445 |
| ENSG00000274010.1 | ZFP91P1 | 5.07078875 | 0.038381 |
| ENSG00000234118.1 | RPL13AP6 | 5.070427137 | 0.037026 |
| ENSG00000217930.8 | PAM16 | 5.07041871 | 0.038221 |
| ENSG00000225885.7 | ENSG00000225885 | 5.067678155 | 0.035704 |

| | | | |
|---|---|---|---|
| ENSG00000185038.15 | MROH2A | 5.06256735 | 0.038928 |
| ENSG00000289550.1 | ENSG00000289550 | 5.061156071 | 0.035912 |
| ENSG00000147206.17 | NXF3 | 5.060339159 | 0.038105 |
| ENSG00000249700.9 | SRD5A3-AS1 | 5.054733236 | 0.036953 |
| ENSG00000290127.1 | ENSG00000290127 | 5.053717954 | 0.045688 |
| ENSG00000163645.15 | ERICH6 | 5.052013178 | 0.036755 |
| ENSG00000280660.1 | ENSG00000280660 | 5.051892626 | 0.038346 |
| ENSG00000233340.1 | ENSG00000233340 | 5.050151939 | 0.036398 |
| ENSG00000224781.1 | EIF4A2P4 | 5.043412242 | 0.036797 |
| ENSG00000267172.1 | ENSG00000267172 | 5.043053614 | 0.038525 |
| ENSG00000275223.1 | ENSG00000275223 | 5.040241898 | 0.037501 |
| ENSG00000253939.1 | ENSG00000253939 | 5.039564028 | 0.037116 |
| ENSG00000291090.1 | CCDC144BP | 5.03796311 | 0.045477 |
| ENSG00000229081.1 | LINC01165 | 5.036155991 | 0.039101 |
| ENSG00000250015.1 | ENSG00000250015 | 5.033506279 | 0.037007 |
| ENSG00000267192.1 | ENSG00000267192 | 5.028236189 | 0.03736 |
| ENSG00000173662.21 | TAS1R1 | 5.026912335 | 0.037654 |
| ENSG00000291296.1 | ENSG00000291296 | 5.018725378 | 0.038109 |
| ENSG00000233615.1 | HNRNPA1P42 | 5.014713766 | 0.038116 |
| ENSG00000234562.1 | TPMTP2 | 5.014149271 | 0.043921 |
| ENSG00000122859.5 | NEUROG3 | 5.00477624 | 0.046654 |
| ENSG00000243795.1 | LINC02044 | 5.003443886 | 0.040612 |
| ENSG00000255122.1 | ENSG00000255122 | 5.001988569 | 0.039627 |
| ENSG00000257883.1 | ENSG00000257883 | 4.998510461 | 0.038892 |
| ENSG00000218857.1 | ENSG00000218857 | 4.998340775 | 0.045078 |
| ENSG00000264188.1 | ENSG00000264188 | 4.996302432 | 0.041041 |
| ENSG00000262370.7 | ENSG00000262370 | 4.995813766 | 0.039347 |
| ENSG00000223505.2 | RPL35AP5 | 4.994978634 | 0.040879 |
| ENSG00000130653.16 | PNPLA7 | 4.993725557 | 0.039334 |
| ENSG00000160224.17 | AIRE | 4.993064542 | 0.040646 |
| ENSG00000234566.1 | RPL7AP71 | 4.989799072 | 0.046302 |
| ENSG00000180875.5 | GREM2 | 4.989164148 | 0.041899 |
| ENSG00000147443.13 | DOK2 | 4.988894656 | 0.040367 |
| ENSG00000238057.10 | ZEB2-AS1 | 4.987479355 | 0.043403 |
| ENSG00000254066.2 | LINC01938 | 4.986197858 | 0.04 |
| ENSG00000228718.2 | LINC02521 | 4.978613344 | 0.041508 |
| ENSG00000233170.4 | ENSG00000233170 | 4.975766276 | 0.040292 |
| ENSG00000258315.5 | C17orf49 | 4.972300616 | 0.041118 |
| ENSG00000267649.1 | ENSG00000267649 | 4.968157516 | 0.040711 |
| ENSG00000227331.1 | RPL7AP22 | 4.965886045 | 0.043368 |
| ENSG00000270620.1 | ENSG00000270620 | 4.96273135 | 0.04288 |
| ENSG00000276570.1 | ENSG00000276570 | 4.96267478 | 0.042096 |
| ENSG00000226352.2 | PSPC1-AS2 | 4.962232683 | 0.043708 |
| ENSG00000272420.1 | ENSG00000272420 | 4.960510101 | 0.041248 |
| ENSG00000199990.1 | VTRNA1-1 | 4.960315929 | 0.04879 |
| ENSG00000262833.1 | ENSG00000262833 | 4.955338465 | 0.044202 |
| ENSG00000232373.2 | MTCYBP3 | 4.952607325 | 0.043944 |

| | | | |
|---|---|---|---|
| ENSG00000289534.1 | ENSG00000289534 | 4.952607325 | 0.043944 |
| ENSG00000197177.16 | ADGRA1 | 4.950034857 | 0.041943 |
| ENSG00000100399.16 | CHADL | 4.949836165 | 0.043942 |
| ENSG00000229294.1 | ENSG00000229294 | 4.948637988 | 0.042087 |
| ENSG00000149798.5 | CDC42EP2 | 4.947640139 | 0.045144 |
| ENSG00000104879.5 | CKM | 4.947139139 | 0.042719 |
| ENSG00000223732.2 | ENSG00000223732 | 4.945053807 | 0.042716 |
| ENSG00000265702.2 | ENSG00000265702 | 4.943664262 | 0.042655 |
| ENSG00000234268.1 | ENSG00000234268 | 4.938601973 | 0.045911 |
| ENSG00000107447.8 | DNTT | 4.935748151 | 0.042784 |
| ENSG00000280953.2 | LINC01163 | 4.935322722 | 0.04627 |
| ENSG00000158315.11 | RHBDL2 | 4.935030816 | 0.04514 |
| ENSG00000260744.1 | ENSG00000260744 | 4.934338318 | 0.042793 |
| ENSG00000282876.1 | ENSG00000282876 | 4.931522834 | 0.043122 |
| ENSG00000291187.1 | LINC00032 | 4.930685834 | 0.045016 |
| ENSG00000242060.1 | RPS3AP49 | 4.92222932 | 0.044966 |
| ENSG00000248664.3 | ENSG00000248664 | 4.918634273 | 0.043973 |
| ENSG00000287742.1 | ENSG00000287742 | 4.917074461 | 0.044269 |
| ENSG00000186301.8 | MST1P2 | 4.91590258 | 0.048378 |
| ENSG00000108381.11 | ASPA | 4.913347031 | 0.047947 |
| ENSG00000259581.3 | TYRO3P | 4.912914537 | 0.044602 |
| ENSG00000225928.2 | CACYBPP1 | 4.9114669 | 0.045006 |
| ENSG00000259719.6 | LINC02284 | 4.910010014 | 0.045522 |
| ENSG00000282834.1 | ENSG00000282834 | 4.909271378 | 0.044756 |
| ENSG00000288568.1 | ZCCHC14-DT | 4.909271378 | 0.044756 |
| ENSG00000257835.2 | ENSG00000257835 | 4.908447979 | 0.046461 |
| ENSG00000185775.10 | SPATA31A6 | 4.908341412 | 0.046878 |
| ENSG00000215482.3 | CALM2P3 | 4.90552397 | 0.046133 |
| ENSG00000239282.8 | CASTOR1 | 4.90552397 | 0.046133 |
| ENSG00000284773.1 | ENSG00000284773 | 4.90268623 | 0.045756 |
| ENSG00000278514.1 | ENSG00000278514 | 4.90115073 | 0.044987 |
| ENSG00000219773.1 | RPSAP45 | 4.899708148 | 0.045052 |
| ENSG00000270497.1 | ENSG00000270497 | 4.899708148 | 0.045052 |
| ENSG00000226348.1 | VN2R10P | 4.895388757 | 0.045952 |
| ENSG00000129990.15 | SYT5 | 4.892036223 | 0.049621 |
| ENSG00000286905.1 | ENSG00000286905 | 4.890238403 | 0.045673 |
| ENSG00000118156.13 | ZNF541 | 4.88922548 | 0.048793 |
| ENSG00000159289.6 | GOLGA6A | 4.887353865 | 0.047544 |
| ENSG00000279674.1 | ENSG00000279674 | 4.886452155 | 0.046747 |
| ENSG00000251209.10 | LINC00923 | 4.883500702 | 0.048165 |
| ENSG00000287477.1 | ENSG00000287477 | 4.874559887 | 0.049523 |
| ENSG00000255451.1 | ENSG00000255451 | 4.873271632 | 0.049004 |
| ENSG00000230992.3 | FAM201B | 4.873180863 | 0.048671 |
| ENSG00000185837.5 | HDHD5-AS1 | 4.871665057 | 0.048611 |
| ENSG00000232526.1 | VDAC1P13 | 4.865832134 | 0.047971 |
| ENSG00000237170.3 | RPS7P15 | 4.864355426 | 0.048042 |
| ENSG00000104804.8 | TULP2 | 4.864228081 | 0.047512 |

| | | | |
|---|---|---|---|
| ENSG00000207175.1 | RNU1-67P | 4.864228081 | 0.047512 |
| ENSG00000142606.16 | MMEL1 | 4.861283079 | 0.048169 |
| ENSG00000238966.1 | ENSG00000238966 | 4.856803439 | 0.049836 |
| ENSG00000015592.17 | STMN4 | 4.847740673 | 0.048613 |
| ENSG00000213791.4 | ENSG00000213791 | 4.847740673 | 0.048613 |
| ENSG00000237487.1 | VN1R48P | 4.847740673 | 0.048613 |
| ENSG00000184100.6 | BRD7P2 | 4.844665567 | 0.049839 |
| ENSG00000188801.9 | ZNF322P1 | 4.63316489 | 0.003309 |
| ENSG00000159409.15 | CELF3 | 4.533344802 | 0.004477 |
| ENSG00000254946.2 | LINC02751 | 4.428607348 | 9.15E-05 |
| ENSG00000280145.3 | ENSG00000280145 | 4.427450029 | 9.25E-13 |
| ENSG00000261474.1 | ENSG00000261474 | 4.322496551 | 0.007055 |
| ENSG00000289441.1 | ENSG00000289441 | 4.124315452 | 0.000526 |
| ENSG00000271852.1 | SNORD11B | 4.098904413 | 0.012081 |
| ENSG00000105392.17 | CRX | 4.003884692 | 0.000677 |
| ENSG00000128610.12 | FEZF1 | 3.993445418 | 2.48E-06 |
| ENSG00000270022.3 | ENSG00000270022 | 3.948241618 | 0.016861 |
| ENSG00000100362.13 | PVALB | 3.93177582 | 0.001017 |
| ENSG00000092345.14 | DAZL | 3.803856777 | 2.22E-20 |
| ENSG00000207475.1 | SNORA80E | 3.766359278 | 0.025505 |
| ENSG00000261002.5 | VPS39-DT | 3.683415102 | 0.035535 |
| ENSG00000177468.7 | OLIG3 | 3.662087503 | 3.96E-05 |
| ENSG00000266598.1 | ENSG00000266598 | 3.622952473 | 0.035069 |
| ENSG00000286403.1 | ENSG00000286403 | 3.61198466 | 0.005087 |
| ENSG00000078018.22 | MAP2 | 3.498346872 | 4.66E-55 |
| ENSG00000173825.7 | TIGD3 | 3.497042884 | 0.005081 |
| ENSG00000104332.12 | SFRP1 | 3.494656094 | 2.96E-73 |
| ENSG00000247121.8 | ENSG00000247121 | 3.484795794 | 0.000483 |
| ENSG00000204581.3 | ACOXL-AS1 | 3.472256148 | 0.045259 |
| ENSG00000257543.1 | ENSG00000257543 | 3.435791386 | 0.005503 |
| ENSG00000173673.8 | HES3 | 3.424225291 | 2.2E-12 |
| ENSG00000260314.3 | MRC1 | 3.417057815 | 0.00769 |
| ENSG00000236714.3 | LINC01844 | 3.395185186 | 0.000336 |
| ENSG00000227659.1 | CLYBL-AS2 | 3.353109894 | 0.006568 |
| ENSG00000196544.8 | BORCS6 | 3.330871911 | 0.007537 |
| ENSG00000273253.2 | TRABD-AS1 | 3.30427177 | 0.010784 |
| ENSG00000231728.5 | TMSB15B-AS1 | 3.302545143 | 0.008374 |
| ENSG00000125775.15 | SDCBP2 | 3.287899907 | 0.008571 |
| ENSG00000234945.8 | GTF3C2-AS1 | 3.281617611 | 0.008833 |
| ENSG00000289159.2 | ENSG00000289159 | 3.256484183 | 0.009662 |
| ENSG00000226744.1 | ENSG00000226744 | 3.233094535 | 0.010622 |
| ENSG00000130528.12 | HRC | 3.228646418 | 0.001966 |
| ENSG00000286703.1 | ENSG00000286703 | 3.200513474 | 0.0116 |
| ENSG00000153993.14 | SEMA3D | 3.175446544 | 4.37E-08 |
| ENSG00000286909.1 | ENSG00000286909 | 3.1714726 | 0.013165 |
| ENSG00000289643.2 | ENSG00000289643 | 3.161599061 | 7.96E-07 |
| ENSG00000228168.1 | HNRNPA1P21 | 3.151978702 | 0.000445 |

| | | | |
|---|---|---|---|
| ENSG00000204472.13 | AIF1 | 3.14032799 | 1.11E-07 |
| ENSG00000288049.1 | ENSG00000288049 | 3.14032352 | 0.013266 |
| ENSG00000156219.18 | ART3 | 3.105311303 | 3.1E-13 |
| ENSG00000168477.19 | TNXB | 3.097231564 | 0.000664 |
| ENSG00000228677.1 | TTC3-AS1 | 3.073778511 | 0.018902 |
| ENSG00000289056.2 | ENSG00000289056 | 3.072997018 | 0.017589 |
| ENSG00000105675.9 | ATP4A | 3.070556869 | 0.016806 |
| ENSG00000160180.17 | TFF3 | 3.048782017 | 5.79E-07 |
| ENSG00000198798.6 | MAGEB3 | 3.034574226 | 0.001285 |
| ENSG00000086717.19 | PPEF1 | 3.034571079 | 0.017529 |
| ENSG00000099338.23 | CATSPERG | 3.031669452 | 0.020131 |
| ENSG00000206899.1 | RNU6-36P | 3.028776231 | 0.018284 |
| ENSG00000232970.1 | POLHP1 | 3.027363797 | 0.018359 |
| ENSG00000002822.16 | MAD1L1 | 3.020985833 | 0.023989 |
| ENSG00000261997.1 | ENSG00000261997 | 3.017917915 | 0.019506 |
| ENSG00000280378.1 | ENSG00000280378 | 3.015072731 | 0.020138 |
| ENSG00000254648.1 | ENSG00000254648 | 3.008115566 | 0.020035 |
| ENSG00000283692.2 | ENSG00000283692 | 3.007462019 | 0.018828 |
| ENSG00000181195.11 | PENK | 3.003526244 | 0.021935 |
| ENSG00000248550.4 | OTX2-AS1 | 2.997040365 | 0.019316 |
| ENSG00000169218.14 | RSPO1 | 2.968347681 | 0.002688 |
| ENSG00000285569.4 | ENSG00000285569 | 2.958108719 | 0.021541 |
| ENSG00000265972.6 | TXNIP | 2.957837333 | 9.69E-94 |
| ENSG00000128606.13 | LRRC17 | 2.939088407 | 0.001556 |
| ENSG00000237921.2 | ENSG00000237921 | 2.936686621 | 0.024072 |
| ENSG00000160471.13 | COX6B2 | 2.931762114 | 8.04E-07 |
| ENSG00000196660.12 | SLC30A10 | 2.922703166 | 0.024208 |
| ENSG00000235945.1 | ENSG00000235945 | 2.916646474 | 0.025412 |
| ENSG00000249853.9 | HS3ST5 | 2.900081961 | 0.002221 |
| ENSG00000279943.1 | FLJ38576 | 2.888312739 | 0.028444 |
| ENSG00000287886.1 | ENSG00000287886 | 2.880156295 | 0.033867 |
| ENSG00000164049.14 | FBXW12 | 2.87454499 | 0.027945 |
| ENSG00000264575.1 | LINC00526 | 2.868781632 | 0.000206 |
| ENSG00000271964.1 | ENSG00000271964 | 2.863693738 | 0.029039 |
| ENSG00000272662.1 | TIMMDC1-DT | 2.861952337 | 0.028896 |
| ENSG00000252186.1 | RNU6-781P | 2.858718649 | 0.030262 |
| ENSG00000247228.3 | ENSG00000247228 | 2.856681092 | 0.032743 |
| ENSG00000260572.1 | PPM1L-DT | 2.835878229 | 0.031503 |
| ENSG00000099840.14 | IZUMO4 | 2.824997889 | 0.032883 |
| ENSG00000081248.12 | CACNA1S | 2.824473114 | 0.011144 |
| ENSG00000240509.1 | RPL34P18 | 2.820469797 | 0.031054 |
| ENSG00000111305.19 | GSG1 | 2.810296038 | 3.37E-08 |
| ENSG00000258732.1 | ENSG00000258732 | 2.785845154 | 0.036612 |
| ENSG00000285427.1 | SOD2-OT1 | 2.785528207 | 0.009775 |
| ENSG00000114654.8 | EFCC1 | 2.773538981 | 0.036117 |
| ENSG00000250397.2 | ENSG00000250397 | 2.762572564 | 0.042488 |
| ENSG00000234338.1 | ENSG00000234338 | 2.760903721 | 0.037439 |

| | | | |
|---|---|---|---|
| ENSG00000161940.11 | BCL6B | 2.757293197 | 0.003307 |
| ENSG00000285530.1 | ENSG00000285530 | 2.750891785 | 0.038646 |
| ENSG00000182968.5 | SOX1 | 2.746527083 | 0.037836 |
| ENSG00000279735.2 | ENSG00000279735 | 2.74212627 | 0.0438 |
| ENSG00000124479.11 | NDP | 2.731913208 | 0.00042 |
| ENSG00000183032.12 | SLC25A21 | 2.723072744 | 8.71E-05 |
| ENSG00000261121.2 | LINC02473 | 2.722562316 | 0.005342 |
| ENSG00000223866.1 | ENSG00000223866 | 2.717078121 | 0.043026 |
| ENSG00000230316.8 | FEZF1-AS1 | 2.710341334 | 0.006919 |
| ENSG00000005379.17 | TSPOAP1 | 2.705101514 | 0.004287 |
| ENSG00000067842.19 | ATP2B3 | 2.679606558 | 0.000934 |
| ENSG00000288868.2 | ENSG00000288868 | 2.67713594 | 0.004875 |
| ENSG00000201900.1 | RNY1P13 | 2.671930471 | 0.047334 |
| ENSG00000273540.5 | AGBL1 | 2.670353508 | 0.048065 |
| ENSG00000092850.12 | TEKT2 | 2.66250386 | 0.000631 |
| ENSG00000230246.8 | SPATA31C1 | 2.662423777 | 4.5E-06 |
| ENSG00000237194.1 | SNAI1P1 | 2.6620434 | 0.049423 |
| ENSG00000178429.9 | RPS3AP5 | 2.654945073 | 0.006094 |
| ENSG00000229271.3 | ENSG00000229271 | 2.653474278 | 8.02E-21 |
| ENSG00000101883.6 | RHOXF1 | 2.649752354 | 0.048603 |
| ENSG00000213997.3 | PGAM1P7 | 2.645675923 | 0.000427 |
| ENSG00000234369.1 | TATDN1P1 | 2.634450262 | 0.006596 |
| ENSG00000172247.4 | C1QTNF4 | 2.620842131 | 9.46E-07 |
| ENSG00000260278.1 | BMP2K-DT | 2.612964146 | 0.008348 |
| ENSG00000124610.5 | H1-1 | 2.584024453 | 1.83E-07 |
| ENSG00000164933.12 | SLC25A32 | 2.578150584 | 0.020271 |
| ENSG00000251143.1 | ENSG00000251143 | 2.570714124 | 0.009129 |
| ENSG00000287516.1 | ENSG00000287516 | 2.568678019 | 0.007801 |
| ENSG00000177910.9 | SPATA31C2 | 2.527651283 | 8.63E-09 |
| ENSG00000100078.4 | PLA2G3 | 2.508687961 | 3.64E-11 |
| ENSG00000288955.1 | ENSG00000288955 | 2.483594029 | 0.01143 |
| ENSG00000231412.3 | ENSG00000231412 | 2.468538882 | 0.000467 |
| ENSG00000063015.21 | SEZ6 | 2.461626243 | 3.03E-05 |
| ENSG00000130711.5 | PRDM12 | 2.459993371 | 0.029317 |
| ENSG00000174776.12 | WDR49 | 2.452196952 | 3.44E-11 |
| ENSG00000146453.13 | PNLDC1 | 2.420583794 | 0.004656 |
| ENSG00000136110.13 | CNMD | 2.420185992 | 4.45E-06 |
| ENSG00000279970.1 | ENSG00000279970 | 2.41890404 | 0.007204 |
| ENSG00000286171.1 | ENSG00000286171 | 2.414292494 | 0.026991 |
| ENSG00000174898.16 | CATSPERD | 2.40112457 | 0.035342 |
| ENSG00000130943.7 | PKDREJ | 2.392048613 | 0.000181 |
| ENSG00000075884.14 | ARHGAP15 | 2.387578343 | 0.015579 |
| ENSG00000197993.9 | KEL | 2.37965509 | 0.003353 |
| ENSG00000237972.1 | TUBG1P | 2.364901963 | 0.000152 |
| ENSG00000161905.13 | ALOX15 | 2.360629838 | 2.46E-05 |
| ENSG00000174527.11 | MYO1H | 2.353487906 | 0.020265 |
| ENSG00000273209.1 | ENSG00000273209 | 2.345032822 | 0.01931 |

| | | | |
|---|---|---|---|
| ENSG00000237187.9 | NR2F1-AS1 | 2.342969081 | 0.000301 |
| ENSG00000170927.15 | PKHD1 | 2.340925263 | 0.021952 |
| ENSG00000249502.3 | MED28-DT | 2.328568314 | 0.019947 |
| ENSG00000113100.10 | CDH9 | 2.315432445 | 1.39E-14 |
| ENSG00000186510.12 | CLCNKA | 2.310271596 | 0.021519 |
| ENSG00000286370.1 | ENSG00000286370 | 2.309302385 | 0.003763 |
| ENSG00000107242.20 | PIP5K1B | 2.30598182 | 1.97E-12 |
| ENSG00000188770.10 | OPTC | 2.30412382 | 0.000102 |
| ENSG00000267270.8 | PARD6G-AS1 | 2.302364255 | 0.004985 |
| ENSG00000198822.11 | GRM3 | 2.300602487 | 0.006349 |
| ENSG00000243230.1 | ENSG00000243230 | 2.299594687 | 0.022007 |
| ENSG00000196542.9 | SPTSSB | 2.299130647 | 2.15E-14 |
| ENSG00000235121.1 | ENSG00000235121 | 2.297324192 | 0.032338 |
| ENSG00000289324.2 | ENSG00000289324 | 2.28833365 | 0.024232 |
| ENSG00000228271.2 | LINC02048 | 2.286405265 | 0.025923 |
| ENSG00000160179.19 | ABCG1 | 2.280379004 | 3.41E-05 |
| ENSG00000231646.6 | FSIP2-AS1 | 2.273961585 | 0.007978 |
| ENSG00000140450.9 | ARRDC4 | 2.269657128 | 7.07E-28 |
| ENSG00000092051.18 | JPH4 | 2.269194519 | 1.07E-06 |
| ENSG00000183908.7 | LRRC55 | 2.268178062 | 4.07E-14 |
| ENSG00000289543.2 | ENSG00000289543 | 2.262505874 | 0.006138 |
| ENSG00000144406.19 | UNC80 | 2.259337901 | 8.42E-18 |
| ENSG00000260455.2 | NBAT1 | 2.256403019 | 0.006531 |
| ENSG00000290917.1 | RAET1K | 2.249242106 | 0.007021 |
| ENSG00000200959.1 | SNORA74A | 2.243256623 | 0.005619 |
| ENSG00000214796.10 | TUBA5P | 2.235635726 | 0.026952 |
| ENSG00000252826.1 | RNVU1-23 | 2.224728558 | 0.029978 |
| ENSG00000260774.1 | ENSG00000260774 | 2.21543583 | 0.029853 |
| ENSG00000204815.10 | ODAD4 | 2.2085497 | 0.009505 |
| ENSG00000124772.12 | CPNE5 | 2.207339844 | 2.08E-05 |
| ENSG00000278238.2 | CKAP2-DT | 2.205212654 | 0.003539 |
| ENSG00000267053.9 | ENSG00000267053 | 2.203801712 | 0.00094 |
| ENSG00000229046.1 | HMGN1P2 | 2.193194511 | 0.036 |
| ENSG00000200169.1 | RNU5D-1 | 2.192377619 | 4.14E-10 |
| ENSG00000184434.8 | LRRC19 | 2.182922691 | 0.000555 |
| ENSG00000237923.2 | LINC02570 | 2.168004634 | 0.00356 |
| ENSG00000242797.3 | GLYCTK-AS1 | 2.161170305 | 0.006204 |
| ENSG00000277130.1 | ENSG00000277130 | 2.156448399 | 0.014135 |
| ENSG00000186910.4 | SERPINA11 | 2.151330595 | 0.042053 |
| ENSG00000129159.9 | KCNC1 | 2.146306341 | 0.000227 |
| ENSG00000111834.13 | RSPH4A | 2.144882806 | 0.041921 |
| ENSG00000259577.2 | CERNA1 | 2.142429662 | 0.013566 |
| ENSG00000215808.4 | LINC01139 | 2.13831548 | 1.06E-05 |
| ENSG00000269289.6 | ENSG00000269289 | 2.126866423 | 0.011555 |
| ENSG00000276255.2 | LINC02809 | 2.126630957 | 0.044172 |
| ENSG00000129654.8 | FOXJ1 | 2.122434092 | 0.041752 |
| ENSG00000167011.9 | NAT16 | 2.119804524 | 0.011909 |

| | | | |
|---|---|---|---|
| ENSG00000135638.14 | EMX1 | 2.119607921 | 0.001305 |
| ENSG00000102290.23 | PCDH11X | 2.119127917 | 1.04E-09 |
| ENSG00000249379.2 | ENSG00000249379 | 2.112889064 | 0.042937 |
| ENSG00000135472.9 | FAIM2 | 2.1115527 | 0.042197 |
| ENSG00000015413.10 | DPEP1 | 2.109329153 | 0.002235 |
| ENSG00000136250.12 | AOAH | 2.106016021 | 1.18E-05 |
| ENSG00000130055.14 | GDPD2 | 2.105399716 | 5.21E-16 |
| ENSG00000158816.16 | VWA5B1 | 2.098855875 | 0.016482 |
| ENSG00000285838.2 | ENSG00000285838 | 2.097523888 | 3.49E-06 |
| ENSG00000184408.10 | KCND2 | 2.096040505 | 9.06E-11 |
| ENSG00000178297.15 | TMPRSS9 | 2.092091606 | 0.033494 |
| ENSG00000231160.12 | KLF3-AS1 | 2.081629365 | 1.92E-05 |
| ENSG00000259775.1 | EIF5-DT | 2.081493272 | 0.025983 |
| ENSG00000257433.8 | RPAP3-DT | 2.081207119 | 0.046033 |
| ENSG00000108823.17 | SGCA | 2.075995425 | 0.045896 |
| ENSG00000112599.9 | GUCA1B | 2.062087339 | 0.048577 |
| ENSG00000144339.12 | TMEFF2 | 2.06065902 | 0.002471 |
| ENSG00000286705.2 | ENSG00000286705 | 2.044185554 | 0.009826 |
| ENSG00000149599.16 | DUSP15 | 2.031997375 | 0.006774 |
| ENSG00000111913.20 | RIPOR2 | 2.029215076 | 6.97E-09 |
| ENSG00000254480.2 | LINC02749 | 2.025975908 | 1.93E-05 |
| ENSG00000212402.1 | SNORA74B | 2.021402293 | 0.000105 |
| ENSG00000126010.6 | GRPR | 2.02018877 | 1.53E-17 |
| ENSG00000161860.8 | SYCE2 | 2.019726692 | 4.45E-10 |
| ENSG00000258096.1 | SLC38A2-AS1 | 2.016056589 | 0.03406 |
| ENSG00000285820.1 | ENSG00000285820 | 2.014288831 | 0.000558 |
| ENSG00000276603.1 | ENSG00000276603 | 2.009620069 | 0.022668 |
| ENSG00000115138.11 | POMC | 2.00757854 | 0.007101 |

**Downregulated genes**

| Gene | Gene symbol | log2FoldChange | pvalue |
|---|---|---|---|
| ENSG00000213218.11 | CSH2 | -7.685117114 | 0.000106 |
| ENSG00000273233.1 | ENSG00000273233 | -7.25045466 | 4.47E-25 |
| ENSG00000280362.1 | ENSG00000280362 | -6.315597969 | 0.000155 |
| ENSG00000132965.10 | ALOX5AP | -6.270048265 | 0.030156 |
| ENSG00000134827.8 | TCN1 | -6.270048265 | 0.030156 |
| ENSG00000150201.15 | FXYD4 | -6.270048265 | 0.030156 |
| ENSG00000228742.11 | LINC02577 | -6.270048265 | 0.030156 |
| ENSG00000229380.1 | PRKAR1B-AS2 | -6.270048265 | 0.030156 |
| ENSG00000230039.1 | CCNYL5 | -6.270048265 | 0.030156 |
| ENSG00000242435.1 | UPK3BP1 | -6.270048265 | 0.030156 |
| ENSG00000252291.1 | ENSG00000252291 | -6.270048265 | 0.030156 |
| ENSG00000259354.5 | ENSG00000259354 | -6.270048265 | 0.030156 |
| ENSG00000266179.2 | ENSG00000266179 | -6.270048265 | 0.030156 |
| ENSG00000277598.1 | ENSG00000277598 | -6.270048265 | 0.030156 |
| ENSG00000283755.2 | CPHXL | -6.270048265 | 0.030156 |

| | | | |
|---|---|---|---|
| ENSG00000287622.1 | ENSG00000287622 | -6.270048265 | 0.030156 |
| ENSG00000288172.1 | ENSG00000288172 | -6.270048265 | 0.030156 |
| ENSG00000242307.1 | RPS26P52 | -6.216430599 | 4.28E-06 |
| ENSG00000104237.11 | RP1 | -5.84999589 | 1.05E-23 |
| ENSG00000037965.6 | HOXC8 | -5.695763257 | 0.003192 |
| ENSG00000254851.1 | ENSG00000254851 | -5.591095778 | 0.004761 |
| ENSG00000177257.3 | DEFB4B | -5.450272388 | 0.007353 |
| ENSG00000128510.12 | CPA4 | -5.338955047 | 0.000224 |
| ENSG00000278075.1 | ENSG00000278075 | -5.308268855 | 0.047902 |
| ENSG00000197646.8 | PDCD1LG2 | -5.220722286 | 0.01612 |
| ENSG00000204577.12 | LILRB3 | -5.220722286 | 0.01612 |
| ENSG00000166396.13 | SERPINB7 | -5.194514836 | 1.91E-08 |
| ENSG00000226629.1 | LINC00974 | -5.054059879 | 0.001098 |
| ENSG00000232810.5 | TNF | -5.038341002 | 0.019519 |
| ENSG00000130997.16 | POLN | -4.905793321 | 0.036788 |
| ENSG00000137473.19 | TTC29 | -4.905793321 | 0.036788 |
| ENSG00000170290.4 | SLN | -4.905793321 | 0.036788 |
| ENSG00000228386.2 | ENSG00000228386 | -4.905793321 | 0.036788 |
| ENSG00000260492.3 | CISTR | -4.905793321 | 0.036788 |
| ENSG00000267052.3 | ENSG00000267052 | -4.905793321 | 0.036788 |
| ENSG00000249641.2 | HOXC13-AS | -4.883237635 | 0.000306 |
| ENSG00000280025.1 | ENSG00000280025 | -4.872104436 | 0.037851 |
| ENSG00000286607.1 | ENSG00000286607 | -4.872104436 | 0.037851 |
| ENSG00000248455.8 | LINC02217 | -4.793830188 | 7.94E-05 |
| ENSG00000145888.11 | GLRA1 | -4.703379276 | 0.043894 |
| ENSG00000169429.12 | CXCL8 | -4.702202488 | 1.49E-09 |
| ENSG00000120937.9 | NPPB | -4.610100577 | 4.47E-22 |
| ENSG00000137674.4 | MMP20 | -4.564572199 | 0.000723 |
| ENSG00000168542.16 | COL3A1 | -4.509686643 | 5.1E-12 |
| ENSG00000230156.3 | LINC00443 | -4.502083722 | 0.039614 |
| ENSG00000162892.16 | IL24 | -4.493403611 | 0.00491 |
| ENSG00000243709.1 | LEFTY1 | -4.457015995 | 0.000362 |
| ENSG00000174145.8 | NWD2 | -4.368686563 | 0.014379 |
| ENSG00000150551.11 | LYPD1 | -4.312141875 | 1.03E-19 |
| ENSG00000134339.9 | SAA2 | -4.246555341 | 0.040978 |
| ENSG00000232530.1 | LIF-AS1 | -4.219509667 | 0.026323 |
| ENSG00000286433.1 | ENSG00000286433 | -4.165353467 | 0.049957 |
| ENSG00000196091.15 | MYBPC1 | -4.102092387 | 0.030889 |
| ENSG00000049249.10 | TNFRSF9 | -4.097410017 | 1.83E-10 |
| ENSG00000167772.12 | ANGPTL4 | -4.036838036 | 4.08E-13 |
| ENSG00000280623.1 | PCAT14 | -3.993295858 | 7.52E-23 |
| ENSG00000233532.6 | LINC00460 | -3.958539448 | 0.003408 |
| ENSG00000229847.9 | EMX2OS | -3.948130176 | 0.009505 |
| ENSG00000268964.3 | ERVV-2 | -3.905011633 | 0.009775 |

| | | | |
|---|---|---|---|
| ENSG00000148677.7 | ANKRD1 | -3.860327534 | 0.000128 |
| ENSG00000236355.2 | DNAH8-DT | -3.855017516 | 0.027939 |
| ENSG00000204941.14 | PSG5 | -3.825518107 | 0.016591 |
| ENSG00000246640.2 | PICART1 | -3.737783113 | 0.031608 |
| ENSG00000156265.16 | MAP3K7CL | -3.709632727 | 2.57E-10 |
| ENSG00000225383.8 | SFTA1P | -3.703532189 | 5.66E-14 |
| ENSG00000137090.12 | DMRT1 | -3.688009429 | 0.000127 |
| ENSG00000115648.14 | MLPH | -3.664891045 | 1.4E-05 |
| ENSG00000240244.3 | GAPDHP33 | -3.656831099 | 0.012717 |
| ENSG00000147883.12 | CDKN2B | -3.591142447 | 1.57E-07 |
| ENSG00000166741.8 | NNMT | -3.585313623 | 4.99E-09 |
| ENSG00000180251.5 | SLC9A4 | -3.533152576 | 0.000778 |
| ENSG00000188015.10 | S100A3 | -3.531247337 | 1.2E-09 |
| ENSG00000272209.1 | ENSG00000272209 | -3.508643073 | 0.039926 |
| ENSG00000105991.10 | HOXA1 | -3.483052337 | 0.002736 |
| ENSG00000105383.15 | CD33 | -3.479796808 | 8.35E-09 |
| ENSG00000087494.16 | PTHLH | -3.43217438 | 0.000462 |
| ENSG00000172716.17 | SLFN11 | -3.386737567 | 0.000829 |
| ENSG00000155897.10 | ADCY8 | -3.376145552 | 0.044645 |
| ENSG00000124785.9 | NRN1 | -3.368547614 | 0.029721 |
| ENSG00000197632.9 | SERPINB2 | -3.353601864 | 0.046411 |
| ENSG00000258404.1 | LINC02320 | -3.349824709 | 0.013205 |
| ENSG00000165617.15 | DACT1 | -3.346365294 | 1.01E-38 |
| ENSG00000163739.5 | CXCL1 | -3.324878914 | 1.71E-20 |
| ENSG00000117148.8 | ACTL8 | -3.283768423 | 8.22E-10 |
| ENSG00000290418.1 | TPTEP1 | -3.280131405 | 0.048234 |
| ENSG00000106366.9 | SERPINE1 | -3.238052143 | 1.06E-07 |
| ENSG00000243137.8 | PSG4 | -3.205721281 | 0.01401 |
| ENSG00000288781.2 | ENSG00000288781 | -3.20030008 | 0.004695 |
| ENSG00000289533.1 | ENSG00000289533 | -3.141128634 | 2.89E-32 |
| ENSG00000237827.1 | RPS15AP29 | -3.134165717 | 0.046337 |
| ENSG00000173391.9 | OLR1 | -3.129874433 | 2.58E-08 |
| ENSG00000175077.6 | RTP1 | -3.098999662 | 0.000571 |
| ENSG00000044524.11 | EPHA3 | -3.092757683 | 0.000386 |
| ENSG00000171860.5 | C3AR1 | -3.073807959 | 0.039166 |
| ENSG00000124143.11 | ARHGAP40 | -3.065952685 | 0.000162 |
| ENSG00000287492.1 | ENSG00000287492 | -3.062157304 | 4.09E-11 |
| ENSG00000164112.14 | SMIM43 | -3.039142662 | 0.000695 |
| ENSG00000087589.17 | CASS4 | -3.032294935 | 0.009303 |
| ENSG00000138772.13 | ANXA3 | -3.025003521 | 3.74E-27 |
| ENSG00000164509.16 | IL31RA | -2.995311194 | 0.043284 |
| ENSG00000236975.2 | LINC02814 | -2.976447187 | 9.2E-07 |
| ENSG00000286069.2 | ENSG00000286069 | -2.974905875 | 0.000533 |
| ENSG00000175592.9 | FOSL1 | -2.971746286 | 1.47E-48 |

| | | | |
|---|---|---|---|
| ENSG00000230838.2 | LINC01614 | -2.95361939 | 0.00069 |
| ENSG00000069011.16 | PITX1 | -2.953540363 | 4.22E-05 |
| ENSG00000166317.12 | SYNPO2L | -2.950847466 | 0.002743 |
| ENSG00000078401.7 | EDN1 | -2.94768697 | 3.96E-09 |
| ENSG00000229953.1 | ENSG00000229953 | -2.939625286 | 1.88E-06 |
| ENSG00000248441.9 | LETR1 | -2.92754472 | 0.014782 |
| ENSG00000120217.14 | CD274 | -2.919713925 | 0.001129 |
| ENSG00000244513.7 | EOGT-DT | -2.91786075 | 1.47E-13 |
| ENSG00000110090.13 | CPT1A | -2.906767197 | 2.26E-58 |
| ENSG00000188984.12 | AADACL3 | -2.890158698 | 1.74E-12 |
| ENSG00000122223.13 | CD244 | -2.845268321 | 0.022422 |
| ENSG00000143995.20 | MEIS1 | -2.82952832 | 4.22E-10 |
| ENSG00000286595.1 | ENSG00000286595 | -2.824378156 | 0.042159 |
| ENSG00000074706.14 | IPCEF1 | -2.815646723 | 0.01214 |
| ENSG00000237523.2 | LINC00857 | -2.797091541 | 0.000938 |
| ENSG00000138347.17 | MYPN | -2.779217456 | 0.011844 |
| ENSG00000211448.14 | DIO2 | -2.760249954 | 0.004858 |
| ENSG00000254428.1 | ENSG00000254428 | -2.7486035 | 0.011493 |
| ENSG00000134531.10 | EMP1 | -2.732205072 | 8.32E-18 |
| ENSG00000073756.13 | PTGS2 | -2.699488373 | 1.4E-05 |
| ENSG00000163453.11 | IGFBP7 | -2.694938089 | 9.39E-39 |
| ENSG00000166750.10 | SLFN5 | -2.689489401 | 3.71E-13 |
| ENSG00000118785.15 | SPP1 | -2.67434174 | 1.04E-56 |
| ENSG00000253522.6 | MIR3142HG | -2.667293502 | 0.015258 |
| ENSG00000244300.3 | GATA2-AS1 | -2.657872233 | 0.001132 |
| ENSG00000134107.5 | BHLHE40 | -2.651525653 | 1.33E-31 |
| ENSG00000260941.1 | LINC00622 | -2.636131861 | 0.00015 |
| ENSG00000239093.1 | ENSG00000239093 | -2.627701813 | 0.018472 |
| ENSG00000224063.6 | CALCRL-AS1 | -2.570156431 | 8.53E-07 |
| ENSG00000107485.18 | GATA3 | -2.563904916 | 1.27E-12 |
| ENSG00000113492.14 | AGXT2 | -2.559486602 | 0.037215 |
| ENSG00000159176.14 | CSRP1 | -2.548317126 | 7.67E-15 |
| ENSG00000188404.10 | SELL | -2.5147465 | 0.00337 |
| ENSG00000219445.2 | ENSG00000219445 | -2.489306397 | 0.000112 |
| ENSG00000183625.16 | CCR3 | -2.483268417 | 0.0076 |
| ENSG00000113083.15 | LOX | -2.461990805 | 1.64E-32 |
| ENSG00000168334.9 | XIRP1 | -2.460095479 | 3.97E-05 |
| ENSG00000169594.14 | BNC1 | -2.457543742 | 1.2E-13 |
| ENSG00000268696.3 | ZNF723 | -2.45244702 | 0.002237 |
| ENSG00000170961.7 | HAS2 | -2.446217556 | 1.78E-15 |
| ENSG00000054356.14 | PTPRN | -2.436902101 | 6.19E-11 |
| ENSG00000289003.1 | ENSG00000289003 | -2.436597743 | 0.029036 |
| ENSG00000123572.17 | NRK | -2.433984866 | 1E-07 |
| ENSG00000141448.11 | GATA6 | -2.420291506 | 0.003161 |

| | | | |
|---|---|---|---|
| ENSG00000128040.12 | SPINK2 | -2.416450387 | 0.005516 |
| ENSG00000085276.19 | MECOM | -2.411996054 | 0.000419 |
| ENSG00000184984.10 | CHRM5 | -2.399254005 | 0.034912 |
| ENSG00000266920.1 | ACTBP9 | -2.390397682 | 0.033959 |
| ENSG00000286153.2 | ENSG00000286153 | -2.385267015 | 0.012605 |
| ENSG00000177409.12 | SAMD9L | -2.384287462 | 0.043933 |
| ENSG00000154342.6 | WNT3A | -2.363009333 | 0.010264 |
| ENSG00000168621.15 | GDNF | -2.356712297 | 0.022056 |
| ENSG00000173401.10 | GLIPR1L1 | -2.338472371 | 0.004175 |
| ENSG00000254703.4 | SENCR | -2.324939095 | 0.000229 |
| ENSG00000272068.1 | BCAN-AS1 | -2.31913844 | 0.024086 |
| ENSG00000198729.5 | PPP1R14C | -2.308397208 | 0.02129 |
| ENSG00000135046.14 | ANXA1 | -2.303234516 | 2.19E-06 |
| ENSG00000271524.1 | BNIP3P17 | -2.302286183 | 0.000119 |
| ENSG00000166401.15 | SERPINB8 | -2.288753761 | 6.05E-05 |
| ENSG00000145824.13 | CXCL14 | -2.283896385 | 3.89E-21 |
| ENSG00000164093.18 | PITX2 | -2.268750905 | 0.01581 |
| ENSG00000125148.7 | MT2A | -2.264460239 | 5.71E-06 |
| ENSG00000023445.16 | BIRC3 | -2.257094263 | 0.001276 |
| ENSG00000162849.16 | KIF26B | -2.255058259 | 0.000539 |
| ENSG00000122641.11 | INHBA | -2.247804467 | 0.000524 |
| ENSG00000173432.13 | SAA1 | -2.240049461 | 0.031764 |
| ENSG00000280079.1 | ENSG00000280079 | -2.235964709 | 7.76E-11 |
| ENSG00000013588.9 | GPRC5A | -2.23548735 | 2.3E-14 |
| ENSG00000079263.19 | SP140 | -2.232427808 | 0.000276 |
| ENSG00000272405.1 | ENSG00000272405 | -2.228282449 | 0.000402 |
| ENSG00000232815.1 | DUX4L50 | -2.227719216 | 0.008008 |
| ENSG00000102683.8 | SGCG | -2.218268904 | 0.003406 |
| ENSG00000128422.18 | KRT17 | -2.212085329 | 0.009277 |
| ENSG00000187510.11 | PLEKHG7 | -2.204809443 | 0.02737 |
| ENSG00000179348.13 | GATA2 | -2.202007701 | 3.98E-11 |
| ENSG00000204335.4 | SP5 | -2.195701157 | 1.53E-06 |
| ENSG00000287453.1 | ENSG00000287453 | -2.195013057 | 0.040736 |
| ENSG00000248538.10 | PPP1R3B-DT | -2.18829871 | 2.26E-08 |
| ENSG00000206262.9 | FOXL2NB | -2.171591781 | 4.09E-05 |
| ENSG00000086205.18 | FOLH1 | -2.161369308 | 0.002283 |
| ENSG00000116016.14 | EPAS1 | -2.157302569 | 1.51E-13 |
| ENSG00000008517.19 | IL32 | -2.153386368 | 0.000276 |
| ENSG00000270890.1 | ENSG00000270890 | -2.151504602 | 0.044095 |
| ENSG00000101276.18 | SLC52A3 | -2.151147924 | 1.37E-09 |
| ENSG00000071203.9 | MS4A12 | -2.143738094 | 0.028778 |
| ENSG00000092969.12 | TGFB2 | -2.139600956 | 4.21E-09 |
| ENSG00000169884.14 | WNT10B | -2.132833783 | 2.17E-06 |
| ENSG00000170382.12 | LRRN2 | -2.109660842 | 4.54E-19 |

| | | | |
|---|---|---|---|
| ENSG00000136869.16 | TLR4 | -2.10633814 | 9.73E-05 |
| ENSG00000120149.9 | MSX2 | -2.097340821 | 6.24E-23 |
| ENSG00000261060.1 | ENSG00000261060 | -2.092616532 | 0.045374 |
| ENSG00000135218.19 | CD36 | -2.092418838 | 0.04108 |
| ENSG00000225555.2 | ENSG00000225555 | -2.090787901 | 0.003112 |
| ENSG00000172548.15 | NIPAL4 | -2.083179339 | 0.004337 |
| ENSG00000203780.12 | FANK1 | -2.077812218 | 0.005388 |
| ENSG00000204525.18 | HLA-C | -2.065686317 | 3.35E-17 |
| ENSG00000119283.16 | TRIM67 | -2.064500466 | 0.042078 |
| ENSG00000185269.12 | NOTUM | -2.054479265 | 0.002056 |
| ENSG00000143878.10 | RHOB | -2.052216643 | 2.83E-37 |
| ENSG00000213981.8 | ENSG00000213981 | -2.046509521 | 0.009439 |
| ENSG00000242574.9 | HLA-DMB | -2.0394277 | 0.03391 |
| ENSG00000136244.12 | IL6 | -2.032315271 | 2.05E-05 |
| ENSG00000117322.19 | CR2 | -2.031404952 | 4.43E-07 |
| ENSG00000233056.4 | ERVH48-1 | -2.031206101 | 5.78E-20 |
| ENSG00000240583.14 | AQP1 | -2.03008641 | 0.003841 |
| ENSG00000223715.3 | LINC01208 | -2.023738391 | 0.024542 |
| ENSG00000261175.6 | LINC02188 | -2.022290289 | 9.41E-05 |
| ENSG00000148082.10 | SHC3 | -2.01828979 | 6.85E-15 |
| ENSG00000143512.13 | HHIPL2 | -2.003300443 | 0.000167 |
