## Supplementary Table.2 for "Evolutionary and functional dynamics of chimeric pseudogenes (φgenes)"

**Supplementary table 1: List of all ϕgenes from NCBI and Ensembl 38**

Table 1: List of 9 ϕgenes from NCBI post filteration

| genes5p-genes3p | Longest Read | E value | Per. Identity | Matched Sequence with Pseudogene | Sequence Details | Biotype (Ensembl) |
|---|---|---|---|---|---|---|
| BANP-RUNDC2A | CGAGGGCAACCTCCAGATC CATCACGTGGGGCAGGATG GTCAGAGCCCGTGTTCTGG TACTACGTGAAGGAGGTC | 2.00E-31 | 100.00% | sorting nexin 29 pseudogene 1 (SNX29P1) | Sequence ID: NR_045011.1, Length: 983 | Unprocessed |
| BANP-RUNDC2B | GGAGCAAGTCCAGATCACG CAGGACAGCGAGAGCCCGT GTTCTGGTACTACGTGAAGG AGGTCCTCAACAAGCACGA GCTG | 9.00E-35 | 100.00% | sorting nexin 29 pseudogene 2 (SNX29P2) | Sequence ID: NR_002939.3, Length: 1585 | Linc RNA |
| DHX40-RNFT1 | TTCGTGTGGAAGCTCAACTT CGAGAACTAATCAGGAAGC TTAAACAGGAGACAGAGGC CTGAAGCAAAGACATCTGG GTCAGAGAAAA | 1.00E-38 | 100.00% | TBC1D3P1-DHX40P1 readthrough, transcribed pseudogene (TBC1D3P1-DHX40P1) | Sequence ID: NR_002924.3, Length: 1945 | Linc RNA |
| GKAP1-KIF27 | ACAGCCTATGGGTGAAATTT GGCTTTCATTCATGAATGAG GAATCCAGATCTTACATAAG ATGGAAGTCTCTCACACTAG ATACTGAACATTAAATAGAA AAT | 7.00E-47 | 100.00% | kinesin family member 27 pseudogene (LOC389765) - NCBI only | Sequence ID: NR_029410.1, Length: 1764 | Not found |
| RP4-565E6.1-HYDIN | GGCCACGCCTGGGGTGCCT CCTGGAGTCCAGGCTGCTG GCGATGGAATCCATCATGTT GCTGATGTCACTGATCACGC CCATACGTTGACCTGTGTTA CTGAAAGAGAAAAGTTGATT GTACCCATCAAAGCTAG | 4.00E-61 | 98.52% | HYDIN2, axonemal central pair apparatus protein (pseudogene) (HYDIN2), non-coding RNA | Sequence ID: NR_103556.2Length: 10510 | Unprocessed |
| RMND5A-ANAPC1 | CAGTGTTTCTCGGGTTGGAA ACGCCATTGATAAGGATTCA CTTTAAGAGATTTGGAAACT CTTCCCTTTGGAATT | 8.00E-30 | 98.67% | anaphase promoting complex subunit 1 pseudogene (LOC285074), non-coding RNA | Sequence ID: NR_026846.1Length: 2225 | Unprocessed |
| PMS2L11-POLR2J3 | ACGCTACCTGTGCGCCATAA GGAATTTCAAAGGTATATTA AGAAGACGTGCCTGCTTCC CCTTCGACTTCTGCCG | 4.00E-28 | 97.33% | PMS1 homolog 2, mismatch repair system component pseudogene 3 (PMS2P3), non-coding RNA | Sequence ID: NR_028059.1Length: 1518 | Processed |
| GATSL1-GTF2I | CTGTTCACCTACGGCCTGAT CAAACTTGCCTTCCTGTCCT CCAAGACCAGAGGATGATT ATTCTCCACCGTCTAAGAGA CCAAAGGCCAATGAGCTAC CGCAGCCACCAGTCCCGGG | 2.00E-52 | 99.14% | general transcription factor IIi pseudogene 1 (GTF2IP1) | Sequence ID: NR_002206.3, Length: 3627 | Unprocessed |
| FRG1B-FRG1 | AGCTGAAGCCACTGCCCTT GAGAACCCTCTCGAGGAGT CTGGGCTCATGAGGATGCC AGAATAAGTGGCAGTAAGAA GAAAAAGAACAAAGATAAGA AAAGAAAAAGAGAAGAAGAT GAAGAAACCCAGCG | 4.00E-51 | 95.38% | FSHD region gene 1 family member J, pseudogene (FRG1JP) | Sequence ID: NR_033907.2, Length: 1487 | Not found |

**Supplementary table 1: List of all ϕgenes from NCBI and Ensembl 38**

Table 2: List of 9 ϕgenes from Ensembl 38 post filteration

| genes5p-genes3p | Longest Read | Query Cover | E value | Per. Identity | Matched Sequence with Pseudogene | Sequence Details | Biotype (Ensembl) |
|---|---|---|---|---|---|---|---|
| EMB-TRIM43 | CCAAGCTCGGCGGACGGCAGTGCCCCAGGAGTGAGTCCGTGCTGCTGCACATGCCCCAGC | 100 | 1.00E-37 | 100 | EMBP1 | ENST00000648011.1 | Processed transcript |
| FRG1-RP11-88I18.2 | GGCGTTCAGATGCAATTGGACCAAGAGAACAATGGGAACCAGTCTTTCAAAATGGTGCCTGTGCAGCAGTATTCACTGTGATAGGAAGTGAGAAGCAGTCTGAGT | 100 | 7.00E-53 | 100.00 | FRG1HP | ENST00000611606.4 | Processed transcript |
| GOLGA8G-AC103965.1 | GGGAGGCCATGGTGGCATTTTTCAACTCCGCTGGAGCCAATGCCCAGGAGGAACAAAGGGTGTGCTGCCAGCCCC | 100 | 3.00E-35 | 100 | GOLGA2P10 | ENST00000634182.1 | Processed transcript |
| HIC2-PI4KA | GCTCAGGAGGCCCCCCCCCCGCTGGCTGCTGCTCACATGGTGTCTGGGCCCTTGGCACTCCGTTCATGCGGGAGA | 100 | 3.00E-29 | 96.1 | PI4KAP1 | ENST00000611748.1, ENST00000612500.4 | Processed transcript |
| LRRC37A3-LRRC37A4 | CTGTCTGCAAGACAGTCAAGCTGCATTGCAACAGTGCATGTCTGACAAACACCATACATTGTCGTTTAGGATTGGGTGTGTGTGTCTCCTCTC | 100 | 8.00E-46 | 100.00 | LRRC37A17P | ENST00000570478.5 | Processed transcript |
| RP11-313P13.3-POLR2J3 | AGTCAGCGTGAAGCAGTTATTTTCTACGCTACCTGTGCGCCATAAGGAATTTCAAAGGAATATTAAGAAGACGTGCCTGCTTCCCCTTCGACTTCTGCCG | 100 | 1.00E-47 | 99 | PMS2P3 | ENST00000418756.5 | Processed transcript |
| RP11-488L18.8-ZNF195 | CATAGCTAAGCAGCCTGGACACCCGGGAAGCTGGGAAATGGTGTGATTTGTCCAGATAATC | 100 | 1.00E-37 | 100 | AL390728.4 | ENST00000606454.1 | Processed transcript |
| RP4-565E6.1-HYDIN | GGCCACGCCTGGGGTGCCTCCTGGAGTCCAGGCTGCTGGCGATGGAATCCATCATGTTGCTGATGTCACTGATCACGCCCATACGTTGACCTGTGTTACTGAAAGAGAAAAGTTGATTGTACCCATCAAAGCTAG | 100 | 7.00E-66 | 98.52 | HYDIN2 | ENST00000648235.1 | Processed transcript |
| GATSL1-GTF2I | CTGTTCACCTACGGCCTGATCAAACTTGCCTTCCTGTCCTCCAAGACCAGAGGATGATTATTCTCCACCGTCTAAGAGACCAAAGGCCAATGAGCTACCGCAGCCACCAGTCCCGGG | 99.145299 | 4.00E-29 | 100 | GTF2IP4, GTF2IP5 | ENST00000544802.5, ENST00000616377.5 | Processed transcript |

**Supplementary table 1: List of all ϕgenes from NCBI and Ensembl 38**

Table 3: List of final 9 ϕgenes from NCBI and Ensembl 38

| Name | Database | φgene | Length (bp) | 5' parent gene | Length (bp) | 3' parent gene | Length (bp) |
|---|---|---|---|---|---|---|---|
| *RP4-565E6.1-HYDIN* | *Both* | *HYDIN2* | 10510 | RP4-565E6.1 | 1523 | HYDIN | 8987 |
| *GATSL1-GTF2I* | *Both* | *GTF2IP1* | 3618 | GATSL1 | 404 | GTF2I | 3214 |
| *RMND5A-ANAPC1* | *Both* | *ANAPC1P2* | 2225 | RMND5A | 636 | ANAPC1 | 1332 |
| *GKAP1-KIF27* | *NCBI* | *ENSG00000290924* | 1955 | GKAP | 372 | KIF27 | 1583 |
| *HIC2-PI4KA* | *Ensembl* | *PI4KAP2* | 2272 | HIC2 | 258 | PI4KA | 2014 |
| *BANP-RUNDC2A* | *NCBI* | *SNX29P1* | 983 | BANP | 105 | SNX29 | 878 |
| *EMB-TRIM43* | *Ensembl* | *EMBP1* | 2614 | EMB1 | 1-85; 232-1215 | TRIM43 | 85-232 |
| *FRG1-RP11-88I18.2* | *NCBI* | *FRG1HP* | 945 | FRG1 | 1-175; 463-770 | RP11-88I18.2 | 771-945 |
| *PMS2L11-POLR2J3* | *NCBI* | *DTX2P1-UPK3B1-PMS2P11/PMS2P5* | 1782 | PMS2L11 | 1097-1322 | POLR2J3 | 1323-1763 |
