## Supplementary Table.3 for "Evolutionary and functional dynamics of chimeric pseudogenes (φgenes)"

**Supplementary table 2: Exon/ intron conservation of all 5 φgenes in 23 vertebrate species**

| Name of the species | RMND5A (143 bp) | ANAPC1 (179 bp) | BP (474 bp) | RMND5A intron 2 (7301) | ANAPC1 intron 24 (2365) | Intron fusion (NCBI) (2816 bp) |
|---|---|---|---|---|---|---|
| Drosophila M. | Nil | Nil | Nil | No | No | No |
| C. Elegans | Nil | Nil | Nil | No | No | No |
| Zebrafish | Nil | Nil | Nil | No | No | No |
| Tetraodon | Nil | Nil | Nil | No | No | No |
| Stickleback | Nil | Nil | Nil | No | No | No |
| X. tropicalis | Nil | Nil | Nil | No | No | No |
| Lizard | Nil | Nil | Nil | No | No | No |
| Zebra finch | Nil | Nil | Nil | No | No | No |
| Chicken | Nil | 3-161 | Nil | No | No | No |
| Platypus | 1-143 | Nil | Nil | 2% 1-153 (Chr 18) | No | No |
| Opossum | 1-143 | 1-179 | Nil | | | No |
| Elephant | 1-143 | 1-179 | Nil | 14% 1-1090 | No | No |
| Cow | 1-143 | 1-179 | Nil | 3-947, 4914-5193 (chr 11) | No | No |
| Dog | 1-143 | 1-179 | Nil | 1-1323, 6525-6796 (chr 17) | No | No |
| Cat | 1-143 | 1-179 | Nil | 8% 1-634 | No | No |
| Horse | 1-143 | 1-179 | Nil | 1-1332, 6031-6966 (chr 15) | No | No |
| Mouse | 1-143 | 1-179 | Nil | 2% 3-176 (chr 6) | No | No |
| Rat | 1-143 | Nil | Nil | 2% 3-156 (Chr 4) | No | No |
| Guinea Pig | 1-143 | 1-179 | Nil | 7% 1-546 | No | No |
| Rabbit | 1-143 | 1-179 | Nil | 4% 3-299 (chr 2) | No | No |

| | | | | | | |
|---|---|---|---|---|---|---|
| Marmoset | 1-143 | 1-179 | Nil | 75% (5464-7301, 3342-5049, 2013-3356, 1-438, 5243-5462 (chr 14) | 69% 1-1645 (chr 14) | No |
| Rhesus_macaque | 1-143 | 1-179 | Nil | 1 to 1383, 1442 to 7301 (Chr 13) | 4 to 2362(Chr 13) | No |
| Gibbon | 1-143 | 1-179 | Nil | 95.5% 1 to 7301 (few bases are missing) | 82% and 30% to different locations | No |
| Orangutan | 1-143 | 1-179 | Nil | 1 to 7301 chr 12 | 1 to 2365 (Chr 12) | No |
| Gorilla | 1-143 | 1-179 | Nil | 1 to 7301 (chr 2A) 1 duplication on chr 2A | 1 to 2365 (Chr 2A- 1 duplication on same chr) | No |
| Chimp | 1-143 | 1-179 | Nil | 1 to 7301 (chr 12) | 1 to 2365 (Chr 12- 1 duplication on same chr) | No |
| Bonobo | 1-143 | 1-179 | Nil | 1 to 7301 (chr 12) | 1 to 2365 (Chr 12- 1 duplication on same chr) | No |
| Human | 1-143 | 1-179 | 1-474 | 1 to 7301 (Chr 2) | 1 to 2365 (chr 2) | Yes |

| Name of the species | GATSL1 exon 1 (394Bp) | GTF2I exon 17 (111 Bp) | GATSL1-GTF2I BP (224 Bp) Refseq | GATSL1 intron 1 (42,789 Bp) | GTF2I Intron 717bp | GATSL1-GTF2I Intronic 616 BP |
|---|---|---|---|---|---|---|
| Drosophila M. | Nil | Nil | Nil | Nil | Nil | Nil |
| C. Elegans | Nil | Nil | Nil | Nil | Nil | Nil |
| Zebrafish | Nil | Nil | Nil | Nil | Nil | Nil |
| Tetraodon | Nil | Nil | Nil | Nil | Nil | Nil |
| Stickleback | Nil | Nil | Nil | Nil | Nil | Nil |
| X. tropicalis | Nil | Nil | Nil | Nil | Nil | Nil |
| Lizard | Nil | Nil | Nil | Nil | Nil | Nil |
| Zebra finch | Nil | Nil | Nil | Nil | Nil | Nil |
| Chicken | Nil | Nil | Nil | Nil | Nil | Nil |
| Platypus | Nil | Nil | Nil | Nil | Nil | Nil |
| Opossum | Nil | Nil | Nil | Nil | Nil | Nil |
| Elephant | Nil | Nil | Nil | Nil | Nil | Nil |
| Cow | 1-394 | Nil | Nil | Nil | Nil | Nil |
| Dog | 83- 394 | Nil | Nil | Nil | Nil | Nil |
| Cat | 1-394 | Nil | Nil | Nil | Nil | Nil |
| Horse | 58- 394 | Nil | Nil | Nil | Nil | Nil |
| Mouse | 234- 394 | Nil | Nil | Nil | Nil | Nil |
| Rat | 244- 394 | Nil | Nil | Nil | Nil | Nil |
| Guinea Pig | 205- 394 | Nil | Nil | Nil | Nil | Nil |
| Rabbit | 1-394 | 1-111 | Nil | Nil | Nil | Nil |
| Marmoset | 1-394 | 1-111 | Nil | 1- 42,798 | 1-717 | Nil |
| Rhesus_macaque | 1-394 | 1-111 | Nil | 1-42,789 | 87-717 | Nil |
| Gibbon | 1-394 | 1-111 | Nil | 1- 42,798 | 2-717 | Nil |
| Orangutan | 1- 394 | 1-111 | Nil | 1- 42,798 | 1-717 | Nil |
| Gorilla | 1-394 | 1-111 | Nil | 1- 42,798 | 1-717 | Nil |
| Chimp | 1-394 | 1-111 | Nil | 1- 42,798 | 1-717 | Nil |
| Bonobo | 1-394 | 1-111 | Nil | 1- 42,798 | 1-717 | Nil |
| Human | 1-394 | 1-111 | 1-224 | 1-42, 798 | 1-717 | 1-616 |

| Name of the species | HIC2 exon 1 (101BP) | PI4KA exon 33 (159 Bp) | HIC2-PI4KA BP (258 Bp) | HIC2 Intron (3075bp) | PI4KA Intron (1152BP ) | HIC2-PI4KA Intron BP (3612bp) |
|---|---|---|---|---|---|---|
| Drosophila M. | Nil | Nil | Nil | nil | Nil | nil |
| C. Elegans | Nil | Nil | Nil | Nil | Nil | Nil |
| Zebrafish | Nil | 1.-41 | Nil | nil | Nil | nil |
| Tetraodon | Nil | 4.-43 | Nil | Nil | Nil | Nil |
| Stickleback | Nil | 1.-43 | Nil | nil | Nil | nil |
| X. tropicalis | Nil | 1-155 | Nil | nil | Nil | nil |

| | | | | | | |
|---|---|---|---|---|---|---|
| Lizard | Nil | | Nil | | Nil | nil |
| Zebra finch | Nil | 1.-83 | Nil | nil | | |
| Chicken | Nil | 1.- 83 | Nil | nil | Nil | nil |
| Platypus | Nil | 1-159 | Nil | Nil | Nil | Nil |
| Opossum | Nil | 1-159 | Nil | nil | Nil | nil |
| Elephant | Nil | 1-159 | Nil | Nil | Nil | Nil |
| Cow | Nil | 1-159 | Nil | nil | Nil | nil |
| Dog | Nil | 4-155 | Nil | nil | Nil | nil |
| Cat | Nil | 4-155 | Nil | nil | Nil | nil |
| Horse | Nil | 4- 155 | Nil | Nil | Nil | Nil |
| Mouse | Nil | 4-158 | Nil | Nil | Nil | Nil |
| Rat | Nil | 1-155 | Nil | nil | Nil | nil |
| Guinea Pig | Nil | 4-155 | Nil | nil | Nil | nil |
| Rabbit | Nil | 1-158 | Nil | nil | Nil | nil |
| Marmoset | Nil | 1-159 | Nil | 21-3075 | 1-1015 | nil |
| Rhesus_macaque | Nil | 1-159 | Nil | 1.-3075 | 121-1152 | Nil |
| Gibbon | Nil | 1-159 | Nil | 1.-3075 | 1-1152 | nil |
| Orangutan | Nil | 1-159 | Nil | 1.-3075 | 1- 1152 | nil |
| Gorilla | Nil | 1-159 | Nil | 1.-3075 | 1-1152 | Nil |
| Chimp | Nil | 1-159 | Nil | 1.-3075 | 1- 1152 | Nil |
| Bonobo | Nil | 4-159 | Nil | 1.-3075 | 1- 1152 | Nil |
| Human | 1-101 | 1-159 | 1-258 | 1.-3075 | 1- 1152 | 1 to 2012 |

| **Name of the species** | **GKAP Exon-6 (132Bp)** | **KIF27 Exon 2 (385bp)** | **GKAP-KIF25 bp(401bp)** | **GKAP1 Intron 6 (10,435bp)** | **KIF27 Intron 1 (6641bp)** | **GKAP1-KIF27 intron BP** |
|---|---|---|---|---|---|---|
| Drosophila M. | Nil | Nil | Nil | Nil | Nil | Nil |
| C. Elegans | Nil | Nil | Nil | Nil | Nil | Nil |
| Zebrafish | Nil | Nil | Nil | Nil | Nil | Nil |
| Tetraodon | Nil | Nil | Nil | Nil | Nil | Nil |
| Stickleback | Nil | Nil | Nil | Nil | Nil | Nil |
| X. tropicalis | Nil | Nil | Nil | Nil | Nil | Nil |
| Lizard | Nil | Nil | Nil | Nil | Nil | Nil |
| Zebra finch | Nil | Nil | Nil | Nil | Nil | Nil |
| Chicken | Nil | Nil | Nil | Nil | Nil | Nil |
| Platypus | Nil | Nil | Nil | Nil | Nil | Nil |
| Opossum | Nil | 86 - 385 | Nil | Nil | Nil | Nil |
| Elephant | Nil | 1 - 380 | Nil | Nil | Nil | Nil |

| | | | | | | |
|---|---|---|---|---|---|---|
| Cow | 1-132 | 4 - 382 | Nil | Nil | Nil | Nil |
| Dog | Nil | 1 - 385 | Nil | Nil | Nil | Nil |
| Cat | Nil | 65 - 358 | Nil | Nil | Nil | Nil |
| Horse | Nil | 4 - 385 | Nil | Nil | Nil | Nil |
| Mouse | Nil | 4 - 385 | Nil | Nil | Nil | Nil |
| Rat | Nil | 1 - 381 | Nil | Nil | Nil | Nil |
| Guinea Pig | Nil | 3 - 385 | Nil | Nil | Nil | Nil |
| Rabbit | Nil | 4 - 385 | Nil | Nil | Nil | Nil |
| Marmoset | 1-132 | 1-385 | Nil | 1-10,435 | 4193 - 5692bp<br>704 - 1123bp<br>4705 - 4929bp | Nil |
| Rhesus_macaque | 1 - 132 | 1-385 | Nil | 1-10,435 | 1559 - 2955bp<br>2941 - 4182bp<br>534 - 1540bp<br>4890 - 5692bp<br>1 - 471bp | Nil |
| Gibbon | 1 - 132 | 1-385 | Nil | 1 - 4636bp<br>6895 - 10435bp | 1559 - 5692bp<br>502 - 1541<br>1 - 503bp | Nil |
| Orangutan | 1-132 | 1-385 | Nil | 4754 - 10435bp<br>1 - 4712bp | 1559 - 5692bp<br>470 - 1541<br>1 - 535bp<br>5281 - 5692bp | 1 – 958bp chr 13 one range |

| | | | | | | |
|---|---|---|---|---|---|---|
| Gorilla | 1-132 | 1-385 | Nil | 4767 - 104435bp<br>1 - 4734bp | 1559 - 5692bp<br>1 - 1541bp<br>5281 - 5692bp<br>5281 - 5692bp | 1 – 958bp chr 9 Two ranges on same chromosome 56902889 – 56903630 and |
| Chimp | 1-132 | 1-385 | Nil | 1-10,435 | 470 - 5692bp<br>1 - 535bp<br>5281 - 5692bp | 1 – 958bp chr 11 one range |
| Bonobo | 1-132 | 1-385 | Nil | 1-10,435 | | 1- 958bp chr 11 one |
| Human | 1-132 | 1-385 | 1-401 | 1-10,435 | 1 - 5692bp<br>5281 - 5692bp | 1 – 958bp chr 9 one |
| **Name of the species** | **RP4-565E6.1 Exon-6 (323Bp)** | **HYDIN Exon 7 (200Bp)** | **RP4-565E6.1-HYDIN BP (243 Bp)** | **RP4-565E6.1 Intron 15 (21,565bp)** | **RP4-565E6.1-HYDIN Intron BP (420Bp)** | |
| Drosophila M. | NIL | NIL | nil | Nil | NIL | |
| C. Elegans | NIL | NIL | nil | Nil | NIL | |
| Zebrafish | NIL | NIL | nil | Nil | NIL | |
| Tetraodon | NIL | NIL | nil | Nil | NIL | |
| Stickleback | NIL | NIL | nil | Nil | NIL | |
| X. tropicalis | NIL | NIL | nil | Nil | NIL | |
| Lizard | NIL | NIL | nil | Nil | NIL | |
| Zebra finch | NIL | NIL | nil | Nil | NIL | |
| Chicken | NIL | NIL | nil | Nil | NIL | |
| Platypus | NIL | 17-167 | nil | Nil | NIL | |
| Opossum | NIL | 17-169 | nil | Nil | NIL | |

| | | | | | |
|---|---|---|---|---|---|
| Elephant | NIL | NIL | nil | Nil | NIL |
| Cow | NIL | 1-195 | nil | Nil | NIL |
| Dog | NIL | 15-194 | nil | Nil | NIL |
| Cat | NIL | 1-195 | nil | Nil | NIL |
| Horse | NIL | 1-195 | nil | Nil | NIL |
| Mouse | NIL | 1-171 | nil | nil | NIL |
| Rat | NIL | 17-171 | nil | nil | NIL |
| Guinea Pig | NIL | 1-195 | nil | 1232 – 2037bp | NIL |
| Rabbit | NIL | 1-195 | nil | 315 – 1,913bp | NIL |
| Marmoset | NIL | 1-200 | nil | 1 – 6,141bp and 18,254 – 21,558bp | NIL |
| Rhesus_macaque | 1-322 | 1-200 | nil | 1 – 21,565bp | NIL |
| Gibbon | 1-323 | 1-200 | nil | 1 – 21,565bp | NIL |
| Orangutan | 1-323 | 1-200 | nil | 1 – 21,565bp | NIL |
| Gorilla | 1-323 | 1-200 | nil | 1 – 21,565bp | NIL |
| Chimp | 1-323 | 1-200 | nil | 1 – 21,565bp | NIL |
| Bonobo | NIL | 1-200 | nil | 1 – 21,565bp | NIL |
| Human | 1-323 | 1-200 | 1-243 | 1 – 21,565bp | 1-420 |
