## Supplementary Table.4 for "Evolutionary and functional dynamics of chimeric pseudogenes (φgenes)"

**Supplementary Table 3.Splicing site predictions around intron fusion and their respective parental sequences for 5 φgenes.**

Splice sites retained from 5' parental sequecne
Splice sites retained from 3' parental sequecne

| RMND5A-ANAPC1 (Donor sites) | | | |
|---|---|---|---|
| Start | End | Score | Exon-Intron |
| 137 | 151 | 0.95 | tgataag**gt**atggta |
| 625 | 639 | 0.76 | tgaatga**gt**gagtgg |
| 1008 | 1022 | 0.89 | aacagag**gt**aataga |
| 1149 | 1163 | 0.86 | ttcactg**gt**aggtgt |
| 1171 | 1185 | 0.9 | caagact**gt**gagtgg |
| 1467 | 1481 | 0.87 | ctgtttg**gt**aactcc |
| 1715 | 1729 | 0.97 | aggtacg**gt**aactac |
| 1871 | 1885 | 0.87 | acagaag**gt**cagcat |
| 2082 | 2096 | 0.93 | atgccat**gt**atgtcc |
| 2161 | 2175 | 0.93 | tcacaag**gt**caggag |

| RMND5A-ANAPC1 (Acceptor sites) | | | |
|---|---|---|---|
| Start | End | Score | Intron-Exon |
| 181 | 221 | 0.88 | tcattctttttcttctgtc**ag**tgctttgataatactagctt |
| 255 | 295 | 0.93 | tgcttttgatgacttttac**ag**gtctagttttactatgttat |
| 344 | 384 | 0.92 | taaaaccctctgcttttat**ag**ctttgttgtttatccatttg |
| 1617 | 1657 | 0.96 | cactttattttcttccagc**ag**ctgtatccttctagaatgtg |
| 1694 | 1734 | 0.89 | aagaaccctcagttcctct**ag**aggtacggtaactaccggtg |
| 2043 | 2083 | 0.9 | caccaattatgctttctgc**ag**ctggtagggtgaccatatat |
| 2087 | 2127 | 0.82 | atgtatgtccctgtttcct**ag**tttatgtttttttgtcctgtc |
| 2419 | 2459 | 0.78 | aattattattgctctacat**ag**ttgccaacatttagtattga |
| 2550 | 2590 | 0.96 | catttatgtttccccttct**ag**gaaatgtctttttgtccttt |
| 2585 | 2625 | 0.93 | tcctttatatttttctatt**ag**attgtcttttttttcttcagg |
| 2604 | 2644 | 0.99 | agattgtcttttttttcttc**ag**gtttttacaagttctttcaa |
| 2788 | 2828 | 0.77 | ctctcttcctttaaccatt**ag**gattcactttaagagatttg |

| GKAP1-KIF27 (Donor sites) | | | |
|---|---|---|---|
| Start | End | Score | Exon-Intron |
| 253 | 267 | 0.71 | cattaag**gt**gcttca |
| 858 | 872 | 0.98 | acttaca**gt**aagtgt |
| 1630 | 1644 | 0.99 | gtagagg**gt**aaggtc |
| 1809 | 1823 | 0.98 | caacgtg**gt**gagacc |
| 1893 | 1907 | 0.98 | agtttac**gt**aagcaa |
| 2210 | 2224 | 0.92 | tacaggc**gt**aagcca |
| 2623 | 2637 | 0.99 | aataata**gt**aagttt |
| 2679 | 2693 | 0.76 | ggaacag**gt**cagaga |
| 2938 | 2952 | 0.72 | tttttgggt**at**gaag |
| 3616 | 3630 | 0.96 | attacag**gt**gtgcac |

| | | | |
|---|---|---|---|
| 3797 | 3811 | 0.89 | gactaat**gt**atgtaa |
| 4516 | 4530 | 0.82 | ggctgag**gt**aggagg |
| 4626 | 4640 | 0.86 | gtcccag**gt**actccg |
| 4882 | 4896 | 0.87 | aagcata**gt**gagtaa |
| 4916 | 4930 | 0.96 | cattttg**gt**atgtgt |
| 5340 | 5354 | 0.89 | tgtgcag**gt**tagtta |
| 5408 | 5422 | 0.78 | gcattag**gt**atatct |
| 7311 | 7325 | 0.96 | aataata**gt**aagaac |
| 7552 | 7566 | 0.92 | ggcttgt**gt**aagtac |
| 7746 | 7760 | 0.94 | acttaca**gt**aagtag |
| 7786 | 7800 | 0.87 | gctccag**gt**cagtgg |
| 8972 | 8986 | 0.81 | tgatgag**gt**agatta |
| 9169 | 9183 | 0.98 | taccctg**gt**aagata |
| 9502 | 9516 | 0.74 | aaaaaaa**gt**gtgtac |
| 10256 | 10270 | 0.95 | catattg**gt**aagatt |
| 10332 | 10346 | 0.91 | acacatg**gt**aggtgt |
| 13300 | 13314 | 0.84 | ttcattt**gt**aagtca |
| 14038 | 14052 | 0.71 | cacgcct**gt**aagctc |
| 15441 | 15455 | 0.83 | agattat**gt**atgtat |
| 15717 | 15731 | 0.85 | attacag**gt**gtgagc |
| 16330 | 16344 | 0.95 | cctccag**gt**acaatt |
| 16543 | 16557 | 0.99 | attcttg**gt**aagggg |

| GKAP1-KIF27 (Acceptor sites) | | | |
|---|---|---|---|
| **Start** | **End** | **Score** | **Intron-Exon** |
| 338 | 378 | 0.87 | ttcactctttaccttcgcc**ag**atttttcagagtagatgcta |
| 585 | 625 | 0.98 | ctagatttcttttctcaac**ag**atttataacactatagtagc |
| 749 | 789 | 0.95 | tttaactttcttggcctac**ag**aaagtgggagctgcagattg |
| 947 | 987 | 0.77 | agttttgctcttgttgccc**ag**gctggagtgtaatggcacga |
| 997 | 1037 | 0.98 | acagcaccttctgccttcc**ag**gttcaagcaactctcctgcc |
| 1085 | 1125 | 0.98 | tttttttttttgtatttttagtac**ag**atggtgtttcgcctt |
| 1295 | 1335 | 0.97 | cacaattttttatttcacc**ag**gtgttaaatcagtgatactt |
| 1413 | 1453 | 0.81 | ttaataccttttttttggtc**ag**gctaattggtagtgttaata |
| 1597 | 1637 | 0.82 | tgctgtctgtcttgattct**ag**tgaggtcatagtgtagaggg |
| 1963 | 2003 | 0.87 | agtctctcgctctgttgcc**ag**gctggaatgtggtggcgcaa |
| 2103 | 2143 | 0.96 | tagtttttgtatttttaat**ag**agatggggtttcactacgtt |
| 2405 | 2445 | 0.95 | tatttctttttctacttgtag**ag**cacatttttcactgcata |
| 2426 | 2466 | 0.83 | agcacatttttcactgcat**ag**acagctgtgtacttgaagaa |
| 2498 | 2538 | 0.8 | aatgtgtcttctatgtgcc**ag**gtcctgtggtatttgtaagg |
| 3647 | 3687 | 0.8 | ccttgaaattatttttcaa**ag**gaactcaaagggaagagggg |
| 3972 | 4012 | 0.83 | atttatcctaacatctctt**ag**atgtgactgtatgtggagat |
| 4434 | 4474 | 0.96 | ttgagattttttttttttta**ag**aactaatatcattgtagagc |
| 5111 | 5151 | 0.79 | ctattttttttttaagtat**ag**catagctcagatgattgaac |
| 5163 | 5203 | 0.91 | tttcatctcctatcccctt**ag**ctgcagattaagtagtgcta |
| 5547 | 5587 | 0.83 | ggttttttgttcttgcgat**ag**tttactgagaatgatggttt |
| 6160 | 6200 | 0.72 | acttgtttgagttcattgt**ag**attctggatattagcccttt |
| 6221 | 6261 | 0.98 | aaattttctcccatgttgt**ag**gttgcctgttcactctgatg |

| | | | |
|---|---|---|---|
| 6777 | 6817 | 0.97 | ctttccattttaccctttt**ag**aagttaagacatgttaacac |
| 7030 | 7070 | 0.78 | tgactcttgggtttctggt**ag**ataggtattttccttctgta |
| 7051 | 7091 | 0.84 | ataggtattttccttctgt**ag**tatgttatgtgtaatcttta |
| 7072 | 7112 | 0.95 | tatgttatgtgtaatcttt**ag**gtctttgactcttttcagtg |
| 7090 | 7130 | 0.79 | taggtctttgactcttttc**ag**tggctgaccactcacctact |
| 7142 | 7182 | 0.85 | aaaaactttttatttcaa**ag**attaaagccttagcttatat |
| 7407 | 7447 | 0.99 | tttattgtaccttttcttt**ag**atatacaaatacttaccatt |
| 7692 | 7732 | 0.82 | taacttattcagtctttac**ag**caactcaagagagatcaagt |
| 8009 | 8049 | 0.81 | aaaacttcagactcttttc**ag**ggtttctaatgagagaattg |
| 8996 | 9036 | 0.95 | ggagcacttcccttttcat**ag**ttgacataactccagagcag |
| 9102 | 9142 | 0.93 | cagaggcttttactttttc**ag**gggtttcctggctgccccaa |
| 9182 | 9222 | 0.71 | tagctctgccttatgctac**ag**aaagaaaatttctgtgagcc |
| 9412 | 9452 | 0.8 | aagctaccatgtccaccac**ag**gttggtaaaggggctttgtt |
| 9857 | 9897 | 0.77 | ctttttatttttgatacac**ag**gaatccagatcttacataag |
| 10659 | 10699 | 0.91 | tctttttttttcatattct**ag**ccagatttcttttttaaatt |
| 11198 | 11238 | 0.82 | aacaatctgtatattctgt**ag**gattaggagatcacagatac |
| 11287 | 11327 | 0.94 | taaattgtggcttttcttt**ag**gattgtagggtttttttttt |
| 12126 | 12166 | 0.79 | tgagcattttctgtttccc**ag**aatgtctctatagaacacca |
| 12207 | 12247 | 0.97 | gttttctttttccccctcc**ag**tgaatatttattgagcatct |
| 12532 | 12572 | 0.99 | tttttttttaattttttgc**ag**agacaaggttttgccatgtt |
| 12701 | 12741 | 0.98 | attcattttcctgcttttc**ag**aacgttggaattacctttac |
| 12962 | 13002 | 0.98 | attgtgctttgttatctat**ag**atgtggggagtaactgaact |
| 13063 | 13103 | 0.74 | aggtttagatatttcctac**ag**gctagttcctataagctctt |
| 13666 | 13706 | 0.98 | tttactttctgtctctttt**ag**tttgcctattctagatattt |
| 13680 | 13720 | 0.9 | cttttagtttgcctattct**ag**atatttcatataagtggagg |
| 14503 | 14543 | 0.83 | tatttttccttctgtgtct**ag**cttctctctcttagcatgtt |
| 14517 | 14557 | 0.8 | tgtctagcttctctctctt**ag**catgttatgtcacctatcag |
| 14639 | 14679 | 0.74 | tggtttgcttccccccttgt**ag**ctcctgtaaataatgttgct |
| 15228 | 15268 | 0.94 | tcagtttcctgctttttac**ag**ctattttgtgctttagaaat |
| 15244 | 15284 | 0.76 | tacagctattttgtgcttt**ag**aaattatatgtcatctaaaa |
| 15543 | 15583 | 0.83 | caagtaatcctcccttctc**ag**cctcccaagcagctgagacc |
| 15610 | 15650 | 0.99 | aattttttgtgttttttgt**ag**acaaggggtcttgccatgtt |
| 15902 | 15942 | 1 | tctttttttttttttttt**ag**acagtttccctctgtcgctc |
| 15924 | 15964 | 0.78 | cagtttccctctgtcgctc**ag**gctggagtgcagtggcgcta |
| 16068 | 16108 | 0.95 | taatttttgtatttttagt**ag**ggacagggtttcaccatgtt |
| 16202 | 16242 | 0.98 | ctctttcaagtcttttctc**ag**atgtcaccttctgaatgatg |
| 16316 | 16356 | 0.99 | gccgtgttttttttcctcc**ag**gtacaattaacactttttaa |
| 16397 | 16437 | 0.91 | ttgtctgtctttcctcagc**ag**aatctcaccagggaagggat |
| 16825 | 16865 | 0.92 | taacttttcttttgtttta**ag**ttccagatatgctttctttt |
| 16858 | 16898 | 0.85 | tttctttttctaaattttcagcttc**ag**ttgtggagggccaa |

| RP4-HYDIN (Donor sites) | | | |
|---|---|---|---|
| **Start** | **End** | **Score** | **Exon-Intron** |
| 113 | 127 | 0.79 | ctgtatt**gt**aagcag |
| 360 | 374 | 0.83 | tttacag**gt**tcgggc |
| 683 | 697 | 0.97 | tgtcact**gt**aagtcg |
| 1119 | 1133 | 0.79 | aattaat**gt**aggttc |
| 1580 | 1594 | 0.92 | agataag**gt**aatgtt |

| | | | |
|---|---|---|---|
| 2212 | 2226 | 0.85 | gaatctg**gt**atgcag |
| 2285 | 2299 | 0.75 | acaggag**gt**gattag |
| 3587 | 3601 | 0.91 | gcaacag**gt**gtggac |
| 3943 | 3957 | 0.99 | ccattag**gt**tagtgt |
| 4190 | 4204 | 0.98 | tcaccag**gt**taggag |
| 4954 | 4968 | 0.95 | agttcag**gt**aaaaga |
| 5487 | 5501 | 0.77 | aaagcag**gt**ttgtga |
| 5581 | 5595 | 0.86 | attaatt**gt**aagcca |
| 5866 | 5880 | 0.89 | cggatat**gt**gagtac |
| 6428 | 6442 | 0.92 | actacag**gt**gtgagc |
| 6663 | 6677 | 0.9 | ttttgat**gt**aagcat |
| 7921 | 7935 | 1 | acagaag**gt**aaggga |
| 8021 | 8035 | 0.94 | tgaagag**gt**gagggg |
| 8197 | 8211 | 0.82 | agcctag**gt**actttt |
| 8397 | 8411 | 0.71 | tttgaag**gt**agatcc |
| 9613 | 9627 | 0.95 | tcatcag**gt**agggaa |
| 9705 | 9719 | 0.73 | catccag**gt**gcacat |
| 10010 | 10024 | 0.78 | tcacgag**gt**caggat |
| 10781 | 10795 | 0.79 | tctgctg**gt**acgaaa |
| 10831 | 10845 | 0.91 | cttgtag**gt**atgcat |

| RP4-HYDIN (Acceptor sites) | | | |
|---|---|---|---|
| **Start** | **End** | **Score** | **Intron-Exon** |
| 346 | 386 | 0.92 | tgagattgtttggatttac**ag**gttcgggccacaacatccgt |
| 1064 | 1104 | 0.79 | tctaattgtctcctgtctc**ag**agacctcatgattatttctg |
| 1250 | 1290 | 0.83 | cttggttctattcttgccc**ag**tcccattgggataacagcgg |
| 1389 | 1429 | 0.86 | tttcttcattttgctatgc**ag**ctattgcacagtgggggaaa |
| 1551 | 1591 | 0.97 | actttatatttttcttttt**ag**tgggacctagataaggtaat |
| 2430 | 2470 | 0.96 | tccacttcctcttccttct**ag**ctttctcttcttcactaatt |
| 2805 | 2845 | 0.76 | acttcatctgatctttacc**ag**ttttcctgcaaatatcattt |
| 2939 | 2979 | 0.96 | aatatcatttttctgcctc**ag**gagcctatccgggaccccgc |
| 2986 | 3026 | 0.94 | ttgagttgtatttctccct**ag**ttgactccaatctgtgacag |
| 3140 | 3180 | 0.83 | cctcagttcttgccccctc**ag**aagaaagaatttgactgagg |
| 3324 | 3364 | 0.83 | cttgatcctaagactttat**ag**gctggcctctttcccatgat |
| 3573 | 3613 | 0.79 | gttcactctttagtgcaac**ag**gtgtggaccatcaggaaaag |
| 3632 | 3672 | 0.77 | ctgccaatttatcactttt**ag**agaggcaatgtgataactgc |
| 3768 | 3808 | 0.91 | gtagaatgtccttcacttt**ag**gtttgtctaatgttttgtca |
| 3854 | 3894 | 0.8 | tgatactgtatccctttc**ag**cacatcatgccaagggattc |
| 4336 | 4376 | 0.85 | ccttttgagatgccttttc**ag**atttttgcgtttctgaaaat |
| 4698 | 4738 | 0.87 | ggttacaactctgccccac**ag**ggtgacatcgaggatgggaa |
| 5039 | 5079 | 0.97 | ttttttttttttttgaagc**ag**ggttttgctcttgttgc |
| 5063 | 5103 | 0.83 | ggttttgctcttgttgcct**ag**gctggagtgcaatggtgtga |
| 5204 | 5244 | 0.96 | aattttttgtattttt**ag**tagagatggggtttcactatgtt |
| 5342 | 5382 | 0.81 | atgtatacttttattgtac**ag**ctgccaggatttactgatgg |
| 5835 | 5875 | 0.97 | gctgccctttaattccttct**ag**aacaaggcagcggatatgtg |
| 6059 | 6099 | 0.91 | aaatagtacttatctcttt**ag**gattctgtagtaatcagact |
| 6158 | 6198 | 0.79 | tttgtgcttcctcatgtct**ag**atagcataagttttgagtct |

| | | | |
|---|---|---|---|
| 6390 | 6430 | 0.73 | tcaagcagttctctgcctc**ag**cttactgagtagctgagact |
| 6447 | 6487 | 0.77 | atgccccgctaatttttat**ag**aggctgggttttgccatgtt |
| 6829 | 6869 | 0.89 | tatgactcactcctgcctt**ag**gacttgtatgtttgcagttc |
| 6859 | 6899 | 0.84 | gtttgcagttccccttggc**ag**gagcactctgttcctagatc |
| 6908 | 6948 | 0.98 | ctggctccttttgctgttc**ag**gtccccatccgaaggtcacc |
| 7071 | 7111 | 0.76 | attatccttatcagtatat**ag**atggctagatggctccctac |
| 7304 | 7344 | 0.92 | gcctccctcttcccgctgc**ag**tccgtggcccacatagcagc |
| 7500 | 7540 | 0.89 | cttgctcccctctggccac**ag**gttccaaatcacacttcccc |
| 7589 | 7629 | 0.74 | tgaagtgtcacctcctcat**ag**aggcctttcctgaccgcccc |
| 7766 | 7806 | 0.86 | agatactaggctctgttct**ag**gtactgggtatacagaaggc |
| 8900 | 8940 | 0.71 | cctaattggttgtattcct**ag**acctagcacagtgctgtgct |
| 9293 | 9333 | 0.75 | tttttctttgctacactgc**ag**gcaaatattctatttctatc |
| 9599 | 9639 | 0.77 | ttgcctgggccctctcatc**ag**gtagggaactctgcttattg |
| 10451 | 10491 | 0.97 | ctctctctctctctctctc**ag**aaaagacaatctcatttgaa |
| 10562 | 10602 | 0.79 | atcgacccttttcttgaac**ag**ttatcgtgatcatgttgctt |
| 10617 | 10657 | 0.96 | gatcatacccttcccttgt**ag**gattacgcccatacgttgac |
| 10856 | 10896 | 0.76 | cttgtttttgattgattat**ag**ctaaatctagtcagctcaag |
| 10981 | 11021 | 0.93 | agacttcctcttcatcctc**ag**aggaaacaaaaattcaaggt |
| 11082 | 11122 | 0.91 | aaagtccttcttttagtac**ag**agaatggcccatgagaatat |

| **GATSL1-GTF2I (Donor sites)** | | | |
|---|---|---|---|
| **Start** | **End** | **Score** | **Exon-Intron** |
| 398 | 412 | 1 | agaccag**gt**aagcgc |
| 751 | 765 | 0.87 | aaaacag**gt**gtgggg |
| 1249 | 1263 | 1 | atgccag**gt**aagggt |
| 1490 | 1504 | 0.95 | ggtcaag**gt**tagcgt |
| 1953 | 1967 | 0.76 | cttggag**gt**aacaga |
| 2413 | 2427 | 0.76 | tcagaag**gt**gcggaa |
| 2715 | 2729 | 0.93 | catccag**gt**agaacc |
| 3081 | 3095 | 0.73 | acctcag**gt**gatcca |
| 4142 | 4156 | 0.98 | gtcagag**gt**gcgtgc |
| 4291 | 4305 | 1 | cccccag**gt**acgttc |
| 4370 | 4384 | 0.85 | gctcaag**gt**caggca |
| 4891 | 4905 | 0.9 | acgcccg**gt**aaattt |
| 5665 | 5679 | 0.94 | accctgg**gt**aggtct |
| 6542 | 6556 | 0.97 | acctcag**gt**aatcct |
| 6986 | 7000 | 0.91 | actcaag**gt**ggggcc |
| 7574 | 7588 | 0.85 | attacag**gt**gtgagc |
| 7710 | 7724 | 0.85 | attacag**gt**gtgagc |
| 8308 | 8322 | 0.75 | ttacgag**gt**atttat |
| 8564 | 8578 | 0.99 | ctgtaac**gt**aagtgg |
| 8640 | 8654 | 0.8 | agctcca**gt**aagctt |
| 9295 | 9309 | 0.92 | gtacggg**gt**gagccc |
| 9540 | 9554 | 0.99 | tgaaaga**gt**aagttg |
| 9590 | 9604 | 0.82 | caaattg**gt**atgctc |
| 9690 | 9704 | 0.99 | aaaagac**gt**gagtcc |
| 9717 | 9731 | 0.9 | agcaaag**gt**agaggg |
| 10107 | 10121 | 0.73 | acctcag**gt**gatcca |

| | | | |
|---|---|---|---|
| 10251 | 10265 | 0.74 | gttttag**gt**aaccat |
| 10421 | 10435 | 0.92 | gttacag**gt**accacc |
| 10812 | 10826 | 0.93 | tcacaag**gt**caggag |
| 11974 | 11988 | 0.93 | aagtctt**gt**gagtcc |
| 12054 | 12068 | 0.93 | gtgcctg**gt**gagtgg |
| 12267 | 12281 | 0.8 | caagcgg**gt**acatcg |
| 12281 | 12295 | 0.95 | gcctgag**gt**cagtag |
| 13267 | 13281 | 0.85 | attacag**gt**gtgagc |
| 14058 | 14072 | 0.93 | ctgttgg**gt**aagatg |
| 14157 | 14171 | 0.88 | taagatg**gt**aggttt |
| 14252 | 14266 | 1 | ggctgag**gt**gagtag |
| 14539 | 14553 | 0.91 | ttaagag**gt**acatgt |
| 14579 | 14593 | 0.9 | catactg**gt**agggac |
| 14983 | 14997 | 0.77 | catacgc**gt**gagccg |
| 15449 | 15463 | 0.98 | actacag**gt**gtgcac |
| 15591 | 15605 | 0.94 | tataggc**gt**aagcca |
| 16156 | 16170 | 1 | ccccctg**gt**aagagc |
| 16727 | 16741 | 0.73 | acctcag**gt**gatcca |
| 17361 | 17375 | 0.7 | tgacaaa**gt**gagacc |
| 17538 | 17552 | 0.97 | cttagag**gt**gaggct |
| 17562 | 17576 | 0.98 | ccctggg**gt**aagggg |
| 17963 | 17977 | 0.91 | taatatg**gt**aaaacc |
| 18460 | 18474 | 0.96 | attacag**gt**gtgcac |
| 19233 | 19247 | 0.91 | gccgcag**gt**gtgtgc |

| GATSL1-GTF2I (Acceptor sites) | | | |
|---|---|---|---|
| **Start** | End | Score | **Intron-Exon** |
| 379 | 419 | 0.73 | cttgccttcctgtcctcca**ag**accaggtaagcgcgcgggga |
| 646 | 686 | 0.9 | ccaccccctcccccatcgc**ag**ccccctccccgaaatccggg |
| 1235 | 1275 | 0.73 | cttggattttccctatgcc**ag**gtaagggtttgacattgtga |
| 1966 | 2006 | 0.92 | gaacagccttgtttttcct**ag**ccttggccctcagccctggg |
| 2289 | 2329 | 0.98 | ccccccttcctccctcctgc**ag**aaagcagcctgtcctgtccc |
| 2741 | 2781 | 0.82 | tcctgcaatacctgcccct**ag**gtggtgacccatagctctgt |
| 2862 | 2902 | 0.97 | ttttttttttttttttttgagactg**ag**ttttgctcttgtcc |
| 2958 | 2998 | 0.72 | caagtgattctcctgcctc**ag**cctcccgagtagctgggatt |
| 3253 | 3293 | 0.99 | ttttttattttctcttttagagac**ag**agtcttgctctgtc |
| 3279 | 3319 | 0.75 | gagtcttgctctgtcgtac**ag**gctggagtacagtggcgcca |
| 3604 | 3644 | 0.86 | tttccctccgctccttagc**ag**gcttaagtcacttctctctc |
| 3886 | 3926 | 0.89 | ttttttttgtttgtttttta**ag**agatggggtctctgttgccc |
| 4295 | 4335 | 0.88 | caggtacgttctgtgtatt**ag**gtattaaacctgctttgcaa |
| 4492 | 4532 | 0.79 | gtccagcccttctgtctac**ag**cacagtgttctgaaaggctt |
| 4601 | 4641 | 0.92 | tttttttttttttttttggagat**ag**ggtcttcttctgtcac |
| 4625 | 4665 | 0.76 | gggtcttcttctgtcaccc**ag**gccggagtgcagtggcgtga |
| 4715 | 4755 | 0.97 | agagctgttttcttctccc**ag**ggatacttggtccttatcac |
| 5138 | 5178 | 0.93 | ttgtctttttctcttacct**ag**attatcaaccttcaagtttt |
| 5251 | 5291 | 0.89 | tgactttgccatattttgc**ag**agcggtgaagacactagctt |
| 5369 | 5409 | 0.85 | gtgtagcaccctgctttgt**ag**gcacttagtaaacgtctgtt |
| 5822 | 5862 | 0.9 | gcctttgttctttctgcccag**ag**tattacttcttgccttac |

| | | | |
|---|---|---|---|
| 5869 | 5909 | 0.93 | cttccccgtgcatccttct**ag**attccatttcagcttccctg |
| 5881 | 5921 | 0.84 | tccttctagattccatttc**ag**cttccctgagggcacgttgg |
| 5921 | 5961 | 0.93 | ggctcactcttcttctggt**ag**gacgggcttaactcaaaacg |
| 6477 | 6517 | 0.87 | ctaattttgtatttttagtag**ag**acggggtttagccgtgtt |
| 7464 | 7504 | 0.96 | tttattttactctgtttttagt**ag**agacaaggtctcactgt |
| 7602 | 7642 | 0.92 | taatttttgtatttttagtag**ag**atggggtttcaacatgtt |
| 8073 | 8113 | 0.94 | actgcaaccttcccctccc**ag**gttcaagcaattctcctgcc |
| 8160 | 8200 | 0.89 | agctaatttttgtatttttagt**ag**agacggggtttcaccat |
| 8657 | 8697 | 0.96 | ttttttttttttttttttgagac**ag**agtcttgctctgttgc |
| 8731 | 8771 | 0.88 | actgcaacctctgtctccc**ag**gttcaagcgattctcctgct |
| 9056 | 9096 | 0.77 | aggacctcacctgtcattt**ag**gtgtcaggcctctggcaggg |
| 9425 | 9465 | 0.77 | tctctttcctgaactatcc**ag**atatcttcctgcctcctgcc |
| 9457 | 9497 | 0.99 | cctcctgcctcctgcttct**ag**ggcggctcaaatgctcattg |
| 9882 | 9922 | 0.79 | tttgtttgttttttgagac**ag**ggtctcactctgtcacccaa |
| 10042 | 10082 | 0.75 | taattttgtgtattttaatag**ag**acggggttttgccttgtt |
| 10237 | 10277 | 0.8 | ccaacacatgccctgtttt**ag**gtaaccatggtagttttgtt |
| 10451 | 10491 | 0.89 | ttttttgtatttttaattt**ag**atgggttttcccaggttgga |
| 10465 | 10505 | 0.82 | aatttagatgggttttccc**ag**gttggacaggctggtcttga |
| 10593 | 10633 | 0.74 | gttttagattttttgtac**ag**agggggtctcactctgttgt |
| 11153 | 11193 | 0.95 | gtggtgctccctctctccc**ag**atcccttgctctgtaggaaa |
| 11169 | 11209 | 0.95 | cccagatcccttgctctgt**ag**gaaaccaacccccaaggcct |
| 12021 | 12061 | 0.96 | cttcagcatcctctcctcc**ag**gaagccaccctggtgcctgg |
| 12123 | 12163 | 0.95 | cccttctgtcctcaccccc**ag**gtccatgtccagggcttagg |
| 12177 | 12217 | 0.71 | tggtgttgactcatctccc**ag**ggatcaaaaaagtcagttgt |
| 12571 | 12611 | 0.95 | aaaggttttctcttataac**ag**atctgcagtatgaaaaacca |
| 12796 | 12836 | 0.82 | cacgccattctcctgcctc**ag**cctcccgagtagctgggact |
| 12864 | 12904 | 0.95 | atttttttgtatttttagtagagac**ag**agtttcactgtgtg |
| 13160 | 13200 | 0.99 | taatttttgttttttttgtag**ag**atggggtctcgctatgtt |
| 13429 | 13469 | 0.97 | attttttttttttttttggagac**ag**ggtcttgctgtcaccc |
| 13591 | 13631 | 0.82 | taatttttgcatttttagt**ag**agacggggtttcatcatgtt |
| 14516 | 14556 | 0.99 | taagattttattttatttt**ag**atttaagaggtacatgtgga |
| 14712 | 14752 | 0.93 | catattttttttttttttg**ag**acggagtcttactctattgc |
| 14874 | 14914 | 0.96 | aatttttgtattttttagt**ag**agatggggtttcaccatgta |
| 15061 | 15101 | 0.75 | tcccatcattcaccctcat**ag**cagctctacaggaaggagct |
| 15669 | 15709 | 0.83 | gtaatcccatcactttggc**ag**gtcgaagcgggaggattgcc |
| 16044 | 16084 | 0.98 | gaagctgtttatttgtttc**ag**attggctgagctgtggggat |
| 16223 | 16263 | 0.71 | gtgtgacttctctgctccc**ag**caatgtcttgggcacccttc |
| 16335 | 16375 | 0.86 | gcttgcaagtcttcctccc**ag**actgccccctgcttctgtgt |
| 16443 | 16483 | 0.77 | tgacccccagtctttctcc**ag**tcgcacctcctctgcacgct |
| 16501 | 16541 | 0.87 | ctctttttcttttttttttgagac**ag**agtcttgctctgtcat |
| 16602 | 16642 | 0.8 | caagtgattctcctgcctc**ag**tcctccctgggactacaggg |
| 16662 | 16702 | 0.78 | taatttttatattttcagtag**ag**tcagggtttcaccatgtt |
| 16795 | 16835 | 0.88 | ggctgtgtgcttctattct**ag**ctccccgactcctgatcccc |
| 16855 | 16895 | 0.86 | cgaaggtctcatcccctcc**ag**gcaggccctccttggcaggg |
| 16986 | 17026 | 0.83 | ctcctaccttggcctccta**ag**gtgctgggattacaagtatg |
| 17522 | 17562 | 0.94 | taccctccttcattgcctt**ag**aggtgaggctggggaacgac |
| 17819 | 17859 | 0.75 | tggggctgtttcgtccact**ag**agcagcacagtcaagaatga |
| 18228 | 18268 | 0.73 | cgtatccattcctcaagcc**ag**gcctctgaggagccctggtg |

| | | | |
|---|---|---|---|
| 18488 | 18528 | 0.72 | taattttggcgtttttagtag**ag**acagggtttcgccatgtt |
| 18662 | 18702 | 0.95 | gcttttctccaccctcttc**ag**actttgttctagtctacaat |
| 18674 | 18714 | 0.77 | cctcttcagactttgttct**ag**tctacaatttccctcctgca |
| 18695 | 18735 | 0.7 | tctacaatttccctcctgc**ag**cttctctctgatgacgtccc |
| 19104 | 19144 | 0.87 | ctttcttgttttcttttgaagac**ag**ggtctcgctcggtcac |
| 20061 | 20101 | 0.96 | tatccccttttataacttt**ag**gtggctgtctttattaaaag |

| HIC2-PI4KA (Donor sites) | | | |
|---|---|---|---|
| Start | End | Score | Exon-Intron |
| 95 | 109 | 0.94 | cactctg**gt**gaggcg |
| 677 | 691 | 0.81 | ggttgcg**gt**gagcca |
| 3255 | 3269 | 0.75 | gatgaag**gt**atcggg |
| 3428 | 3442 | 0.97 | gcatcag**gt**acatcc |

| HIC2-PI4KA (Acceptor sites) | | | |
|---|---|---|---|
| Start | End | Score | Intron-Exon |
| 245 | 285 | 0.76 | gaggccactgccctcttcc**ag**tccccttgccagtctcaggg |
| 263 | 303 | 0.91 | cagtccccttgccagtctc**ag**ggtaccaggtgtctgggtgc |
| 408 | 448 | 0.81 | catatgctgcttcagcctc**ag**acctcggtgggccttaagag |
| 770 | 810 | 0.86 | gtagctgttcccatttcac**ag**ggcactgggggtgctcctga |
| 868 | 908 | 0.98 | ctctccacttccttagttc**ag**gtgcccttgaacctctggcc |
| 906 | 946 | 0.89 | gcctctctgtctggtcctc**ag**ataggagtgggtcctcagat |
| 1043 | 1083 | 0.77 | gctgctcacagcctccttc**ag**agttctgggtcttgaaagca |
| 1138 | 1178 | 0.9 | tgtagccctttcatcctcc**ag**aataaagggaccttcttgag |
| 1442 | 1482 | 0.97 | gggcccctccctttactgt**ag**atgccagtgtggcccctccc |
| 1472 | 1512 | 0.97 | tggcccctccctttactgt**ag**atgtgagcgaggtccctccc |
| 1530 | 1570 | 0.95 | aggtccctccctttactgt**ag**atgccagcgtgacccttccc |
| 1560 | 1600 | 0.98 | tgacccttcccttcactat**ag**atgccagcgaggtccctccc |
| 1590 | 1630 | 0.95 | aggtccctcccttcactgt**ag**atgccagcgaggtccttctc |
| 2015 | 2055 | 0.85 | cctgggctgctcccaccat**ag**gctgcgtcatgcattccctc |
| 2042 | 2082 | 0.98 | tcatgcattccctcttctc**ag**atggccatgccttgcgcctc |
| 2081 | 2121 | 0.9 | tcactcctgcgtctcctcc**ag**ggctcatctcagatgtcccc |
| 2093 | 2133 | 0.83 | ctcctccagggctcatctc**ag**atgtcccctccttgccaggg |
| 2111 | 2151 | 0.74 | cagatgtcccctccttgcc**ag**ggctctccctggccacctgg |
| 2459 | 2499 | 0.87 | ctgtagtgcctcgtcccc**ag**gagtcctcaccccatgggca |
| 2573 | 2613 | 0.77 | cctttgtctccaccctcac**ag**cctccagatggcttctgtgc |
| 2603 | 2643 | 0.93 | ggcttctgtgcctcccagc**ag**gaaggaagacaaggacagag |
| 2684 | 2724 | 0.72 | caccctggcatgtgttccc**ag**ggccttgcccagttcactct |
| 2718 | 2758 | 0.81 | tcactctccctggctcatt**ag**agtggccggggagaacagtg |
| 2870 | 2910 | 0.8 | ggtgtgtgttctactcatt**ag**gcctcgttgccacctctgca |
| 3051 | 3091 | 0.97 | tgcacctcttactctctgc**ag**cacccaggtgtctcccgggc |
| 3317 | 3357 | 0.88 | accctcttctgcccttgcc**ag**ctcagcagcccacagagggg |
| 3627 | 3667 | 0.96 | cgtgttcctttttttaattag**ag**aaaacttgacagtagccc |
