## Supplementary Table.5 for "Evolutionary and functional dynamics of chimeric pseudogenes (φgenes)"

**Supplementary Table 4: Chimeric φgene ORF-associated protein structure-function prediction**

Table 1: Kozak Sequence prediction for chimeric and single parent φgenes

Note: Highlighted cells are most likely coded ORFs after considering score, Stop codon and redundancy

| | ATG from 5' end | Reliability score | Identity to kozak rule A/GXXATGG | ORF length (aa) | Stop codon Found | Is in same frame as parent? | Sequence |
|---|---|---|---|---|---|---|---|
| ANAPC1P1 | **2** | **0.23** | **cXXATGt** | **454** | **Yes** | **Yes** | **MSNFYEERTTKIAARDLQEFVPFGGDHCKHHPNALNLQLRQLQPASELWSSNGAAGFVGSLQEVTIHEKQKESWQLRKGVSEIGEDVDYDEELYVAGNMVIWSKGSKSQALAVYKAFTVDSPVQQALWCDFIISQDKSEKAYSSNEVEKCICILQSSCINMHSIEGKDYIASLPFQVANVWPTKYGLLFERSASSHEVPPGSPRESLPTMFSMLHPLDEITPLVCKSGSLFGSLRVQYVVDHAMKIVFLNTDPSIVMTYDAVQNVHSVWTLRRVKSEEENVVLKFSEHGGTPRNVATSSSLTAHLRSLSKGDSPVISPFRNYSSIHSQSRSTSSPSLHSRSPSISNMAALSRAHSPALGVHSFSGVQRFNISSHNQSPKRHSISHSPNSKSNGFFLAPETEPIVPELCIDHLWTETITNIREKNSQASKVFITSDLCGQISCAFQWSPSSSYAV** |
| | 41 | 0.23 | AXXATGt | 185 | No | | MSNTMPQPSTPLDGVSAPKPLSKLLGSLDEVLLLSPVPELRDSSKLHDSLYNEDCTFQQLGTYIDSIRDPVHNRVTLELSNGSMVRITVPEIATSELVQTCLQTIKFILPKEIAVQMLVKWYNVHSAPGGPSYHSEWNLFVTCLMNVMGFNTDRLAWTRNFDFEGSLSPVIAPKKARPSETGSDD |
| | **36** | **0.2** | **cXXATGt** | **269** | **No** | **Yes** | **MWANFLCFSVESQLQLCCVKFQESNDKTQLIFVSVTNIPAKDAAPVEKIDTMLVLEGSGNLVLYTGVVRVGKVFIPGLPAPSLTMSNTMPQPSTPLDGVSAPKPLSKLLGSLDEVLLLSPVPELRDSSKLHDSLYNEDCTFQQLGTYIDSIRDPVHNRVTLELSNGSMVRITVPEIATSELVQTCLQTIKFILPKEIAVQMLVKWYNVHSAPGGPSYHSEWNLFVTCLMNVMGFNTDRLAWTRNFDFEGSLSPVIAPKKARPSETGSDD** |
| | 40 | 0.13 | AXXATGc | 218 | No | | MLVLEGSGNLVLYTGVVRVGKVFIPGLPAPSLTMSNTMPQPSTPLDGVSAPKPLSKLLGSLDEVLLLSPVPELRDSSKLHDSLYNEDCTFQQLGTYIDSIRDPVHNRVTLELSNGSMVRITVPEIATSELVQTCLQTIKFILPKEIAVQMLVKWYNVHSAPGGPSYHSEWNLFVTCLMNVMGFNTDRLAWTRNFDFEGSLSPVIAPKKARPSETGSDD |
| | 34 | 0.13 | AXXATGG | 108 | Yes | | MAALSRAHSPALGVHSFSGVQRFNISSHNQSPKRHSISHSPNSKSNGFFLAPETEPIVPELCIDHLWTETITNIREKNSQASKVFITSDLCGQISCAFQWSPSSSYAV |
| ANAPC1P2 | **2** | **0.37** | **GXXATGG** | **57** | **Yes** | **Yes** | **MDQCVTVERELEKVLHKFSGYGQLCERGLEELIDYTGGLKHEILQSHGQDAELSGTL** |
| | **7** | **0.25** | **GXXATGa** | **368** | **Yes** | **Yes** | **MNDMNHEVMSLIWSEDLRVQDVRRLLQSAHPVRVNVVQYPELSDHEFIEEKENRLLQLCQRTMALPVGRGMFTLFSYHPVPTEPLPIPKLNLTGRAPPRNTTVDLNSGNIDVPPNMTSWASFHNGVAAGLKIAPASQIDSAWIVYNKPKHAELANEYAGFLMALGLNGHLTKLATLNIHDYLTKGHEMTSIGLLLGVSAAKLGTMDMSITRLLSIRIPALLPPTSTELDVPHNVQVAAVVGIGLVYQGTAHRHTAEVLLAEIGRPPGPEMEYCTDRKSYSLAAGLALGMVCLGHGSNLIGMSDLNVPEQLYQYMVGGHRRFQTGMHREKHKSPSYQIKEGDTINVDVTCPGATLALAMIYLKTNNSVF** |
| | 27 | 0.18 | AXXATGG | 164 | Yes | | MDMSITRLLSIRIPALLPPTSTELDVPHNVQVAAVVGIGLVYQGTAHRHTAEVLLAEIGRPPGPEMEYCTDRKSYSLAAGLALGMVCLGHGSNLIGMSDLNVPEQLYQYMVGGHRRFQTGMHREKHKSPSYQIKEGDTINVDVTCPGATLALAMIYLKTNNSVF |
| | 17 | 0.15 | AXXATGa | 253 | Yes | | MTSWASFHNGVAAGLKIAPASQIDSAWIVYNKPKHAELANEYAGFLMALGLNGHLTKLATLNIHDYLTKGHEMTSIGLLLGVSAAKLGTMDMSITRLLSIRIPALLPPTSTELDVPHNVQVAAVVGIGLVYQGTAHRHTAEVLLAEIGRPPGPEMEYCTDRKSYSLAAGLALGMVCLGHGSNLIGMSDLNVPEQLYQYMVGGHRRFQTGMHREKHKSPSYQIKEGDTINVDVTCPGATLALAMIYLKTNNSVF |
| | 15 | 0.14 | GXXATGt | 298 | Yes | | MFTLFSYHPVPTEPLPIPKLNLTGRAPPRNTTVDLNSGNIDVPPNMTSWASFHNGVAAGLKIAPASQIDSAWIVYNKPKHAELANEYAGFLMALGLNGHLTKLATLNIHDYLTKGHEMTSIGLLLGVSAAKLGTMDMSITRLLSIRIPALLPPTSTELDVPHNVQVAAVVGIGLVYQGTAHRHTAEVLLAEIGRPPGPEMEYCTDRKSYSLAAGLALGMVCLGHGSNLIGMSDLNVPEQLYQYMVGGHRRFQTGMHREKHKSPSYQIKEGDTINVDVTCPGATLALAMIYLKTNNSVF |

| | | | | | | | |
|---|---|---|---|---|---|---|---|
| ANAPC1P3 | **1** | **0.28** | **GXXATGG** | **97** | **No** | **Yes** | **MEYCTDRESYSLAAGLALGMVCLGHGSNLIGMSDLNVPEQLYQYMVGGRRRFQTGMHREKHKSPSYQIKEGDTINVDVTCPGATLALAMIYLKTNNR** |
| | 2 | 0.06 | GXXATGG | 78 | No | | MVCLGHGSNLIGMSDLNVPEQLYQYMVGGRRRFQTGMHREKHKSPSYQIKEGDTINVDVTCPGATLALAMIYLKTNNR |
| | 3 | 0.04 | GXXATGG | 4 | Yes | | MAAI |
| | 4 | 0.04 | GXXATGt | 66 | No | | MSDLNVPEQLYQYMVGGRRRFQTGMHREKHKSPSYQIKEGDTINVDVTCPGATLALAMIYLKTNNR |
| | 5 | 0.04 | tXXATGt | 38 | Yes | | MCLSSSISTWLEDVGAFKQECIGRNINHQVIKSKKEIP |
| ANAPC1P4 | **4** | **0.27** | **cXXATGa** | **634** | **No** | **Yes** | **MKDEDFSQNLSLDSSTLLFTHIPAIFFVLRLVYEELKLNTLMGEGICSLLNFSFSWQDHYYRDYPTLVRTTGQVCTIDPGQTGFMHHPSFFTSEPPSIYQWVSSCLKGEGMPPYPYLPGICERSRLVVLSIALYILGDESSVSDESSQYLTRITVAPQKLQAEQEENRFSFRHSTSVSSLAERLVGWMTNVGFTLRDLETLPFGIALPIRDAIYHCREQPASDWPEAVCLLTGRQDLSKQACEGNLPKGKSVLSSDVPSGTETEEEDDGMNDMNHEVMSLIWSEDLRVQDVRRLLQSAHPVRVNVVQYPELSDHEFIEEKENRLLQLCQRTMALPVGRGMFTLFSYHPVPTEPLPIPKLNLTGRAPPRNTTVDLNSGNIDVPPNMTSWASFHNGVAAGLKIAPASQIDSAWIVYNKPKHAELANEYAGFLMALGLNGHLTKLATLNIHDYLTKGHEMTSIGLLLGVSAAKLGTMDMSITRLLSIRIPALLPPTSTELDVPHNVQVAAVVGIGLVYQGTAHRHTAEVLLAEIGRPPGPEMEYCTDRKSYSLAAGLALGMVCLGHGSNLIGMSDLNVPEQLYQYMVGGHRRFQTGMHREKHKSPSYQIKEGDTINVDVTCPGATLALAMIYLKTNN** |
| | 36 | 0.18 | AXXATGG | 161 | No | | MDMSITRLLSIRIPALLPPTSTELDVPHNVQVAAVVGIGLVYQGTAHRHTAEVLLAEIGRPPGPEMEYCTDRKSYSLAAGLALGMVCLGHGSNLIGMSDLNVPEQLYQYMVGGHRRFQTGMHREKHKSPSYQIKEGDTINVDVTCPGATLALAMIYLKTNN |
| | 26 | 0.15 | AXXATGa | 250 | No | | MTSWASFHNGVAAGLKIAPASQIDSAWIVYNKPKHAELANEYAGFLMALGLNGHLTKLATLNIHDYLTKGHEMTSIGLLLGVSAAKLGTMDMSITRLLSIRIPALLPPTSTELDVPHNVQVAAVVGIGLVYQGTAHRHTAEVLLAEIGRPPGPEMEYCTDRKSYSLAAGLALGMVCLGHGSNLIGMSDLNVPEQLYQYMVGGHRRFQTGMHREKHKSPSYQIKEGDTINVDVTCPGATLALAMIYLKTNN |
| | 9 | 0.14 | GXXATGc | 524 | No | | MPPYPYLPGICERSRLVVLSIALYILGDESSVSDESSQYLTRITVAPQKLQAEQEENRFSFRHSTSVSSLAERLVGWMTNVGFTLRDLETLPFGIALPIRDAIYHCREQPASDWPEAVCLLTGRQDLSKQACEGNLPKGKSVLSSDVPSGTETEEEDDGMNDMNHEVMSLIWSEDLRVQDVRRLLQSAHPVRVNVVQYPELSDHEFIEEKENRLLQLCQRTMALPVGRGMFTLFSYHPVPTEPLPIPKLNLTGRAPPRNTTVDLNSGNIDVPPNMTSWASFHNGVAAGLKIAPASQIDSAWIVYNKPKHAELANEYAGFLMALGLNGHLTKLATLNIHDYLTKGHEMTSIGLLLGVSAAKLGTMDMSITRLLSIRIPALLPPTSTELDVPHNVQVAAVVGIGLVYQGTAHRHTAEVLLAEIGRPPGPEMEYCTDRKSYSLAAGLALGMVCLGHGSNLIGMSDLNVPEQLYQYMVGGHRRFQTGMHREKHKSPSYQIKE |
| | 24 | 0.14 | GXXATGt | 295 | No | | MFTLFSYHPVPTEPLPIPKLNLTGRAPPRNTTVDLNSGNIDVPPNMTSWASFHNGVAAGLKIAPASQIDSAWIVYNKPKHAELANEYAGFLMALGLNGHLTKLATLNIHDYLTKGHEMTSIGLLLGVSAAKLGTMDMSITRLLSIRIPALLPPTSTELDVPHNVQVAAVVGIGLVYQGTAHRHTAEVLLAEIGRPPGPEMEYCTDRKSYSLAAGLALGMVCLGHGSNLIGMSDLNVPEQLYQYMVGGHRRFQTGMHREKHKSPSYQIKEGDTINVDVTCPGATLALAMIYLKTNN |
| ANAPC1P5 | **1** | **0.43** | **AXXATGt** | **185** | **No** | **Yes** | **MSNTMPQPSTSLDGVSAPKPLSKLLGSLDEVLLLFPVPELRDSSKLHDSLYNEDCTFQQLGTYIDSIRDPVHNRVTLELSNGSMVRITVPEIATSELVQTCLQTIKFILPKEIAVQMLVKWYNVHSAPVGPSYHSEWNLFVTCLMNVMGFNTDRLAWTRNFDFEGSLSPVIAPKKARPSETGSDD** |
| | 2 | 0.11 | AXXATGc | 181 | No | | MPQPSTSLDGVSAPKPLSKLLGSLDEVLLLFPVPELRDSSKLHDSLYNEDCTFQQLGTYIDSIRDPVHNRVTLELSNGSMVRITVPEIATSELVQTCLQTIKFILPKEIAVQMLVKWYNVHSAPVGPSYHSEWNLFVTCLMNVMGFNTDRLAWTRNFDFEGSLSPVIAPKKARPSETGSDD |
| | 7 | 0.09 | tXXATGG | 102 | No | | MVKITVPEIATSELVQTCLQTIKFILPKEIAVQMLVKWYNVHSAPVGPSYHSEWNLFVTCLMNVMGFNTDRLAWTRNFDFEGSLSPVIAPKKARPSETGSDD |
| | 9 | 0.08 | cXXATGc | 69 | No | | MLVKWYNVHSAPVGPSYHSEWNLFVTCLMNVMGFNTDRLAWTRNFDFEGSLSPVIAPKKARPSETGSDD |
| | 5 | 0.04 | AXXATGa | 25 | Yes | | MRIVLSNSLELTLILSEILSITESP |

| | | | | | | | |
|---|---|---|---|---|---|---|---|
| ANAPC1P6 | **3** | **0.15** | **GXXATGa** | **201** | **Yes** | **Yes** | **MKIVFLNTDPSIVTTYDAVQNVHSVWTLRRVKSEEENVVLKFSEQGGTPQNVATSSSLTAHLRSLSKGDSPVISPFRNYSSIHSQSRSTSSPSLHSRSPSISNMAALSRAHSPALGVHSFSGVQRFNISSHNQSPKRHSISHSPNSNSNDSFLAPETEPIVPELCIDHLWTEMITNIREKNSQASKVFITSDLCGQNSCAF** |
| | 9 | 0.12 | AXXATGG | 98 | Yes | | MAALSRAHSPALGVHSFSGVQRFNISSHNQSPKRHSISHSPNSNSNDSFLAPETEPIVPELCIDHLWTEMITNIREKNSQASKVFITSDLCGQNSCAF |
| | 12 | 0.11 | cXXATGt | 69 | No | | MWAKPLCPLVESQLHLRCVKPQESNDKTQLIPDSVTNIPAKDAAPVEKIDTMLVLEGSGNLVLYTGVVP |
| | 8 | 0.06 | AXXATGt | 21 | Yes | | MWPLAAPSQHISEASPKEIPL |
| | 5 | 0.04 | AXXATGc | 23 | Yes | | MLFKMCILCGLSGESNQRKRMLF |
| RMND5A φ | **1** | **0.97** | **AXXATGG** | **57** | **Yes** | **Yes** | **MDQCVTVERELEKVLHKFSGYGQLCERGLEELIDYTGGLKHEILQSHGQDAELSGTL** |
| | 2 | 0.06 | AXXATGc | 11 | Yes | | MLNYQGHFDLF |
| GTF2IP1 | **28** | **0.3** | **cXXATGa** | **145** | **Yes** | | **MMIMKDSRKLKKLSLREQVNDLFSRKFGEAIGMGFPVKVPYRKITINPGCVVVDGMPPGVSFKAPSYLEISSMRRILDSAEFIKFTVIRPFPGLVINNQLVDQSESEGPVIQESAEPSQLEVPATEEIKETDGSSQIKQEPDPTW** |
| | **15** | **0.28** | **AXXATGG** | **404** | **Yes** | **Yes** | **MVDQLFCKKFAEALGSTEAKAVPYQKFEAHPNDLYVEGLPENIPFRSPSWYGIPRLEKIIQVGNRIKFVIKRPELLTHSTTEVTQPRTNTPVKEDWNVRITKLRKQVEEIFNLKFAQALGLTEAVKVPYPVFESNPEFLYVEGLPEGIPFRSPTWFGIPRLERIVRGSNKIKFVVKKPELVISYLPPGMASKINTKALQSPKRPRSPGSNSKVPEIEVTVEGPNNNNPQTSAVRTPTQTNGSNVPFKPRGREFSFEAWNAKITDLKQKVENLFNEKCGEALGLKQAVKVPFALFESFPEDFYVEGLPEGVPFRRPSTFGIPRLEKILRNKAKIKFIIKKPEMFETAIKESTSSKSPPRKINSSPNVNTTASGVEDLNIIQVTIPDDDNERLSKVEKAVAKRTSE** |
| | 21 | 0.17 | GXXATGG | 216 | Yes | | MASKINTKALQSPKRPRSPGSNSKVPEIEVTVEGPNNNNPQTSAVRTPTQTNGSNVPFKPRGREFSFEAWNAKITDLKQKVENLFNEKCGEALGLKQAVKVPFALFESFPEDFYVEGLPEGVPFRRPSTFGIPRLEKILRNKAKIKFIIKKPEMFETAIKESTSSKSPPRKINSSPNVNTTASGVEDLNIIQVTIPDDDNERLSKVEKAVAKRTSE |
| | 30 | 0.14 | AXXATGa | 142 | Yes | | MKDSRKLKKLSLREQVNDLFSRKFGEAIGMGFPVKVPYRKITINPGCVVVDGMPPGVSFKAPSYLEISSMRRILDSAEFIKFTVIRPFPGLVINNQLVDQSESEGPVIQESAEPSQLEVPATEEIKETDGSSQIKQEPDPTW |
| | **1** | **0.12** | **cXXATGa** | **37** | **Yes** | | **MSRRGRGAGGRWNSTSWSTGCKLPASPRRVSRCSPTA** |
| GTF2IP2 | **1** | **0.23** | **tXXATGG** | **82** | **Yes** | **Yes** | **MAQVLVSTLSIEDEDSYESRMVVTFLMSALESTFKELAKFKAKVACITEYKADLFAFRTEGQRTQFFQYQKGFSSRFCEILC** |
| | 5 | 0.08 | GXXATGc | 49 | Yes | | MHKIKSTTQVNQMSVDVVEMATLGKKQLRTISAFARESFTKTHRGTCTI |
| | 10 | 0.08 | AXXATGc | 56 | Yes | Yes | MLQDQSALIVQGLPEGVAFKHPENYDLATLKWILENTAGISFIINRSFLEPKKPLD |
| | 4 | 0.04 | cXXATGt | 56 | Yes | | MSALESTFKELAKFKAKVACITEYKADLFAFRTEGQRTQFFQYQKGFSSRFCEILC |
| | 2 | 0.04 | AXXATGa | 10 | Yes | | MRTPTRAGWW |
| GTF2IP3 | **2** | **0.26** | **AXXATGa** | **104** | **No** | **Yes** | **MKSTTQVNQMSVDAVEVATLGKTVEDYFCFCYGKALQKPTEVPVPYEKMRQDQSALIVQGLPEDVAFKHPENYDLATLKWILENTAGISFIIKRSFLEPKKHLG** |
| | 3 | 0.11 | cXXATGa | 95 | No | | MSVDAVEVATLGKTVEDYFCFCYGKALQKPTEVPVPYEKMRQDQSALIVQGLPEDVAFKHPENYDLATLKWILENTAGISFIIKRSFLEPKKHLG |
| | 1 | 0.05 | GXXATGc | 27 | Yes | | MHKNEIYNPSESDECRCCRSGNTRKNS |
| | 4 | 0.04 | tXXATGc | 2 | Yes | | ML |
| | 5 | 0.04 | GXXATGG | 24 | Yes | | MGKLYKNPQRYLYHMRRCDKTSQL |

| | | | | | | | |
|---|---|---|---|---|---|---|---|
| GTF2IP4 | **15** | **0.29** | **AXXATGG** | **530** | **Yes** | **Yes** | **MVDQLFCKKFAEALGSTEAKAVPYQKFEAHPNDLYVEGLPENIPFRSPSWYGIPRLEKIIQVGNRIKFVIKRPELLTHSTTEVTQPRTNTPVKEDWNVRITKLRKQVEEIFNLKFAQALGLTEAVKVPYPVFESNPEFLYVEGLPEGIPFRSPTWFGIPRLERIVHGSNKIKFVVKKPELVISYLPPGMASKINTKALQSPKRPRSPGSNSKVPEIEVTVEGPNNNNPQTSAVRTPTQTNGSNVPFKPRGREFSFEAWNAKITDLKQKVENLFNEKCGEALGLKQAVKVPFALFESFPEDFYVEGLPEGVPFRRPSTFGIPRLEKILRNKAKIKFIIKKPEMFETAIKESTSSKSPPRKINSSPNVNTTASGVEDLNIIQVTIPDDDNERLSKVEKARQLREQVNDLFSRKFGEAIGMGFPVKVPYRKITINPGCVVVDGMPPGVSFKAPSYLEISSMRRILDSAEFIKFTVIRPFPGLVINNQLVDQSESKGPVIQESAEPSQLEVPATEEIKETDGSSQIKQEPDPTW** |
| | 26 | 0.21 | GXXATGt | 189 | Yes | | MFETAIKESTSSKSPPRKINSSPNVNTTASGVEDLNIIQVTIPDDDNERLSKVEKARQLREQVNDLFSRKFGEAIGMGFPVKVPYRKITINPGCVVVDGMPPGVSFKAPSYLEISSMRRILDSAEFIKFTVIRPFPGLVINNQLVDQSESKGPVIQESAEPSQLEVPATEEIKETDGSSQIKQEPDPTW |
| | **21** | **0.18** | **GXXATGG** | **342** | **Yes** | | **MASKINTKALQSPKRPRSPGSNSKVPEIEVTVEGPNNNNPQTSAVRTPTQTNGSNVPFKPRGREFSFEAWNAKITDLKQKVENLFNEKCGEALGLKQAVKVPFALFESFPEDFYVEGLPEGVPFRRPSTFGIPRLEKILRNKAKIKFIIKKPEMFETAIKESTSSKSPPRKINSSPNVNTTASGVEDLNIIQVTIPDDDNERLSKVEKARQLREQVNDLFSRKFGEAIGMGFPVKVPYRKITINPGCVVVDGMPPGVSFKAPSYLEISSMRRILDSAEFIKFTVIRPFPGLVINNQLVDQSESKGPVIQESAEPSQLEVPATEEIKETDGSSQIKQEPDPTW** |
| | **2** | **0.12** | **AXXATGG** | **38** | **Yes** | | **MELHILEHRLQVASVAKESIPLFTYGLIKLAFLSSKTR** |
| | **1** | **0.12** | **cXXATGa** | **37** | **Yes** | | **MSRRGRGAGGRWNSTSWSTGCKLPASPRRVSRCSPTA** |
| GTF2IP5 | **1** | **0.8** | **AXXATGa** | **31** | **Yes** | | **MMMILHLLRDQRPTNYHSHQFRNLPMMGNGK** |
| | **11** | **0.15** | **AXXATGG** | **100** | **Yes** | **Yes** | **MVDQLFCKKTEALGSTEAKVLLYQKFEGHANDLYAEGLPENIPFRSPSWYGIPRLENIIQVGNRIKFLIKSPELLTHNTTVVTQPRTNTTGLKLLDSLRQ** |
| | 9 | 0.09 | GXXATGc | 89 | Yes | | MPESLNEERWWISFSAKKLKPWGALKPRFYCTKNLKAMRMICMRKDYQKTFLSEVPRGMESQGWKTSFKWAIELNFLLKVQNFLLTIQL |
| | 4 | 0.04 | cXXATGa | 6 | Yes | | MMGNGK |
| | 5 | 0.04 | AXXATGG | 5 | Yes | | MGNGK |
| GTF2IP6 | **1** | **0.47** | **AXXATGG** | **15** | **Yes** | | **MVVTFLMSALETGQV** |
| | **5** | **0.25** | **cXXATGa** | **91** | **Yes** | **Yes** | **MSVDAVEVATLRKNTLQKPTEVTVPYEKMLQDQSALIVQGLPEGVAFKHPENYYLATLKWILENTAGISFIIKRHFPGSGSCDSKGRIRRS** |
| | 3 | 0.04 | cXXATGt | 31 | Yes | | MSLPSELKDRGHNFFNTRKDFQTDFVKYCKH |
| | 4 | 0.04 | GXXATGc | 2 | Yes | | MH |
| | 2 | 0.04 | cXXATGt | 9 | Yes | | MSALETGQV |
| GTF2IP7 | 1 | 0.04 | tXXATGa | 2 | Yes | | MT |
| | 2 | 0.04 | tXXATGG | 38 | Yes | | MGVIKSNLLVKPELVISYLPPGVANKINTKASQFLKRP |
| | 3 | 0.04 | AXXATGa | 3 | Yes | | MKP |
| | 4 | 0.04 | GXXATGa | 12 | No | | MKRFLKLRSLWK |
| GTF2IP8 | 5 | 0.09 | GXXATGt | 52 | Yes | | MCKIKSTTQVNQMSVDVVEMATLGKKQLRTISAFAVLKGESFTKTHRGTCTI |
| | 1 | 0.08 | AXXATGG | 20 | Yes | | MAQVTVSTLAVEDEESSAGW |
| | 10 | 0.07 | AXXATGc | 52 | No | | MLQDQSALIVQGLPEGVAFKHPENYDLATLEWILEYTAGISFIIKILRAKEA |
| | 4 | 0.04 | cXXATGt | 12 | Yes | | MSALESIRTGQV |
| | 2 | 0.04 | AXXATGa | 16 | Yes | | MRSPQQDGSDIPHVSS |
| GTF2IP9 | 1 | 0.04 | tXXATGt | 20 | Yes | | MFRRFARQDSLLKPSWFGIP |
| | 2 | 0.04 | tXXATGa | 2 | Yes | | MT |
| | 3 | 0.04 | GXXATGa | 12 | No | | MKRFLKLRSLWK |
| GTF2IP11 | 1 | 0.06 | cXXATGt | 22 | Yes | | MCRRLARGDSLSKPYLLWNSMT |
| | 2 | 0.04 | tXXATGa | 2 | Yes | | MT |
| | 3 | 0.04 | AXXATGa | 7 | Yes | | MKFVVKK |

| | | | | | | | |
|---|---|---|---|---|---|---|---|
| | 4 | 0.04 | GXXATGG | 13 | Yes | | MVAHTCNPSTLGG |
| | 5 | 0.04 | GXXATGG | 13 | Yes | | MASKINTKQSPKR |
| GTF2IP12 | **1** | **0.46** | **AXXATGa** | **24** | **Yes** | | **MMIIPHLLGHKDQRATTATSPRTR** |
| | 4 | 0.24 | AXXATGa | 86 | No | | MREFNSKWNNCIAHLRKQIEELSERKYRAVKAKGPVMIPYPFFQSHVEDFYVEGLPKGIFFRLSTNGIPGLERILLSTERIFLLLK |
| | 3 | 0.05 | cXXATGc | 6 | Yes | | MLGNGK |
| | 2 | 0.04 | AXXATGa | 23 | Yes | | MIIPHLLGHKDQRATTATSPRTR |
| | 5 | 0.04 | tXXATGG | 2 | Yes | | ME |
| GTF2IP13 | 2 | 0.08 | AXXATGa | 32 | Yes | Yes | MREFNSEKWNNRIADLRKQIEELSERKYDMNL |
| | 7 | 0.06 | GXXATGa | 51 | No | | MIPYPFFQSHVEDFYVEGLPKGRLSTNGIPGLERILLSKERIFLLLKGKIL |
| | 3 | 0.04 | GXXATGG | 2 | Yes | | ME |
| | 4 | 0.04 | AXXATGa | 2 | Yes | | MT |
| | 5 | 0.04 | GXXATGa | 3 | Yes | | MNL |
| GTF2IP14 | 14 | 0.16 | tXXATGa | 91 | No | | MKKDGGSAFLQKNLALGSTEAKALLYQKFEGHANDLYVEGLPENIPFRSPSWYGIARLENIIQVGNRIKFLIKSQNFLLTINCSYSAKNKY |
| | **3** | **0.13** | **AXXATGa** | **38** | **Yes** | | **MILHLLRDQRPTNYHSWQAVTHSLFRSLPMMGSSGTCQ** |
| | **15** | **0.1** | **AXXATGG** | **77** | **Yes** | **Yes** | **MVDQLFCKKIWPWGALKPRLYCTKNLKAMRMICMWKDYQKTFLSEVPRGMESQGWKTSFKWAIELNFLLKARTSYSQ** |
| | 1 | 0.06 | AXXATGa | 40 | Yes | | MMMILHLLRDQRPTNYHSWQAVTHSLFRSLPMMGSSGTCQ |
| | 5 | 0.04 | AXXATGG | 8 | Yes | | MGSSGTCQ |
| GTF2IP18 | 4 | 0.07 | AXXATGa | 32 | Yes | | MREFNSEKWNHRIADLRKQTEELSERKYDMNL |
| | 1 | 0.05 | AXXATGa | 36 | Yes | | MMIIPHLLRDKDQRATTATSPRTRQCWGTENEGVQL |
| | 3 | 0.04 | cXXATGc | 6 | Yes | | MLGNGK |
| | 2 | 0.04 | AXXATGa | 35 | Yes | | MIIPHLLRDKDQRATTATSPRTRQCWGTENEGVQL |
| | 5 | 0.04 | GXXATGG | 6 | Yes | | MESSYR |
| GTF2IP20 | **4** | **0.25** | **AXXATGa** | **89** | **No** | **Yes** | **MREFNSEKWNHRIADLRKQTEELSERKYARAVKAKGLVMIPYPFFQSHVEDFYVEGLPKGFLGRLSTNGIPGLERMLLSKERIFLLLKG** |
| | 1 | 0.05 | tXXATGa | 36 | Yes | | MMIIPHLLRDKDQRATTATSPRTRQCWGTENEGVQL |
| | 3 | 0.04 | cXXATGc | 6 | Yes | | MLGNGK |
| | 2 | 0.04 | AXXATGa | 35 | Yes | | MIIPHLLRDKDQRATTATSPRTRQCWGTENEGVQL |
| | 5 | 0.04 | GXXATGG | 6 | Yes | | MESSYR |
| GTF2IP22 | 1 | 0.04 | cXXATGa | 10 | Yes | | MRRCYKTSQL |
| | 2 | 0.04 | AXXATGc | 29 | Yes | | MLQDQSALIVQGLPEGVALNTLRIMILQP |
| | 3 | 0.04 | AXXATGa | 5 | Yes | | MILQP |
| | 4 | 0.04 | GXXATGG | 13 | No | | MDFGEHSRGFSYY |
| GTF2IP23 | 3 | 0.05 | AXXATGa | 3 | Yes | | MKP |
| | 2 | 0.04 | GXXATGG | 21 | Yes | | MANKINTKGRRLPALQSPKKP |
| | 1 | 0.04 | tXXATGa | 2 | Yes | | MT |
| | 4 | 0.04 | GXXATGa | 16 | No | | MKSFLTLRSLWKVRAS |
| ENSG000002344 | **13** | **0.36** | **GXXATGa** | **103** | **Yes** | | **MKMKVLERDNMVRELESALDHLKLQCDRRLTLQQKEHEQKMQLLLHHFKEQDGEGIMETFKTYEDKIQQLEKDLYFYKKTSRDHKKKLKELVGEAIRRQLASS** |
| | **21** | **0.28** | **GXXATGc** | **131** | **Yes** | **Yes** | **MLSEELKWASRPESMKLSGREREMDSSASSLRTQPNPQKLWEDIPELPPIHSSLAPPSGHMLGNENKTETDDNQFTKSQSTVIPNSGCGKCGTTSWCHTCKTVSKRITSNFRLGTIIATFQSWSWHWINGC** |
| | **5** | **0.24** | **tXXATGG** | **108** | **Yes** | | **MEQDFSEKNVQLQTSTAEEKTKISEQVEVLQKEKDQLQKRRHIFKMKNLKMVECYHLKKNMFFSNLKKGLKLWKLQLNTRMKVSRIARSHLEHHSITSLVVKQMSWKN** |
| | 14 | 0.12 | AXXATGa | 101 | Yes | | MKVLERDNMVRELESALDHLKLQCDRRLTLQQKEHEQKMQLLLHHFKEQDGEGIMETFKTYEDKIQQLEKDLYFYKKTSRDHKKKLKELVGEAIRRQLASS |
| | 26 | 0.12 | GXXATGa | 81 | No | | MRIKQKQMIISLQNLSRLSSQIQVVGNVGRLHGVTPVKLCRKELRQISALELSLRRSSLGVGIGSMAADSIEVSRKPRDLK |

| | | | | | | | |
|---|---|---|---|---|---|---|---|
| ENSG000001651 | **7** | **0.27** | **AXXATGa** | **160** | **Yes** | **Yes** | **MKKETQCVLAADEVVFNQKELEVKELKNQVQ<br>MMVQENKGHAVSLKEAQKVNRLQNEKIIEQQL<br>LVDQLSEELTKLNLSVTSSAKENCGDGPDARIP<br>EKRPYTVPFDTHLGHYIYIPSRQDSRRGNHLQG<br>PHKSAYVLSGSNICWISNTKSDAVGSRRRTR** |
| | **13** | **0.25** | **cXXATGc** | **111** | **Yes** | | **MPGSLKRDHILYHLILIWGIIFISHQDKIPGGGITC<br>KVHTSPPMYSLDRIFAGFRTQSQMLLDHVEER<br>DEVLHCQFSDNSDDEESEGQEKSGTRCRSRS<br>WIQKPDSVPLLN** |
| | 23 | 0.18 | AXXATGa | 155 | No | | MKELTVNIKMKEDLIKELIKTGNDAKSVSKQYTL<br>KVTKLEHDAEQAKVELTETQKQLQELENKDLSD<br>VAMKVKLQKEFRKKVDAAKLRVQVLQKKQQDS<br>KKLASLSIQNEKRANELEQSVDHMKYQKIQLQR<br>KLQEENEKRKQLDAVIKRDQQKI |
| | 24 | 0.16 | AXXATGa | 146 | No | | MKEDLIKELIKTGNDAKSVSKQYTLKVTKLEHDA<br>EQAKVELTETQKQLQELENKDLSDVAMKVKLQK<br>EFRKKVDAAKLRVQVLQKKQQDSKKLASLSIQN<br>EKRANELEQSVDHMKYQKIQLQRKLQEENEKR<br>KQLDAVIKRDQQKI |
| | 20 | 0.12 | tXXATGG | 214 | No | | MDSEARLCSLVELSDTQDETQKSDSENEDLKID<br>CLQESQELNLQKLKNSERILTEAKQKMRELTVNI<br>KMKEDLIKELIKTGNDAKSVSKQYTLKVTKLEHD<br>AEQAKVELTETQKQLQELENKDLSDVAMKVKLQ<br>KEFRKKVDAAKLRVQVLQKKQQDSKKLASLSIQ<br>NEKRANELEQSVDHMKYQKIQLQRKLQEENEK<br>RKQLDAVIKRDQQKI |
| ENSG000002909 | **7** | **0.27** | **AXXATGa** | **160** | **Yes** | **Yes** | **MKKETQCVLAADEVVFNQKELEVKELKNQVQ<br>MMVQENKGHAVSLKEAQKVNRLQNEKIIEQQL<br>LVDQLSEELTKLNLSVTSSAKENCGDGPDARIP<br>EKRPYTVPFDTHLGHYIYIPSRQDSRRGNHLQG<br>PHKSAYVLSGSNICWISNTKSDAVGSRRRTR** |
| | **13** | **0.25** | **cXXATGc** | **111** | **Yes** | | **MPGSLKRDHILYHLILIWGIIFISHQDKIPGGGITC<br>KVHTSPPMYSLDRIFAGFRTQSQMLLDHVEER<br>DEVLHCQFSDNSDDEESEGQEKSGTRCRSRS<br>WIQKPDSVPLLN** |
| | **23** | **0.17** | **AXXATGa** | **170** | **Yes** | | **MRELTVNIKMKEDLIKELIKTGNDAKSVSKQYT<br>LKVTKLEHDAEQAKVELTETQKQLQELENKDL<br>SDVAMKVKLQKEFRKKVDAAKLRVQVLQKKQ<br>QDSKKLASLSIQNEKRANELEQSVDHMKYQKI<br>QLQRKLQEENEKRKQLDAVIKRDQQKIKVILSYI<br>PAKYNMKC** |
| | 24 | 0.15 | AXXATGa | 161 | Yes | | MKEDLIKELIKTGNDAKSVSKQYTLKVTKLEHDA<br>EQAKVELTETQKQLQELENKDLSDVAMKVKLQK<br>EFRKKVDAAKLRVQVLQKKQQDSKKLASLSIQN<br>EKRANELEQSVDHMKYQKIQLQRKLQEENEKR<br>KQLDAVIKRDQQKIKVILSYIPAKYNMKC |
| | **4** | **0.14** | **tXXATGa** | **73** | **Yes** | | **MTSVVEGQKGIILRAIQEIFQSISEHPSIDFNVKV<br>SYIEVYKEDLRDLLELETSMKDLHIRKDEKGNT<br>VCACC** |
| LOC389765 | **19** | **0.25** | **cXXATGc** | **111** | **Yes** | | **MPGSLKRDHILYHLILIWGIIFISHQDKIPGGGITC<br>KVHTSPPMYSLDRIFAGFRTQSQMLLDHVEER<br>DEVLHCQFSDNSDDEESEGQEKSGTRCRSRS<br>WIQKPDSVPLLN** |
| | **12** | **0.23** | **AXXATGa** | **87** | **Yes** | | **MKKETQCVLAADEVVFNQKELEVKELKNQVQ<br>MMVQENKGHAVSLKEAQKVNRLQDLILLLRLV<br>CAGTITAHCSLDLPGSSDPPASAS** |
| | 8 | 0.09 | GXXATGc | 54 | Yes | Yes | MQLFLPMDKLDLGRHTPLEGAILLQLWRAKRVS<br>FFELFKKYFKASLNILALTLM |
| | 26 | 0.07 | tXXATGG | 80 | No | | MDSEARLCSLVELSDTQDETQKSDSENEDLKID<br>CLQESQELNLQKLKNSERILTEAKQKMRELTVNI<br>KMKEDLIKELIKT |
| | 17 | 0.05 | tXXATGc | 30 | Yes | | MPPRPANFLIFLWRRGLTVLLKLVSNSWAQ |
| HYDINP1 | 2 | 0.22 | AXXATGa | 61 | No | | MTPPTSALGACFVFSPKEGIIEPSGVQAIQISFSS<br>TILGNFEEEFLVNVNGSPEPVKLTIR |
| | 1 | 0.04 | tXXATGc | 5 | Yes | | MLSST |
| | 3 | 0.04 | tXXATGt | 8 | Yes | | MSMGHLSL |
| | 4 | 0.04 | tXXATGG | 6 | Yes | | MGHLSL |
| ENSG000002762 | 1 | 0.04 | tXXATGa | 37 | Yes | | MKKKKTLLIKFEPSYRNDLNNWVAEEILAIKYVE<br>HPQ |
| | 2 | 0.04 | AXXATGt | 22 | Yes | | MWNTLSRQPGPAWMSALPQPQL |
| | 3 | 0.04 | tXXATGG | 12 | Yes | | MDECITPTSALR |
| | 4 | 0.04 | tXXATGa | 9 | Yes | | MSALPQPQL |
| | 5 | 0.04 | GXXATGa | 17 | Yes | | MMELDFGCILNNTEVIR |
| ENSG000002597 | 4 | 0.07 | AXXATGt | 21 | Yes | | MSFIAKGPIQDHILHPITMSL |
| | 7 | 0.07 | GXXATGa | 22 | Yes | | MTLIFLKITGWSFYFVLQFGIA |
| | 3 | 0.05 | cXXATGc | 37 | Yes | | MPNNVLYSKGPNPGSHSAPNYHVSLVSFNLEQ<br>EEGIS |
| | 2 | 0.04 | AXXATGc | 20 | Yes | | MQKGFQSNTRTSIKYLSHAQ |
| | 5 | 0.04 | AXXATGt | 3 | Yes | | MSL |

| | | | | | | |
|---|---|---|---|---|---|---|
| ENSG000002598 | 5 | 0.07 | AXXATGt | 41 | Yes | | MSLVAGTRLHPSLRLKRPRRKVIMASDSEAKLPGFKSQLCH |
| | 2 | 0.04 | tXXATGG | 18 | Yes | | MEGGLASSYQNKRNCHGN |
| | 3 | 0.04 | GXXATGG | 12 | Yes | | MVINNLLLWESK |
| | 4 | 0.04 | AXXATGc | 12 | Yes | | MLGDIIVRSCPW |
| | 1 | 0.04 | GXXATGt | 1 | Yes | | M |
| HYDIN2 | **112** | **0.7** | **GXXATGa** | **635** | **Yes** | | **MIITIILEMRKLECSTVPPLTKVDVKMETIERKISVREQTMSEKEELNKKKRNMGDVSMHGLPLVQDQEDSEGDISKDPDKQLAQKFKTYELTLKDVQNILMYWDRKQGVQLPPAGMEEAPHEPDDQRQVPSGGRRGRKDRERERLEKERLEREKAERERLEKLRALEERSDWEGEGEEDHEEKKEKDLGVPFLNIQTPDFEGLSWKQALESDRLPKGEQAQEEQTSSSKGGKQKMKEKIDQVFESQKDKRHMALNRKVLSGEPAGTISQLSDTDLDNFNGQHSQEKFTRLNHFRWIVPANGEVTLQVHFSSDEFGNFDQTFNFEILGTCCQYQLYCRGICTYPYICQDPKVVFPQRKMDMKTNEVIFKKYVMSTETYYFGPLLCGKSRDKYKSSLFPGNMETLTILNTSLMVVEASFYFQNDVKANTYFLEPNTMVLKPNEKQILNVWAYPTSVGVFEDSIVCCINDNPEPAIFQLSCQGIRPELELEPRQLHFDRLLLHRQESRVVLLRNVTLLPVAWRITSLEHLGDDFTVSLMQGTIPPEAEYGLHLYFQPTKPVNIKKAIRLEVLDAENLLGVVQIENIMVFAEAYDIALDITFPKGAEGGLDFGIVRVTEEAKQPLQLKNRGKYEIAFR** |
| | 62 | 0.69 | AXXATGG | 222 | Yes | | MELYPGQAIDVILEGYSATPRKPNSILKPDYQPLAVKNISTLPVNLLLSTSGPFFICETDKSLLPATPEPIKLEIDEEKNLLIKFDPSYRNDLNNWVAEEILAIKYVEHPQIDSLDLRGEVHYPNLSFETKELDFGCILNDTELIRYVTITNCSPLVVKFRWFFLVNDEENQIRFVTLPKKPYSAPLSQMESIPATSEAASPPAILVTVESPEMDLNDFVKK |
| | **102** | **0.52** | **AXXATGG** | **137** | **Yes** | | **MAQDYAAMKAREKAKKEQEERKHKGALEKEKEGLQNMDEEEYDALTEEEKLTFDRGIQQALRERKKREQERLAKEMQEKKLQQELERQKEEDELKRRVKKGKQGPIKEEPPMKKSQAANKQLLCAEFLLCDTPCWEL** |
| | **97** | **0.44** | **tXXATGG** | **107** | **Yes** | | **MEEVGEVENNPVSKAIARHLGIDISAEGRLAKNRKGIAIIIHGTPLSGKSANAVSVAKYYNAACLSIDSIVLEAVANSNNIPGIRACELCIRAAIEQSMKEGEEAAE** |
| | **26** | **0.43** | **tXXATGc** | **960** | **Yes** | | **MQLEVEFEPQSVGDHSGRLIVCYDTGEKVFVSLYGAAIDMNIRLDKNSLTIEKTYISLANQRTITIHNRSNIIAHFLWKVFATQQEEDREKYRACDDLIKEEKDETDEFFEECITDPLLREHLSVLSRTFANQRRLVQGDSKLFFNNVFTVEPLEGDVWPNSSAEITVYFNPLEAKLYQQTIYCDILGREIRLPLRIKGEGMGPKIHFNFELLDIGKVFTGSAHCYEAILYNKDSIDALFNMTPPTSALGACFVFSPKEGIIEPSGVQAIQISFSSTILGNFEEEFLVNVNGSPEPVKLTIRGCVIGPTFHFNVPALHFGDVSFGFPHTLICSLNNTSLIPMTYKLRIPGDGLGHKSISYCEQHVDYKRPSWTKEEISSMKPKEFTISPDCGTIRPQGFAAIRVTLCSNTVQKYELALVVDVEGIGEEVLALLIAARCVVPALHLVNTEVDFGHCFLKYPYEKTLQLADQDDLPGFYEVQPQVCEEVPTVLFSSPTPSGVISPSSTIHIPLVLETQVTGEHRSTVYISIFGSQDPPLVCHLKSAGEGPVIYVHPNQVDFGNIYVLKDSSRILNLCNQSFIPAFFQAHMAHKKSLWTIEPNEGMVPPETDVQLALTANLNDTLTFKDCVILDIENSSTYRIPVQASGTGSTIVSDKPFAPELNLGAHFSLDTHYYHFKLINKGRRIQQLFWMNDSFRPQAKLSKKGRVKKGHAHVQPQPSGSQEPRDPQSPVFHLHPASMELYPGQAIDVILEGYSATPRKPNSILKPDYQPLAVKNISTLPVNLLLSTSGPFFICETDKSLLPATPEPIKLEIDEEKNLLIKFDPSYRNDLNNWVAEEILAIKYVEHPQIDSLDLRGEVHYPNLSFETKELDFGCILNDTELIRYVTITNCSPLVVKFRWFFLVNDEENQIRFVTLPKKPYSAPLSQMESIPATSEAASPPAILVTVESPEMDLNDFVKK** |
| PI4KAP1 | **16** | **0.69** | **GXXATGc** | **135** | **Yes** | | **MLALQIIDLFKNIFQLVGLDLFVFPYRVVATAPGCGVIECIPDCTSRDQLGRQTDFGMYDYFTRQYGDESTLAFQQACYNFIRSMAAYSLLLFLLQIKDRHNGNIMLDKKGHIIHIGQPATAPPSSPFTPAPRGG** |

| | | | | | | | |
|---|---|---|---|---|---|---|---|
| | **3** | **0.33** | **cXXATGa** | **262** | **Yes** | **Yes** | **MTVEQKFGLFSAEIKEADPLAASEASQPKPCP<br>PEVTPHYIWIDFLVQRFEIAKYCSSDQVEIFSSLL<br>QRSMSLNIGGAKGSMNRHVAAIGPRFKLLTLG<br>LSLLHADVVPNATIRNVLREKIYSTAFDYFSCPP<br>KFPTQGEKRLREDISIMIKFWTAMFSDKKYLTA<br>SQLVPPADIGDLREQLVEENTGSLSGPAKDFY<br>QREFDFFNKITNVSAVIKPYPKGDQRKKACLSA<br>LSEVTVQPGCSLPSNPEAIVLDVDYKSGTPM** |
| | **18** | **0.27** | **GXXATGa** | **103** | **Yes** | | **MSPLWPSSRPATTSSEAWPPTASCCSCCRSR<br>TDTTATLCWTRRAISSTSVSQPQRHPPLPSPR<br>HPGVDRDPHPTERRMPRTTLPGVSGSSSEVRT<br>VGHQAVLL** |
| | 5 | 0.26 | AXXATGa | 181 | Yes | | MNRHVAAIGPRFKLLTLGLSLLHADVVPNATIRN<br>VLREKIYSTAFDYFSCPPKFPTQGEKRLREDISI<br>MIKFWTAMFSDKKYLTASQLVPPADIGDLREQL<br>VEENTGSLSGPAKDFYQREFDFFNKITNVSAVIK<br>PYPKGDQRKKACLSALSEVTVQPGCSLPSNPE<br>AIVLDVDYKSGTPM |
| | 11 | 0.26 | AXXATGa | 113 | Yes | | MIKFWTAMFSDKKYLTASQLVPPADIGDLREQL<br>VEENTGSLSGPAKDFYQREFDFFNKITNVSAVIK<br>PYPKGDQRKKACLSALSEVTVQPGCSLPSNPE<br>AIVLDVDYKSGTPM |
| PI4KAP2 | 18 | 0.44 | GXXATGt | 119 | Yes | | MYDYFTRQYGDESTLAFQQARYNFIRSMAAYSL<br>LLFLLQIKDRHNGNIMLDKKGHIIHIDFGFMFESS<br>PGGNLGWEPDIKLTDEMVMIMGGKMEATPFKW<br>FMEMCVQATWLCGTGLAPT |
| | **5** | **0.4** | **cXXATGa** | **501** | **Yes** | **Yes** | **MTVEQKFGLFSAEIKEADPLAASEASQPKPCP<br>PEVTPHYIWIDFLVQRFEIAKYCSSDQVEIFSSLL<br>QRSMSLNIGRAKGSMNRHVAAIGPRFKLLTLG<br>LSLLHADVVPNATIRNVLREKIYSTAFDYFSCPP<br>KFPTQGEKRLREDISIMIKFWTAMFSDKKYLTA<br>SQLVPPADIGDLLEQLVEENTGSLSGPAKDFY<br>QREFDFFNKITNVSAVIKPYPKGDERKKACLSA<br>LSEVTVQPGCSLPSNPEAIVLDVDYKSGTPMQ<br>SAAKAPYLAKFKVKRCGVSELEKEGLRCRSDS<br>EDECSTQEADGQKISWQAAIFKLGDDCRQDML<br>ALQIIDLFKNIFQLVGLDLFVFPYRVVATAPGCG<br>VIECIPDCTSRDQLGRQTDFGMYDYFTRQYGD<br>ESTLAFQQARYNFIRSMAAYSLLLFLLQIKDRH<br>NGNIMLDKKGHIIHIDFGFMFESSPGGNLGWEP<br>DIKLTDEMVMIMGGKMEATPFKWFMEMCVQA<br>TWLCGTGLAPT** |
| | 17 | 0.39 | GXXATGc | 176 | Yes | | MLALQIIDLFKNIFQLVGLDLFVFPYRVVATAPGC<br>GVIECIPDCTSRDQLGRQTDFGMYDYFTRQYG<br>DESTLAFQQARYNFIRSMAAYSLLLFLLQIKDRH<br>NGNIMLDKKGHIIHIDFGFMFESSPGGNLGWEP<br>DIKLTDEMVMIMGGKMEATPFKWFMEMCVQAT<br>WLCGTGLAPT |
| | 7 | 0.31 | AXXATGa | 420 | Yes | | MNRHVAAIGPRFKLLTLGLSLLHADVVPNATIRN<br>VLREKIYSTAFDYFSCPPKFPTQGEKRLREDISI<br>MIKFWTAMFSDKKYLTASQLVPPADIGDLLEQL<br>VEENTGSLSGPAKDFYQREFDFFNKITNVSAVIK<br>PYPKGDERKKACLSALSEVTVQPGCSLPSNPEA<br>IVLDVDYKSGTPMQSAAKAPYLAKFKVKRCGVS<br>ELEKEGLRCRSDSEDECSTQEADGQKISWQAAI<br>FKLGDDCRQDMLALQIIDLFKNIFQLVGLDLFVF<br>PYRVVATAPGCGVIECIPDCTSRDQLGRQTDFG<br>MYDYFTRQYGDESTLAFQQARYNFIRSMAAYSL<br>LLFLLQIKDRHNGNIMLDKKGHIIHIDFGFMFESS<br>PGGNLGWEPDIKLTDEMVMIMGGKMEATPFKW<br>FMEMCVQATWLCGTGLAPT |
| | 12 | 0.3 | AXXATGa | 352 | Yes | | MIKFWTAMFSDKKYLTASQLVPPADIGDLLEQL<br>VEENTGSLSGPAKDFYQREFDFFNKITNVSAVIK<br>PYPKGDERKKACLSALSEVTVQPGCSLPSNPEA<br>IVLDVDYKSGTPMQSAAKAPYLAKFKVKRCGVS<br>ELEKEGLRCRSDSEDECSTQEADGQKISWQAAI<br>FKLGDDCRQDMLALQIIDLFKNIFQLVGLDLFVF<br>PYRVVATAPGCGVIECIPDCTSRDQLGRQTDFG<br>MYDYFTRQYGDESTLAFQQARYNFIRSMAAYSL<br>LLFLLQIKDRHNGNIMLDKKGHIIHIDFGFMFESS<br>PGGNLGWEPDIKLTDEMVMIMGGKMEATPFKW<br>FMEMCVQATWLCGTGLAPT |

**Supplementary Table 4: Chimeric φgene ORF-associated protein structure-function prediction**

Table 2:Coding potential prediction for chimeric and single parent φgenes

| ID | Label | Coding probablility | Peptide length |
|---|---|---|---|
| HYDIN2 | CODING | 1 | 961 |
| ENSG00000259798 | NON-CODING | 0.0255482 | 51 |
| HYDINP1 | NON-CODING | 0.0357925 | 61 |
| ENSG00000276298 | NON-CODING | 0.0232834 | 38 |
| ANAPC1P1 | CODING | 1 | 455 |
| ANAPC1P2 | CODING | 1 | 369 |
| ANAPC1P3 | NON-CODING | 0.0595955 | 97 |
| ANAPC1P4 | CODING | 0.999106 | 634 |
| ANAPC1P5 | NON-CODING | 0.216475 | 185 |
| ANAPC1P6 | CODING | 0.990617 | 202 |
| PI4KAP1 | CODING | 0.99 | 263 |
| PI4KAP2 | CODING | 1 | 502 |
| GTF2IP1 | CODING | 0.99 | 405 |
| GTF2IP2 | NON-CODING | 0.208 | 83 |
| GTF2IP3 | NON-CODING | 0.05 | 104 |
| GTF2IP4 | CODING | 1 | 531 |
| GTF2IP5 | NON-CODING | 0.17 | 101 |
| GTF2IP6 | NON-CODING | 0.3 | 92 |
| GTF2IP7 | NON-CODING | 0.06 | 39 |
| GTF2IP8 | NON-CODING | 0.09 | 53 |
| GTF2IP9 | NON-CODING | 0.01 | 21 |
| GTF2IP11 | NON-CODING | 0.02 | 23 |
| GTF2IP12 | NON-CODING | 0.1 | 86 |
| GTF2IP13 | NON-CODING | 0.06 | 51 |
| GTF2IP14 | NON-CODING | 0.12 | 88 |
| GTF2IP18 | NON-CODING | 0.075 | 64 |
| GTF2IP20 | NON-CODING | 0.114 | 89 |
| GTF2IP22 | NON-CODING | 0.022 | 30 |
| GTF2IP23 | NON-CODING | 0.024 | 22 |
| ENSG00000165121 | NON-CODING | 0.251164 | 214 |
| ENSG00000234424 | CODING | 0.65448 | 132 |
| ENSG00000290924 | CODING | 0.999496 | 230 |
| LOC389765 | CODING | 0.631056 | 112 |

**Supplementary Table 4: Chimeric φgene ORF-associated protein structure-function prediction**

Table 3: Chimeric φgene ORF proteins structure and function information

| Parent protein | Length (AA) | Chimeric protein | Length (AA) | Sequence homology w.r.t. parent | | Structure comparison with parent (RMSD) | DeepLoc2.1 (writeup P=>0.5) | InterPro and ScanProsite |
|---|---|---|---|---|---|---|---|---|
| | | | | Identity (%) | Query cover (%) | | | |
| **ANAPC1** | 1944 | **ANAPC1P2** | 368 | 99 | 99 | 0.518 (highly similar) | Soluble protein | Anaphase-promoting complex subunit 1 (Apc1) |
| **HYDIN** | 5121 | **HYDIN2_1** | 787 | 99.49 | 100 | 5.513 (highly dissimilar) | Soluble protein | Hydin-like; Cep192 is a centrosomal protein; Hydin adenylate kinase-like domain; Immunoglobulin-like fold |
| | | | | | | | Cytoplasm localization | |
| | | **HYDIN2_2** | 635 | 80.75 | 97 | 2.523 (dissimilar) | Soluble protein | Hydin-like |
| | | | | | | | Cytoplasm localization | |
| **GTF2I** | 998 | **GTF2IP1** | 404 | 99.62 | 100 | 0.642 (highly similar) | Soluble protein | GTF2-like repeats |
| | | | | | | | Nucleus localization | |
| | | **GTF2IP4** | 530 | 100 | 98 | 3.885 (highly dissimilar) | Soluble protein | |
| | | | | | | | Nucleus localization | |
| **PI4KA** | 2102 | **PI4KAP2** | 592 | 90.21 | 99 | 1.115 (similar) | Peripheral | Phosphatidylinositol kinase |
| | | | | | | | Cytoplasm localization | |
| **KIF27** | 1398 | **ENSG00000290924** | 129 | 96.15 | 42 | 0.807 (highly similar) | Soluble protein | Intrinsically disordered region (81-100 AA) |
| | | | | | | | Cytoplasm localization | |
