## Supplementary figures for "Evolutionary and functional dynamics of chimeric pseudogenes (φgenes)"

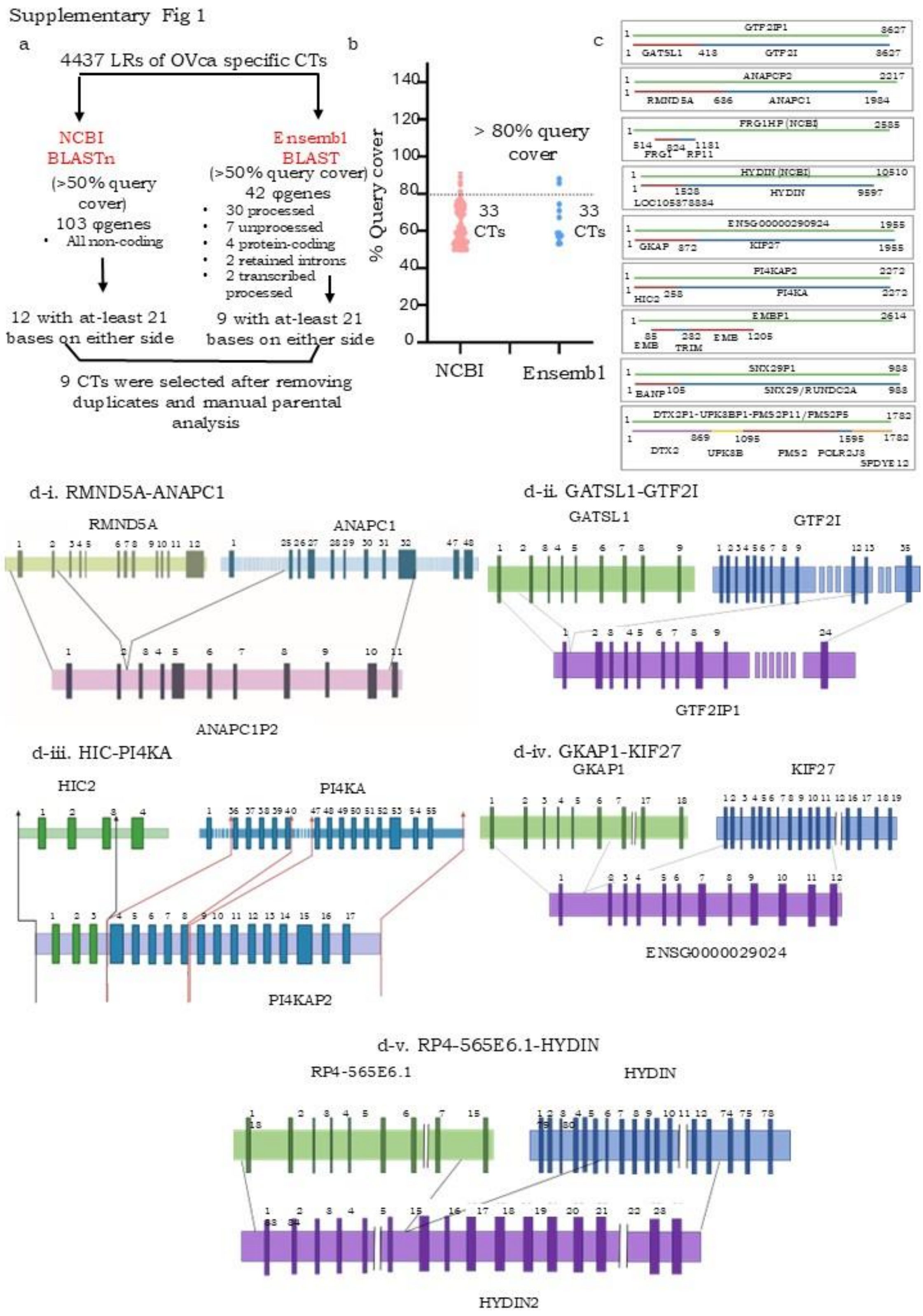


**Supplementary Fig. 1. Sequence similarity analysis of φgenes with their respective parental genes. a. Workflow for identifying chimeric transcripts (CTs) annotated as φgenes; b. Query cover distribution of CTs relative to predicted φgenes; c. Schematic depicting sequence homology of validated candidate CTs with predicted φgenes (green, red and blue lines represent φgene, 5' parental gene of CT and 3' parental gene of CT); d. Schematic representing the 5 simple intron fusions at complete gene level.**

Supplementary Fig.2

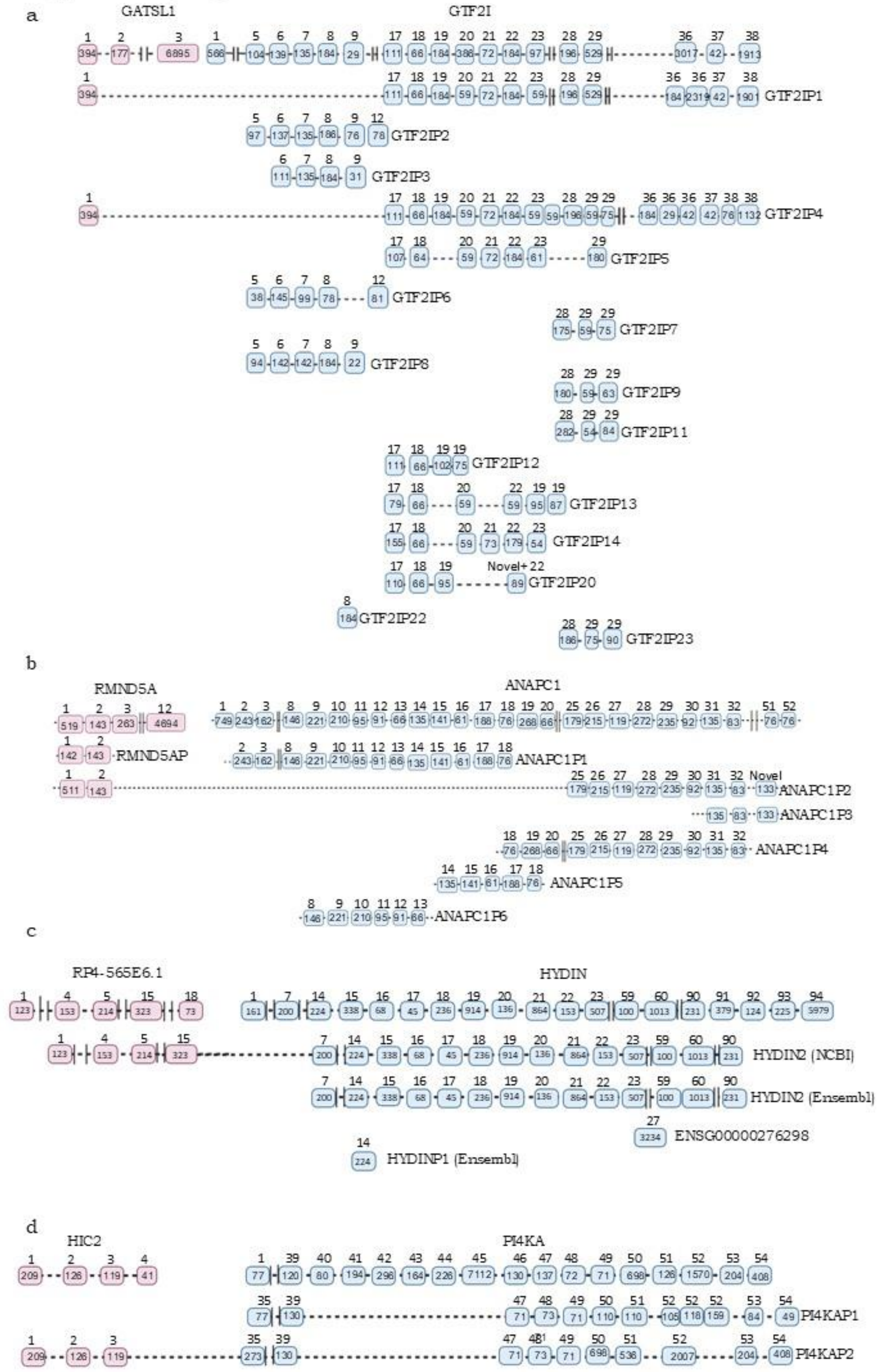


**Supplementary Fig. 2. Segmental duplications may be involved in the generation of chimeric φgenes. a-d. Schematic representations of genomic sequences of chimeric φgenes (*GTF2IP1*, *ANAPC1P2*, *HYDIN2*, *PI4KAP2*) as compared with their parent sequences**.

Supplementary Fig.3

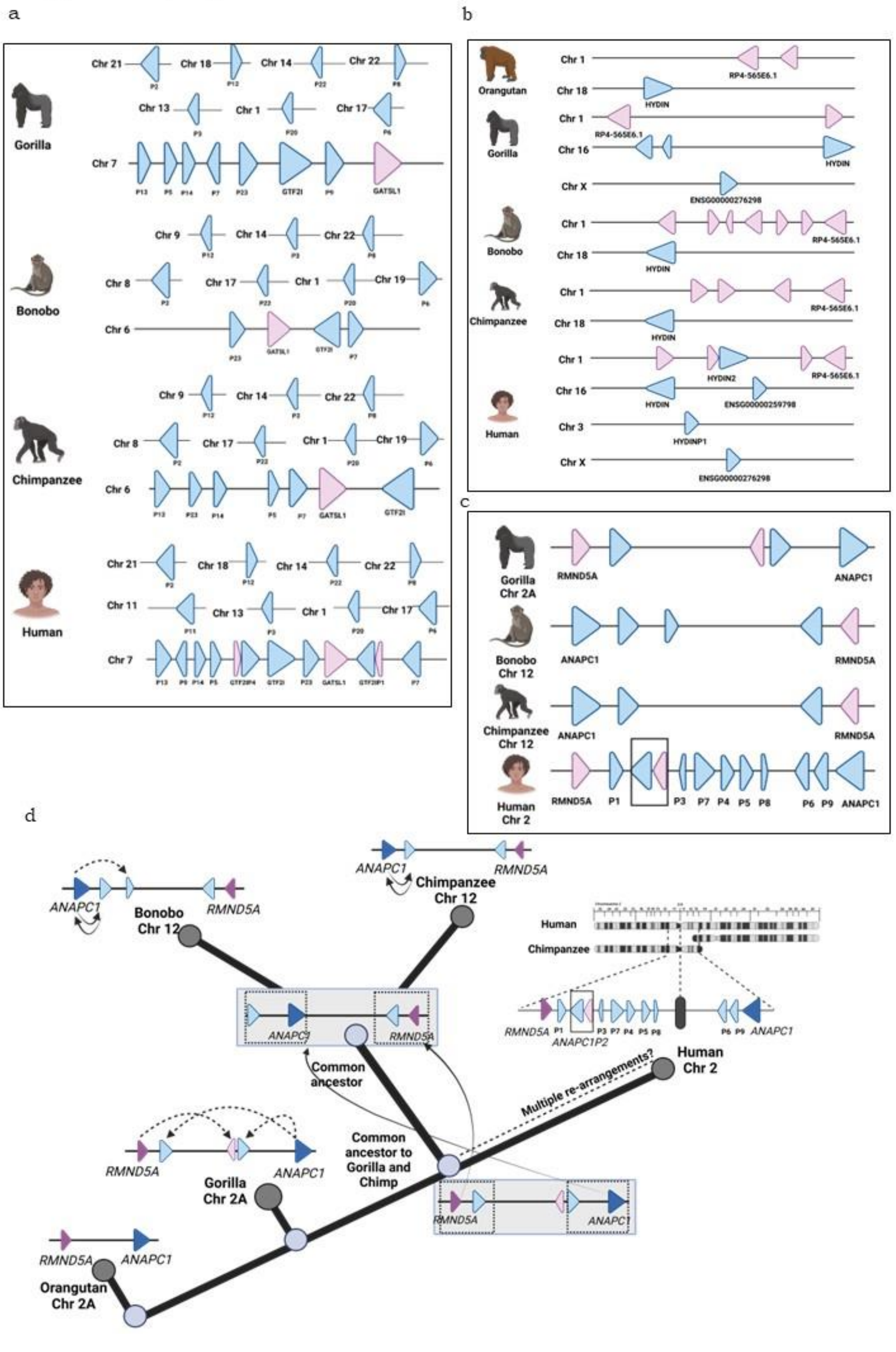


**Supplementary Fig. 3. Parental gene duplications leading to formation of single-gene and chimeric φgenes in hominoids (Orangutan, Gorilla, Bonobo, Chimpanzee and Human). a-c. Segmental duplication events and φgene formation across hominoids for *GTF2I*, *HYDIN2*, and *ANAPC1P2* respectively; d. Representative schematic of *ANAPC1P2* evolution across hominoid species through structural rearrangements involving multiple duplications of parental gene fragments (*RMND5A* and *ANAPC1*) and the predicted fusion event on chromosome 2.**

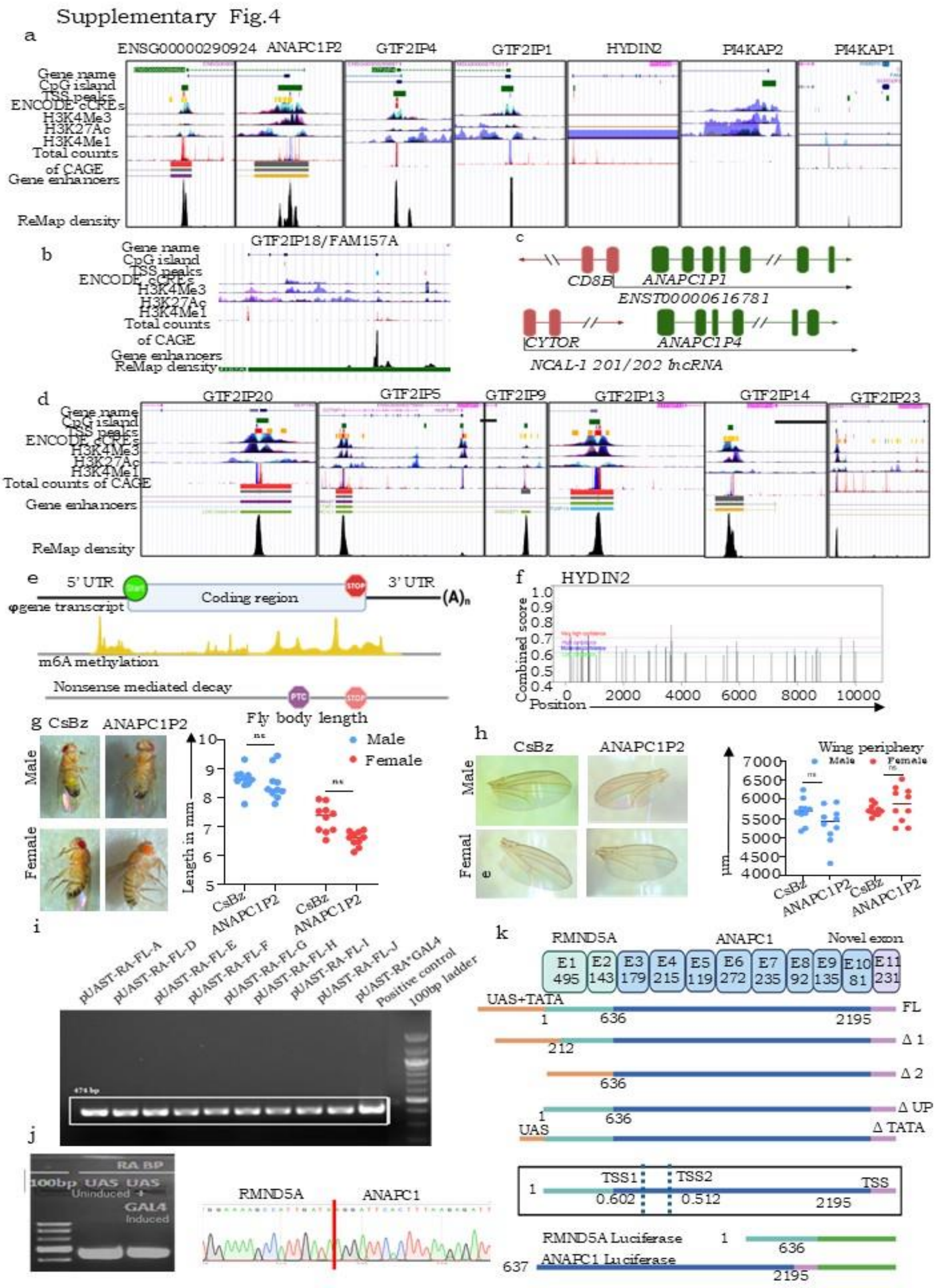


**Supplementary Fig. 4. 5' and 3' regulatory elements in single parent and chimeric φgenes. φgenes rely on promoter-like sequences in their neighbourhood to drive expression after duplication; a. Schematic of regulatory elements within chimeric φgenes from UCSC browser; b. 5' regulatory marks in *GTF2IP18*; c. *ANAPC1P1* and *ANAPC1P4* with promoters shared with upstream genes; d. GTF2I parental φgenes with promoters shared with upstream genes; e. 3' regulatory elements in the φgenes; f. m6A methylation marks in *HYDIN2* φgene; g. Expression of *ANAPC1P2* in cDNA of pUAST-ANAPC1P2 expressing flies; h. Expression of *ANAPC1P2* following transfection in S2 cells (with and without GAL4 induction) and Sanger sequencing; i. Schematic representing predicted transcription start site (TSS) and deletion/insertion variants generated for the study; j. Representative images of male and female *Drosophila melanogaster* from *ANAPCP2*-expressing and CsBz vector control groups (n = 10 per group), quantification of body length measurements for the corresponding groups; k. Representative images of individual fly wings from *ANAPC1P2*-expressing and CsBz control groups (n = 10 per group), quantitative analysis of wing perimeter for the respective experimental groups.**

Supplementary Fig. 5

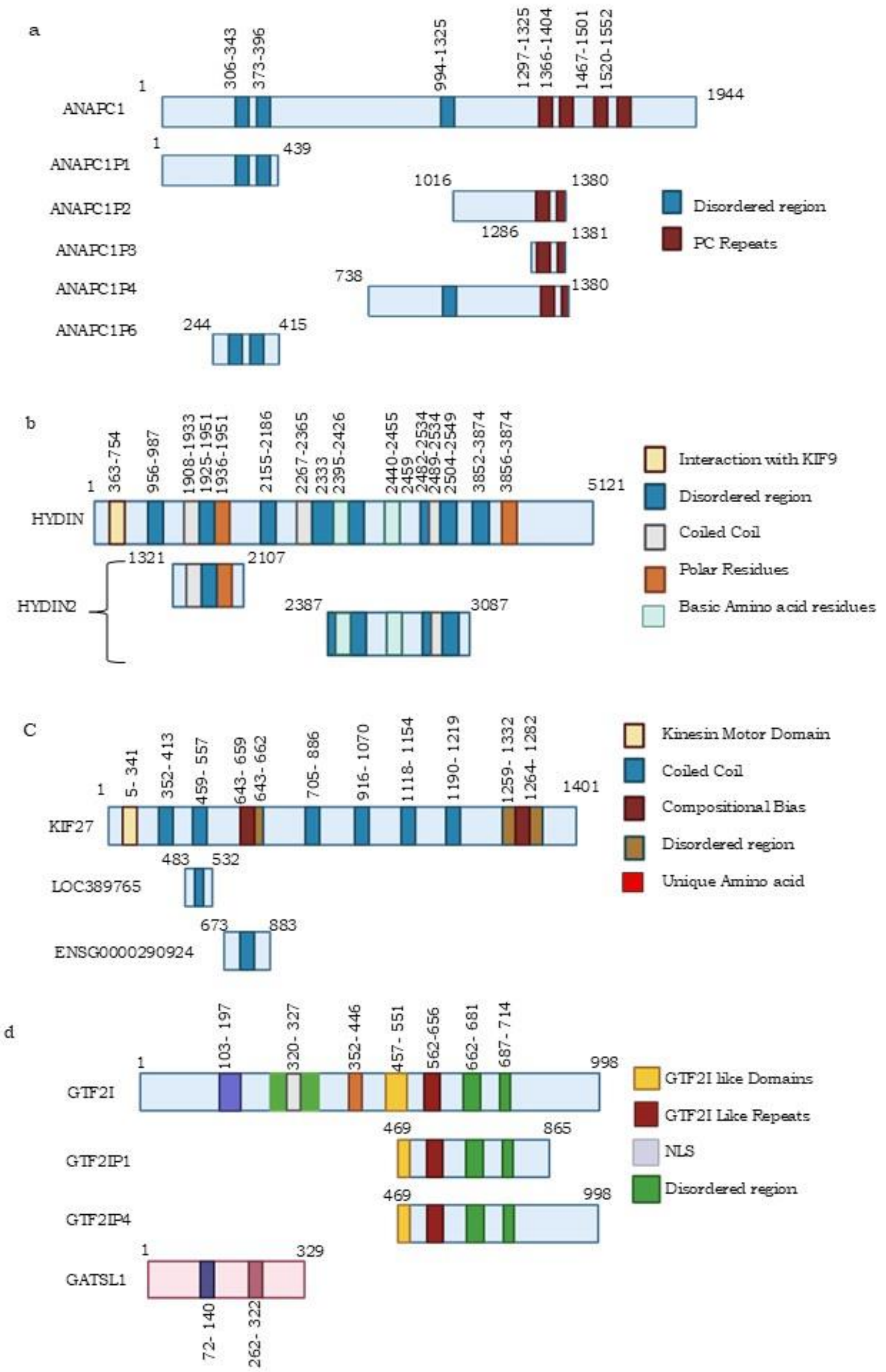


**Supplementary Fig. 5. Schematic representation of comparison between parental protein sequences with corresponding φgenes indicating the putative protein domains of various φgenes with reference to the parental proteins.; a. ANAPC1; b. HYDIN; c. KIF27; d. GTF2I.**

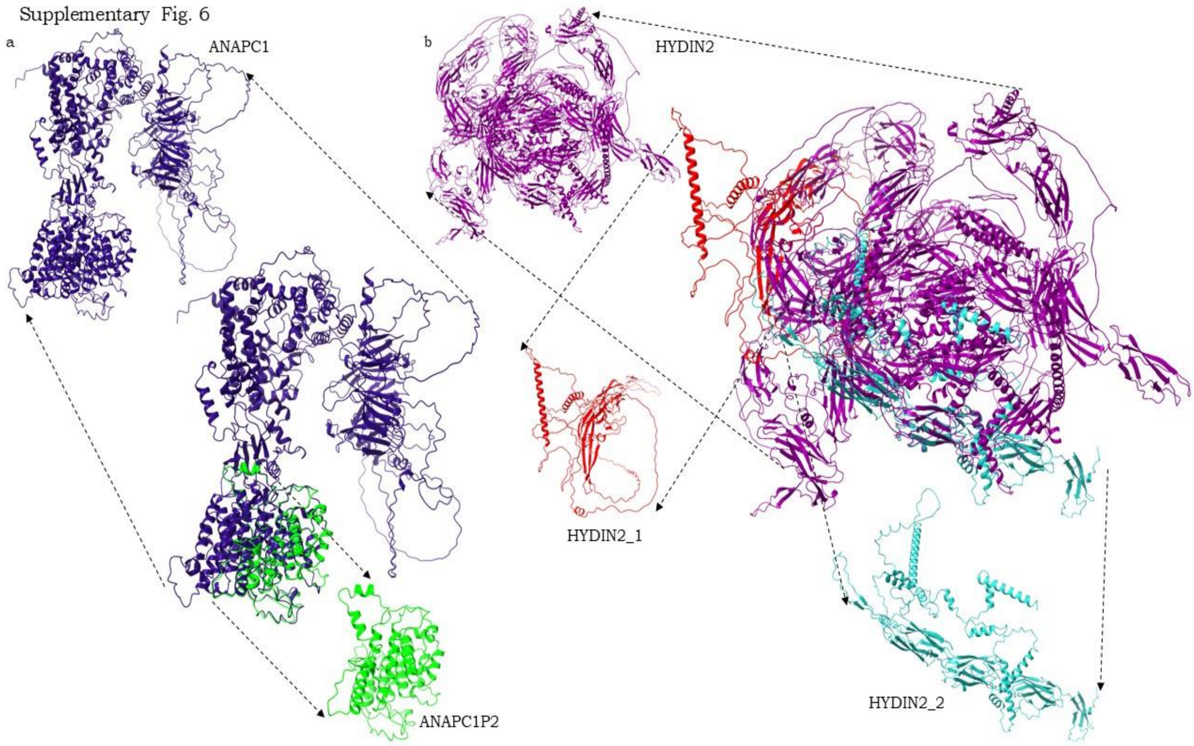


**Supplementary Fig. 6. Superimposed structures of chimeric φgene-derived proteins along with their parental counterparts predicted from the AlphaFold3 Server for; a. ANAPC1P2 (parental ANAPC1); b. HYDIN2_1 and HYDIN2_2 (parental HYDIN).**

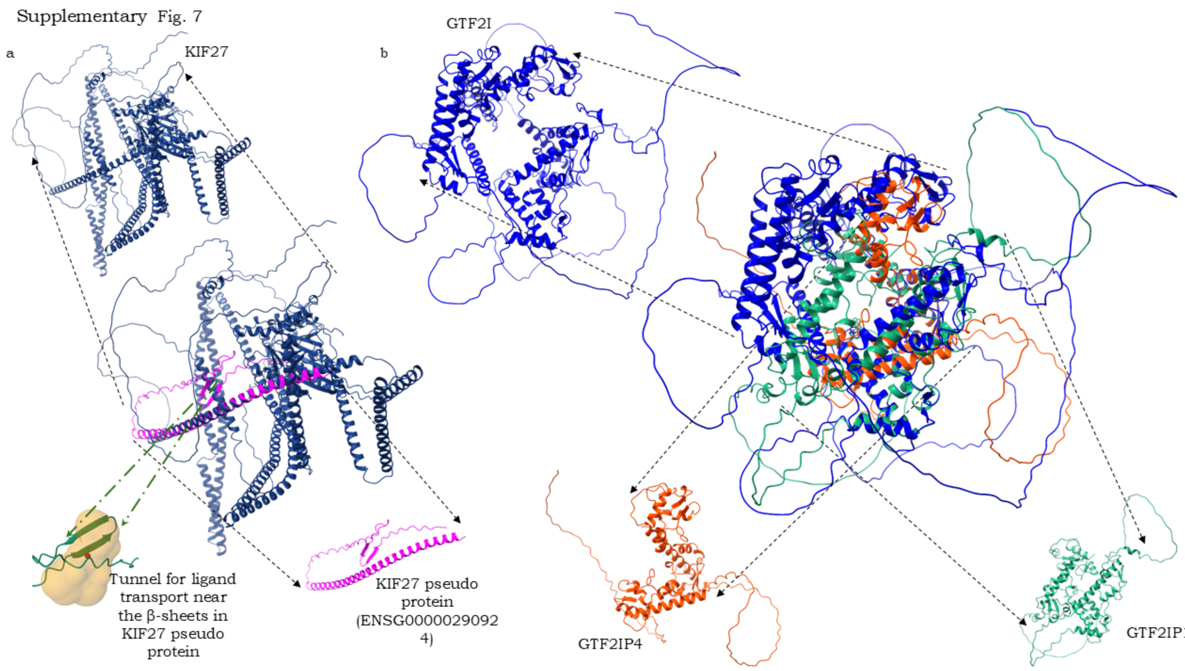


**Supplementary Fig. 7. Superimposed structures of chimeric φgene-derived proteins along with their parental counterparts predicted from the AlphaFold3 Server for a. ENSG000000290924 (parental KIF27); b. GTF2IP1 and GTF2IP4 (parental GTF2I).**

Supplementary Fig.8

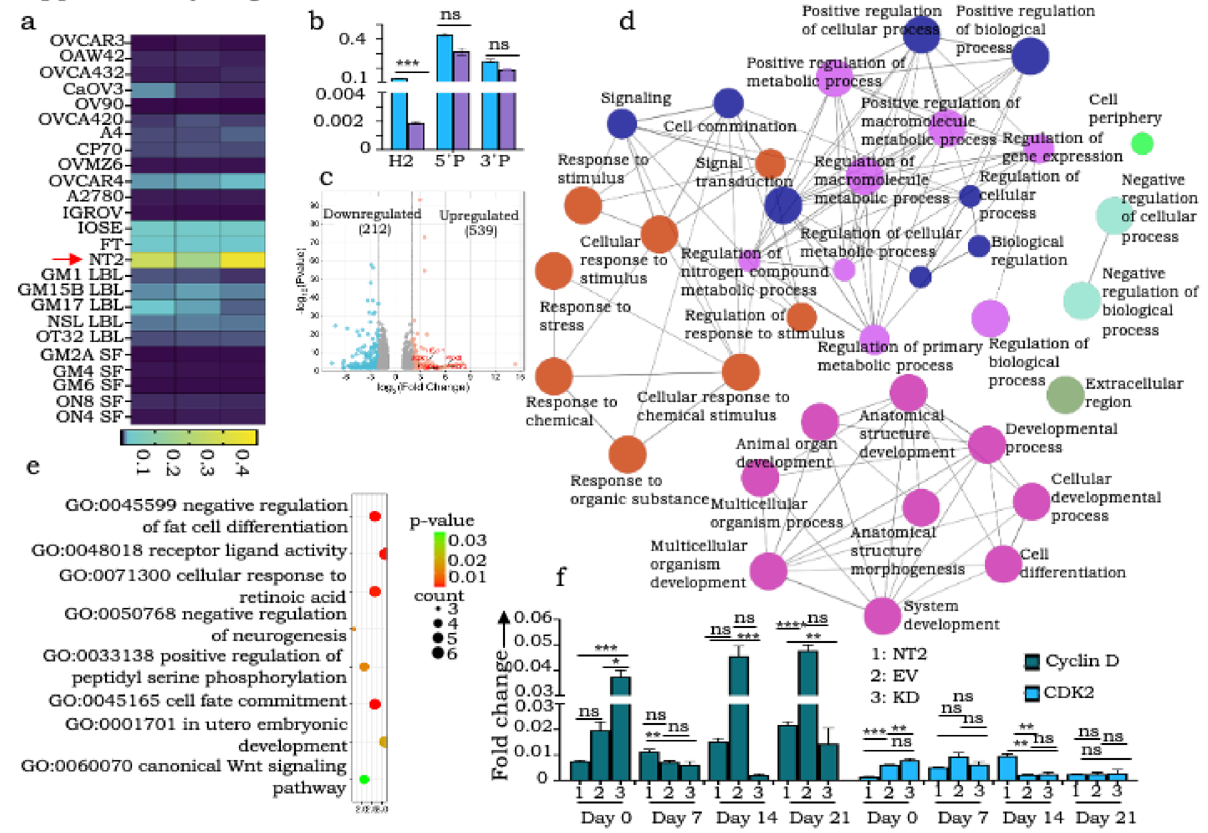


**Supplementary Fig.8. Human specific HYDIN2 knockdown alters neuronal differentiation. a. mRNA profiling of HYDIN2 across high grade serious ovarian cancer (HGSC), lymphoblastoid (LBL), skin fibroblast (SF), human normal (IOSE364, FT282) cell lines ; b. Expression profiling of RP4-565E6.1 (5'P), HYDIN2 (H2) and HYDIN (3'P) in HYDIN2 KD generated clones; c. Volcano plot showing differentially upregulated and downregulated pathways in HYDIN2 KD; d. Cluego networks showing differentially expressed pathways in HYDIN2 KD; e. Bubble plot representing differentially enriched pathways in HYDIN2 KD; f. Cyclin E and CDK2 transcript profiling in RA induced differentiated sample at various time points.**

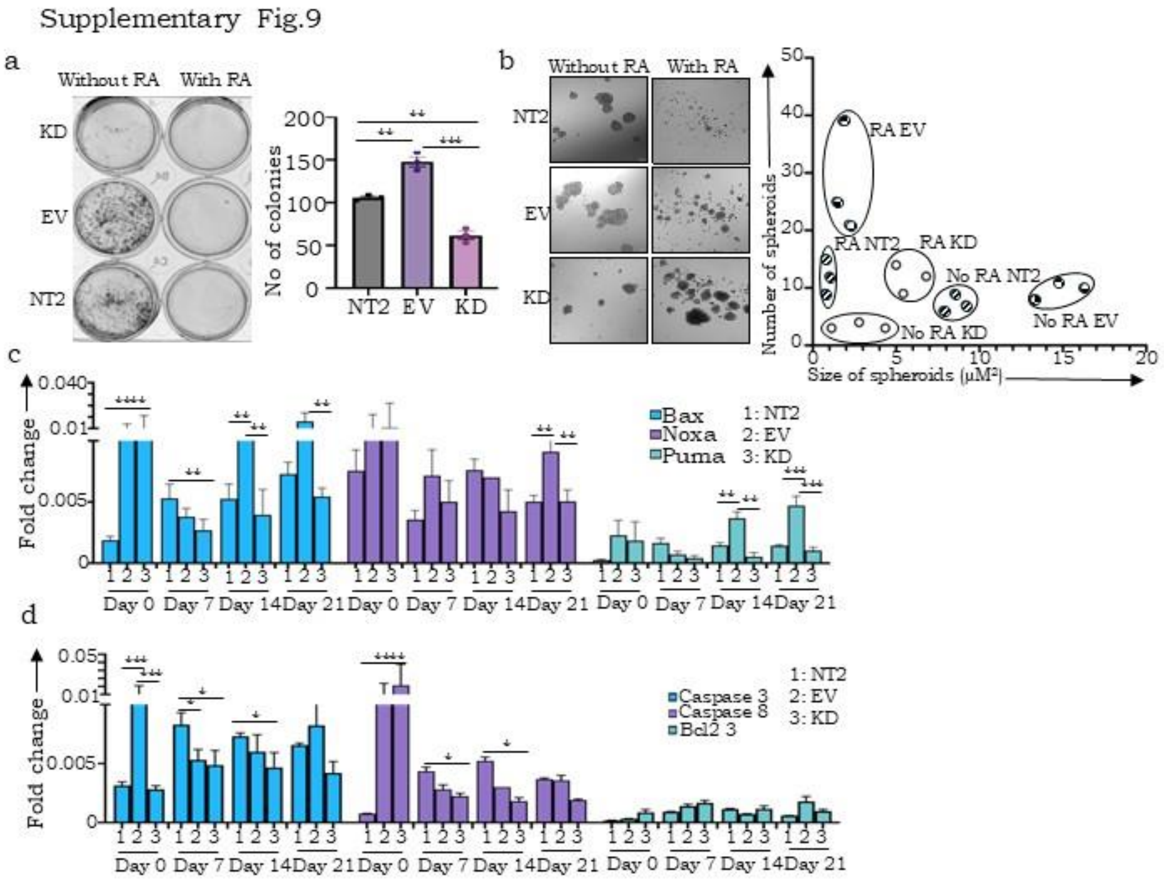


**Supplementary Fig.9. a. Representative images and quantification of spheroid with/ without RA induction; b. Representative images and quantification of number of colonies in all cell derivatives with and without RA induction; c-d. Expression profiling of Bax, Puma, Noxa, Caspase-3, Caspase-8, and BCI-2 in all cell derivates with and without RA induction till 21 days.**
